# Integrated multiomic profiling reveals SWI/SNF subunit-specific pathway alterations and targetable vulnerabilities

**DOI:** 10.1101/2024.07.16.603530

**Authors:** Jorge Bretones Santamarina, Clémence Astier, Marlène Garrido, Leo Colmet Daage, Theodoros I. Roumeliotis, Elodie Anthony, Mercedes Pardo, Marianne Chasseriaud, Pierre Gestraud, Carine Ngo, Daphné Morel, Roman Chabanon, Jyoti Choudhary, Elaine Del Nery, Sophie Postel-Vinay, Annabelle Ballesta

## Abstract

Mutations in subunits of the SWItch Sucrose Non-Fermentable (SWI/SNF) chromatin remodeling complex occur in ≈20% of cancers and represent a highly unmet medical need. To identify novel therapeutic approaches, we systematically characterized transcriptomic and proteomic changes caused by the loss of SWI/SNF subunits or other epigenetic enzymes in isogenic cell lines, which we subsequently integrated with high-throughput drug screening and independent genetic screens of the DepMap project. Using an optimized bioinformatics pipeline for pathway enrichment, we identified *Metabolism of proteins* as the most frequently dysregulated Reactome pathway category in SWI/SNF-defective cell lines. Drug screening and multiomic integration revealed multiple chemicals selectively cytotoxic for SWI/SNF-defective models, including CBP/EP300 or mitochondrial respiration inhibitors. A novel algorithm for the analysis of DepMap CRISPR screens independently identified synthetic lethality between SWI/SNF defects and *EP300* or mitochondrial respiration genes, which we further revalidated in disease-relevant models. These results unravel novel genetic dependencies for SWI/SNF-defective cancers.

## INTRODUCTION

Epigenetic dysregulation has recently been identified as a major cancer hallmark^1^. Notably, deleterious mutations in genes encoding subunits of SWI/SNF (SWItch Sucrose Non-Fermentable), a major chromatin remodeling complex, occur in approximately 20% of human solid tumors^2,3^. They have been linked to poor patient outcomes and represent an unmet medical need^4^. SWI/SNF orchestrates multiple functions in cellular physiology, such as cell differentiation, proliferation, adhesion, chromosome segregation, DNA repair, gene expression, and immunogenicity^5^. Previous studies have shed light on the mechanisms by which mutations in certain SWI/SNF subunits contribute to tumor development^6^, but SWI/SNF subunit-specific oncogenic mechanisms are still far from being unveiled. SWI/SNF is a modular complex composed of 12-15 subunits encoded by 29 genes, which exists in three forms: canonical BRG1/BRM associated factor (cBAF), polybromo-associated BAF (PBAF) and non-canonical (ncBAF/GBAF), which have variable compositions, targets and effects on chromatin remodeling^7^. Each complex includes at least three core subunits (SMARCC1, SMARCC2, and SMARCD1-3) and one of the two mutually exclusive ATPase subunits (SMARCA2 or SMARCA4). Multiple variant subunits then define each complex’s specificity: ARID1A/B and DPF1-3 for cBAF; ARID2, PBRM1, BRD7 and PHF10 for PBAF; GLTSCR1/1L and BRD9 for ncBAF^2^. If this modular structure has recently been deciphered, the phenotypic effects and targetable dependencies that result from the loss of distinct subunits - which are mutated in different cellular contexts and cancer subtypes - are still poorly understood^8^.

Over the past years, novel therapeutic strategies have emerged to target vulnerabilities resulting from defects in SWI/SNF subunits^5,9^, notably based on synthetic lethality and epigenetic antagonism. Such approaches exploit a situation where the simultaneous loss (or inhibition) of two genes (or proteins) leads to cell death, while the loss of either one does not affect cell survival. In SWI/SNF, intra-complex dependencies operate notably between paralog subunits, e.g. *SMARCA4*- or *ARID1A*-mutant tumor cells’ survival depend respectively on SMARCA2 and ARID1B expression^6,10,11^. Extra-complex dependencies also operate with certain signaling pathways or mechanisms in which SWI/SNF is involved. For example, following the description of an epigenetic antagonism between SWI/SNF and EZH2 (Enhancer of Zest Homolog 2)^12,13^, the first EZH2 inhibitor tazemetostat was assessed in *SMARCB1*-deficient tumors. The observed efficacy (15% response rate) in epithelioid sarcoma justified its accelerated approval in 2020, thereby representing the first-ever approved epigenetic drug in solid tumors^12–15^. In October 2023, the 2^nd^ generation EZH1/2 inhibitor tulmimetostat received Fast Track Designation for the treatment of endometrial tumors with *ARID1A* mutations^16^. Other synthetic lethal interactions with SWI/SNF subunit defects have been described, e.g., between ATR or PARP inhibitors and defects in SMARCA4^17,18^ or ARID1A^19,20^, or between BRD9 inhibitors and SMARCB1 alterations^21,22^. Still, except for the examples mentioned above, none of them has reached clinical approval so far, and defects in most subunits are still undruggable, highlighting the urgent need for additional therapeutic approaches^5^.

Multi-omics profiling of cancer cell lines combined with high-throughput genetic, or drug screening allows to uncover protein network dysregulation and targetable vulnerabilities in a systematic fashion^23,24^. Since interpretation of such high-dimensional omics datasets may be challenging, pathway enrichment methodologies have been developed to inform on the dysregulation of gene sets corresponding to cellular functionalities. The choice of database providing pathway definitions may greatly affect the reliability of the results. Indeed, testing large numbers of pathways – which are each considered as an independent hypothesis test - harms the statistical performance of the method because of the need for multiple testing correction^25^ and may violate the assumption of statistical independence between tests, since pathways sharing genes are not independent^26^. Regarding enrichment algorithms, first-generation algorithms (e.g., Overrepresentation Analysis, ORA) assess whether a pathway is overrepresented in a list of differentially expressed genes, regardless of gene expression fold change (FC) between conditions. Second-generation algorithms, such as Gene Set Enrichment Analysis (GSEA), do account for quantitative gene FC information, whereas third-generation enrichment methods such as ROntoTools further include the network structure (topology) of the pathway^27^. A recent benchmark of these methods found that ROntoTools and GSEA were the best-performing methods, despite numerous false negatives, whereas ORA was less efficient with a high rate of false positives^28^. Thus, each of these algorithms has inherent limitations, which calls for further methodological development.

To unravel novel targetable vulnerabilities in SWI/SNF-defective cancers, we integrated in-house multi-omics profiling with high-throughput drug screening, using an isogenic panel of SWI/SNF subunit- and non-SWI/SNF chromatin remodeling-genes knock-out cell lines (herein referred as “SWI/SNF mutants” and “non-SWI/SNF mutants”). To improve pathway enrichment reliability, we developed an optimized gene set analysis pipeline, which allowed us to identify dysregulated networks in each condition. To assess the transferability of our results, we next developed a novel statistical algorithm to predict synthetic lethal interacting genes for each SWI/SNF subunit using genetic screens of the DepMap cancer cell line encyclopedia and compared hits to our *in-house* dataset. The combination of both studies allowed us to identify novel genetic vulnerabilities, notably linked to protein metabolism, mitochondrial respiration and CBP/EP300 inhibition, which we revalidated in independent SWI/SNF-deficient disease-relevant models.

## RESULTS

### Loss of individual SWI/SNF subunits variably alters the complex stoichiometry

To identify cell regulatory network alterations and genetic vulnerabilities induced by SWI/SNF defects, at the complex or subunit level, we molecularly and functionally profiled an isogenic panel of HAP1 cell lines, where the seven SWI/SNF subunits most frequently altered in cancer had been knocked-out (Fig. 1A; Supplementary Tables 1-2). This panel comprised mutants in the core subunit SMARCB1, catalytic subunits SMARCA4 and SMARCA2, cBAF-specific subunits ARID1A and ARID1B, and P-BAF-specific subunits ARID2 and PBRM1. In parallel, we similarly characterized HAP1-derived isogenic models where genes encoding other chromatin remodelers had been disabled, to be able to identify SWI/SNF-specific effects (Supplementary Tables 1-2). The latter panel comprised mutants of CREBBP, BAP1 and EED (Polycomb Repressive Complex), KMT2C and KMT2D (COMPASS), and SETD2.

**Figure 1.**
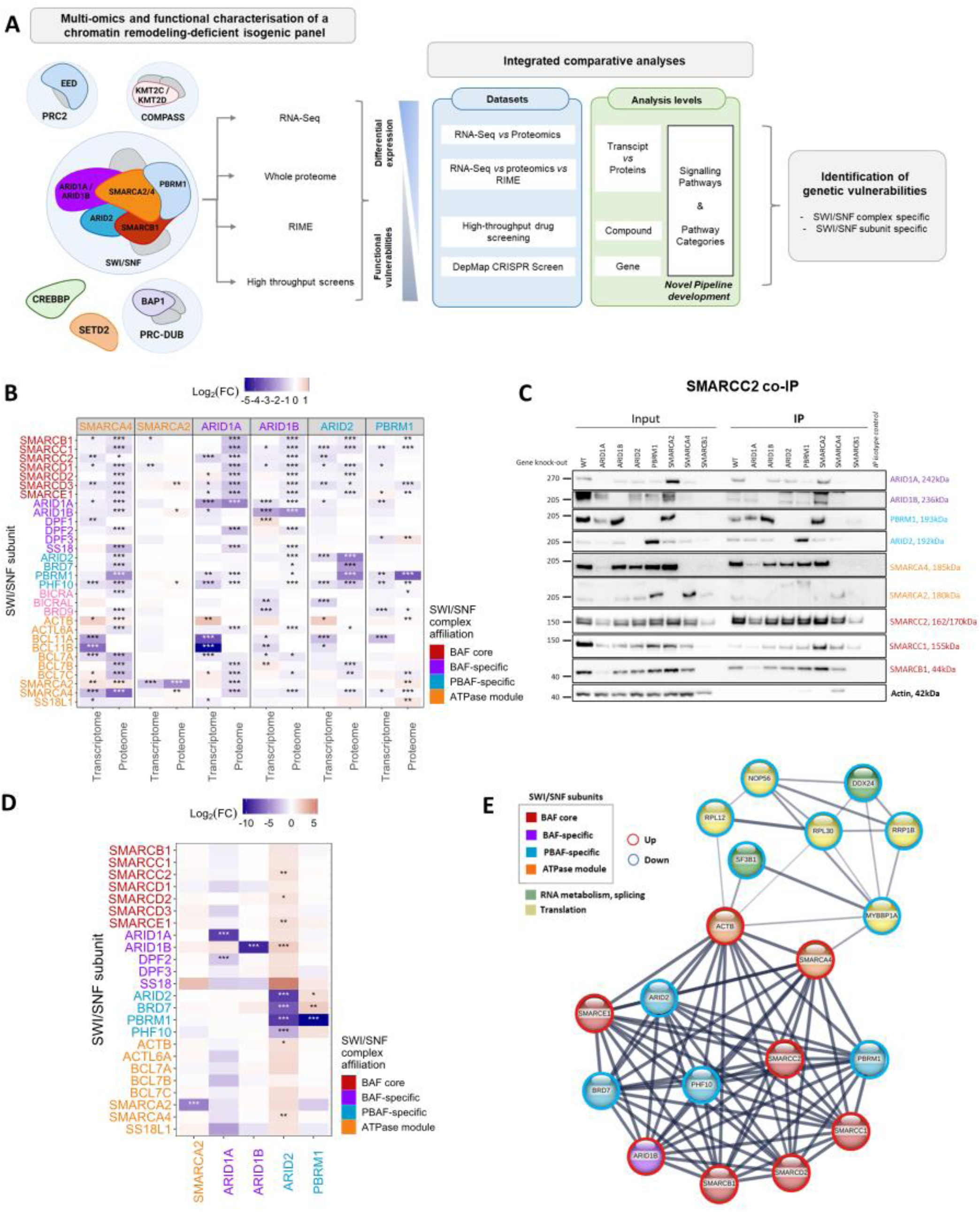
SWI/SNF subunit defects lead to subunit-specific transcriptomic, proteomic and stoichiometric changes in SWI/SNF complex composition. A) Study workflow. SWI/SNF subunits and other chromatin remodelers for which KO isogenic HAP1 mutants were profiled are shown on left panel; molecular and profiling methods, as well as analyses performed are depicted in middle and right panels. B) SWI/SNF related subunits differential expression between indicated SWI/SNF mutant *and* WT in transcriptome and proteome analysis. SWI/SNF mutants and SWI/SNF subunits are colored according to the type of SWI/SNF complex they belong to according to Mashtalir et al^8^. The color scale represents the log_2_(Fold-change) between gene expression in each mutant as compared to the wild-type cell lines. Positive values indicate gene/protein upregulation with respect to HAP1 wild-type, whereas negative values indicate downregulation. p-adj value: *** < 0.0001, ** < 0.001, * < 0.01. C) Western blot of selected SWI/SNF subunits following immunoprecipitation (IP) of the SMARCC2 core subunit. D) SWI/SNF subunits detected by quantitative RIME following SMARCC2 IP. See panel B for legend. E) String network of the ARID2-mutant RIME data showing the significant down-regulation of all PBAF-specific subunits (blue) and the upregulation of SS18 (red).

Using transcriptomics and proteomics data, we first verified the absence of expression of each subunit in its corresponding knock-out (KO) cell line (Supplementary Tables 1-2). SWI-SNF subunit KO resulted in modifications of other components of the complex, which were overall more pronounced at the protein than at the mRNA level, suggesting that these might, at least in part, result from altered protein-protein interactions (e.g. BRD7 or PBRM1 decrease in *ARID2*-KO implicating stabilising protein interactions; Fig. 1B). Since SWI/SNF subunits assemble in a modular fashion^8^, we next characterized the complex composition using immunoprecipitation of the SMARCC2 core subunit, present in cBAF, PBAF, and ncBAF. As previously described^29^, western blotting of selected subunits found that loss of SMARCB1 impaired the incorporation of almost all other BAF or PBAF-specific subunits, whereas loss of either SMARCA2 or SMARCA4 increased expression and incorporation of their respective paralog. In contrast with PBRM1 decrease observed in *ARID2*-KO, PBRM1 loss led to increased expression and incorporation of the PBAF-specific subunit ARID2, highlighting the importance of directionality in protein interactions (Fig. 1C)^30^. To quantitatively interrogate these intra-complex modifications, we performed quantitative Rapid Immunoprecipitation Mass spectrometry of Endogenous proteins (RIME) on SWI/SNF-WT and KO-models, using SMARCC2 immunoprecipitation (Fig. 1D-E, Supplementary Table 3). This revealed additional perturbations that were not previously identified at the transcriptomic or proteomic level. For example, loss of ARID2, beyond significantly decreasing the incorporation of PHF10, BRD7 and PBRM1, increased that of the cBAF subunit ARID1B (log_2_FC = 2.26, p-adj < 0.001) and some SWI/SNF core subunits (Fig. 1E). Loss of ARID2 also led to decreased interactions of SMARCC2 with a subset of proteins involved in RNA metabolism or splicing (e.g. DDX24, log_2_FC = -2.06; SF3B1, log_2_FC = -3.08; p-adj < 0.05), or translation (e.g., RRP1B, log_2_FC = -4.51; NOP56, log_2_FC = -3.61; MYBBP1A, log_2_FC = -2.26; all p-adj < 0.001), reminiscent of the recently described role of SWI/SNF in alternative splicing in mammals^31,32^, endoplasmic reticulum homeostasis in yeast^33^, and dependency of SWI/SNF-altered cells on translation factors^34^.

Altogether, this shows that loss of SWI/SNF subunits destabilizes SWI/SNF stoichiometry in subunit-specific manner and supports that cellular consequences of SWI/SNF subunit defects can both result from direct and indirect effects.

### Loss of individual SWI/SNF subunits distinctly alters gene and protein expression

If most SWI/SNF functions have traditionally been linked to transcription regulation, recent reports^31–34^ and our RIME results suggested that SWI/SNF defects might also alter RNA splicing or translation. We therefore hypothesized that some SWI/SNF-dependent signaling effects might occur uniquely at the protein level and investigated the effects of SWI/SNF subunit defects at both the transcriptome and proteome levels.

As a quality control, we first benchmarked our transcriptomic profiles with the ones of a previously published dataset of HAP-1 SWI/SNF mutants and found a positive moderate correlation across the six cell lines that were common to both datasets (Pearson correlation coefficient R=0.364 ± 0.182 (mean ± standard deviation (SD)), Supplementary Fig. 1)^7^. When performing hierarchical clustering of all chromatin remodeling mutants based on transcriptomic or proteomic data, we found that SWI/SNF-mutants clustered separately from non-SWI/SNF mutants, reinforcing the specificity of SWI/SNF functions (Fig. 2A-B). However, SWI/SNF mutants did not sub-cluster according to their functional domains (e.g., DNA binding, ATPase, etc.) or specificity towards cBAF or PBAF, perhaps because of compensation mechanisms that modulate the complex’s stoichiometry (Fig. 1). Wild-type vs KO differential analysis of transcriptomics data identified a mean number of 6117 (range 4058 to 7433) and 7098 (range 5653 to 8506) significantly dysregulated genes across SWI/SNF-KO and non-SWI/SNF-KO cell lines, respectively, with 1864 and 2153 genes having a |log_2_FC| > 1 and p-adj < 0.05 (range 1106 to 2625 and 1704 to 2849), respectively (Fig. 2C; Supplementary Table 1, see Methods). Differential analysis of proteomics data revealed a mean of 4170 (range 2147 to 5612) dysregulated proteins across SWI/SNF-KO, including a mean of 168 with a |log_2_FC| > 1 (range 32 to 388) (Fig. 2C; Supplementary Table 2). Interestingly, we noticed that the correlation of the proteome between paralogs was weaker than the one between independent subunits (e.g. Pearson’s correlation scores R=0.3255 ARID1A/B, R=0.13 SMARCA2/4 versus R=0.5 ARID1A/PBRM1 or R=0.66 ARID1A/SMARCA4), suggesting that the paralog’s functions are only partly redundant (Supplementary Fig. 2). In the non-SWI/SNF group, there were 3596 differentially expressed proteins on average (range 1043 to 4907), including 233 with a |log_2_FC| > 1 (range 112 to 404) (Fig. 2C; Supplementary Table 2). The most dysregulated genes and proteins were either shared between SWI/SNF-mutants (e.g., increase of *JAK3* mRNA level, decrease of SHISA2 or PGM5 protein expression; Fig. 2D-E) or mutant-specific (e.g., membrane glycoproteins LAMA1 or BACE2 upregulation in *PBRM1*-KO only; log_2_FC = 6.69 and 6.25 respectively, p-adj<0.001; Supplementary Tables 1-2). Interestingly, several sensors, effectors, or receptors of the cell-autonomous innate immune pathways (e.g., OAS2, cGAS, JAK3, IFITM1) were found significantly upregulated across several SWI/SNF-KO cell lines, notably *ARID1A*- and *PBRM1*-KO models, reminiscent of the pro-immunogenic consequences of the loss of these subunits^35–40^.

**Figure 2.**
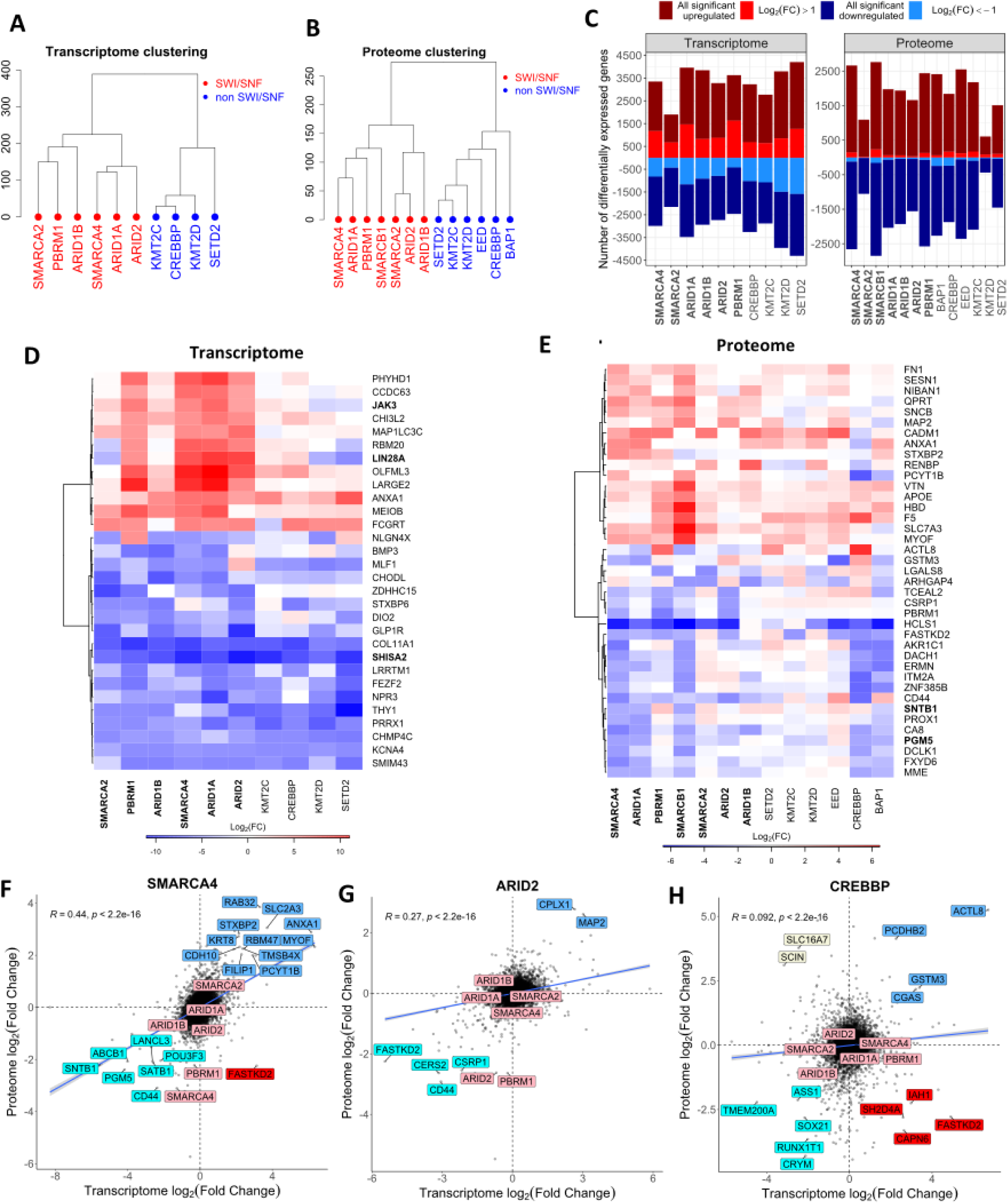
Differential expression analysis of whole transcriptome and proteome profiling shows that SWI/SNF mutants cluster together despite mutant-specific features and reveals a limited correlation between transcript and protein dysregulations. **A-B)** Hierarchical clustering of mutant cell lines according to whole transcriptome (A) and proteome (B) differential expression analysis. **C)** Number of differentially expressed genes and proteins for each mutant cell line as compared to WT. The significance level was set to p-adj < 0.05 for any log_2_FC, light colors flag entries with |log_2_FC| > 1. **D-E)** Most commonly dysregulated genes (D) and proteins (E) across SWI/SNF mutants (genes with |log_2_FC| > 4 and p-adj < 0.0001 in at least four SWI/SNF KO cell lines; proteins with |log_2_FC| > 2 and p-adj < 0.05 in at least two SWI/SNF KO cell lines)). Gene symbols in bold are mentioned in the text. **F-H)** Comparison of transcriptome and proteome differential expression in SMARCA4 (F), CREBBP (G) and ARID2 mutants (H). SWI/SNF-related genes are colored in pink and most differentially expressed genes (i.e. |log_2_FC| > 2) are highlighted as follows: blue, upregulation in both transcriptome and proteome; cyan, downregulation in both transcriptome and proteome; beige, downregulation in transcriptome and upregulation in proteome; red, upregulation in transcriptome and downregulation in proteome. The blue line represents the Pearson correlation line with indicated R coefficients and p values.

We next compared the transcriptomic and proteomic profiles and found a variable correlation across SWI/SNF-mutant cell lines, the highest and lowest ones being respectively observed for *SMARCA4*-KO and *ARID2*-KO cell lines (R = 0.44 and 0.27 respectively, p<0.001; across-mutant mean R = 0.38, Fig. 2F, G; Supplementary Fig. 2). Among all chromatin remodeling mutants, the lowest correlation was observed for the *CREBBP*-KO cell line (R = 0.092, p < 0.001; Fig 2H). Two distinct patterns could be identified: (i) changes that were consistent between RNA and protein (e.g. decreased *PGM5* or *SNTB1* expression in the *ARID1A*- and *SMARCA4*-KO cells); and (ii) changes that were inconsistent between RNA and protein levels. For example, HCLS1 was significantly decreased in all SWI/SNF-mutant cell lines at the proteome level only (log_2_FC = -3.77 to -5.73), possibly indicating post-translational regulatory mechanisms (Fig. 2E, Supplementary Tables 1-2). By contrast, the RNA binding protein *LIN28A,* which was experimentally shown to interact with SMARCA4 and SMARCB1^41^, was upregulated only at the transcriptomic level in almost all SWI/SNF mutants (log_2_FC = 4.66 – 8.38; Fig. 2D, Supplementary Tables 1-2).

Overall, this supports that transcriptomic profiling only partly recapitulates molecular changes induced by defects in SWI/SNF subunits, and that proteomic analysis brings a meaningful additional level of information.

### Development of an optimized pathway enrichment method combining GSEA and RontoTools

To go beyond the transcript or protein level and systematically explore functional consequences of SWI/SNF defects, we developed a new Gene Set Analysis strategy based on i) the design of a pruned version of Reactome pathway database with a reduced number of pathways and ii) the development of an optimal pipeline undertaking the most efficient combination of existing enrichment methods (Fig. 3A, Supplementary Fig. 3, Supplementary File 1). Reactome pathways are arranged as a tree where the 29 largest pathways are called the *categories* and are broken down into smaller pathways, which could be seen as sub-categories, which are themselves divided into smaller ones. Keeping all pathways of such a redundant database may affect enrichment performance in two ways: (i) violating the assumption of statistical independence between pathways, as pathways may share a certain amount of genes^25,26^ and (ii) increasing the multiple testing correction penalty by testing a large number of pathways. Moreover, testing many redundant pathways can compromise results’ interpretability. We therefore reasoned that using a pruned Reactome database may improve the performance of enrichment methods by reducing the number of tested gene sets and the gene redundancy across pathways, while exploring all cellular functions. To do so, we used a two-step approach (see Methods): (i) a top-down step, which eliminated pathways that were exceedingly large (> 500 genes), small (< 10 genes), or redundant (i.e. which could be subdivided into smaller child pathways of acceptable sizes), and (ii) a bottom-up step, which eliminated pathways included in larger ones fulfilling the size criteria.

**Figure 3.**
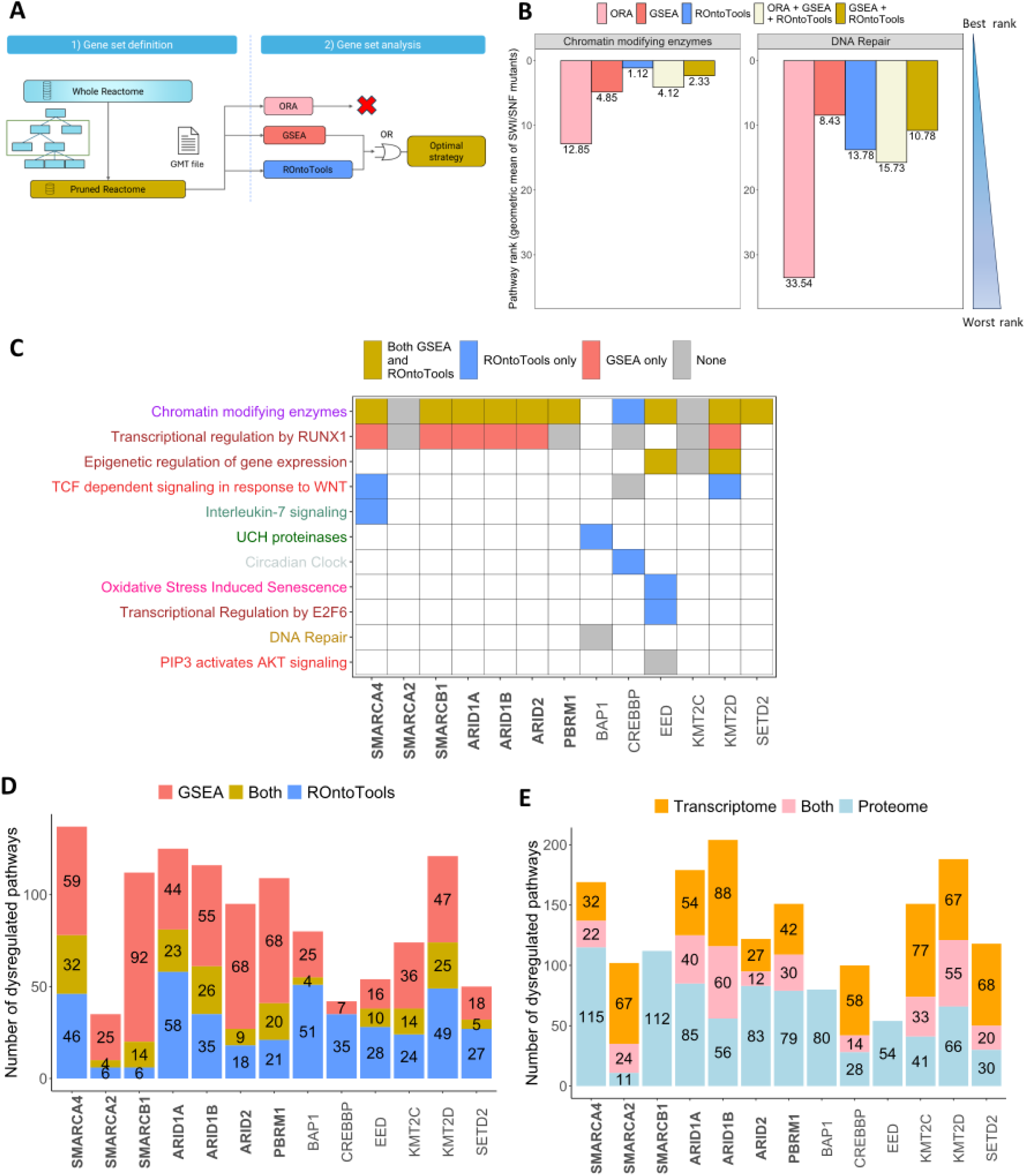
An optimized pathway enrichment pipeline based on pruned pathway database and combination of GSEA and RontoTools algorithms identifies common and mutant-specific dysregulated pathways. **A)** Design of the optimal pathway enrichment strategy: 1) Gene set definition by pruning the Reactome pathway database, 2) Gene set enrichment optimal strategy combines GSEA and ROntoTools by selecting pathways which are significant in at least one of the methods (i.e.: OR rule). ORA was discarded to due to poorer performance. **B)** Proteomics enrichment ranks of *Chromatin remodeling* and *DNA repair* pathways, known to be dysregulated in SWI/SNF mutants (i.e., positive controls). Ranking was based on p-adj values obtained from each indicated enrichment method. For algorithm combination, the geometric mean of ranks obtained from each method individually were used. **C)** Proteomics enrichment results for pathways containing the SWI/SNF gene KO in the corresponding cell line. Cells are filled when the pathway contains the KO gene, and colors indicate the method that identified it. For the sake of visualization, *CREBBP*-only pathways have been cropped; see Supplementary Fig. 2F for full plot. **D)** Number of enriched pathways in proteomics dataset of each indicated mutant cell lines, as detected by GSEA or ROntoTools, using the pruned Reactome database. **E)** Number of enriched pathways in transcriptomics and proteomics datasets of indicated mutant cell line, as computed by the optimized enrichment pipeline.

Overall, the pruning strategy discarded 1802 pathways out of 2502 (72%), fulfilling the goal of drastically reducing the number of pathways while retaining 10999 out of 11360 (97%) genes documented in the Reactome database (Supplementary Fig. 3B-D, Supplementary Tables 4-5). The 361 discarded genes (3%) mostly corresponded to chemical reactions in large pathways, which were not represented in their children and lost in the top-down step as a result of pathway size constraints enforcement; 332 of them belonged to the *Gene Expression* category. Further, as this project aimed at identifying intracellular targetable vulnerabilities, we discarded nine out of the 29 pathway categories (i.e. 291 pathways, 1595 genes) associated with irrelevant physiological systems or organism-level functionalities (see Methods).

Next, we used the proteomic dataset to benchmark three gene set analysis methods - ORA, GSEA, and ROntoTools - and their combinations, based on their ability to predict the enrichment of (i) *Chromatin modifying enzymes* and *DNA repair,* two pathways known to be dysregulated in SWI/SNF mutants; and (ii) pathways containing the KO gene of each corresponding mutant^28^. Pathways were ranked based on their Benjamini-Hochberg adjusted p values (p-adj) in each method. For *Chromatin modifying enzymes*, ROntoTools and GSEA correctly assigned a high enrichment rank to the pathway, while ORA failed to identify it as a top hit (mean rank across SWI/SNF mutants 1.12, 4.48 and 13, respectively, Fig. 3B). Similarly, *DNA repair* was in the top 15 of dysregulated pathways according to ROntoTools or GSEA but was poorly ranked by ORA, further demonstrating the lack of accuracy of this last method. Next, we combined these enrichment tools two at a time by computing for each pathway, in each mutant, the geometric mean of the ranking obtained from each method. Combining ORA with the other methods harmed their performance so that the method was discarded. GSEA and ROntoTools were further benchmarked based on their performance to predict the enrichment of pathways containing the KO gene of each corresponding mutant (Fig. 3C, Supplementary Fig. 3E). Out of the 59 occurrences of KO gene in pruned Reactome pathways, 11 were flagged by both methods, six by GSEA only eight by ROntoTools only, and 34 by none, thus demonstrating the rationale of combining both algorithms to reduce false negative rate. *Transcriptional regulation by RUNX1*, a pathway involved in the differentiation of hematopoietic stem cells which may be dysregulated in acute myeloid leukemia (AML) - hence in our HAP-1 model - was flagged as significant by GSEA in six out of ten mutants, but not by ROntoTools. Conversely, in *SMARCA4*-, *BAP1*-, *CREBBP*-, *EED*- and *KMT2D*-KO cell lines, ROntoTools was able to detect eight pathways that GSEA did not identify. When looking at all pathways, we observed a limited agreement between ROntoTools and GSEA enrichment results in either the proteome (Fig. 3D) or transcriptome analysis (Supplementary Fig. 5F), with on average 20% of shared pathways (9.9% to 21,1% across SWI/SNF-mutants), further justifying the use of both methods combined.

Overall, our results suggested that even the latest enrichment methods yielded false negatives and we decided, as the best strategy for signal seeking, to combine ROntoTools and GSEA enrichment methods by retrieving pathways predicted as significant by either method.

### “Metabolism of proteins” is the most enriched category in SWI/SNF-mutant proteomics

Using our optimized pipeline, we performed pathway enrichment analysis of the transcriptome and proteome datasets (Fig. 4, Supplementary Fig. 4-6, Supplementary Tables 6-7). Hierarchical clustering based on pathway rankings identified two groups: SWI/SNF and non-SWI/SNF mutants for both proteome (Fig. 4A) and transcriptome (Supplementary Fig. 5A), in agreement with the clustering based on differential transcript or protein expression (Fig. 2A-B). However, pathway enrichment only modestly agreed between transcriptomics and proteomics analysis for each mutant (Fig. 3E, Supplementary Tables 6-7), in line with the moderate correlation found at the gene and protein level (Supplementary Fig. 2). Since proteins have a more direct functional relevance than transcripts in cell biology when limited RNA-protein correlation is observed, we first focused on the proteomic dataset.

**Figure 4.**
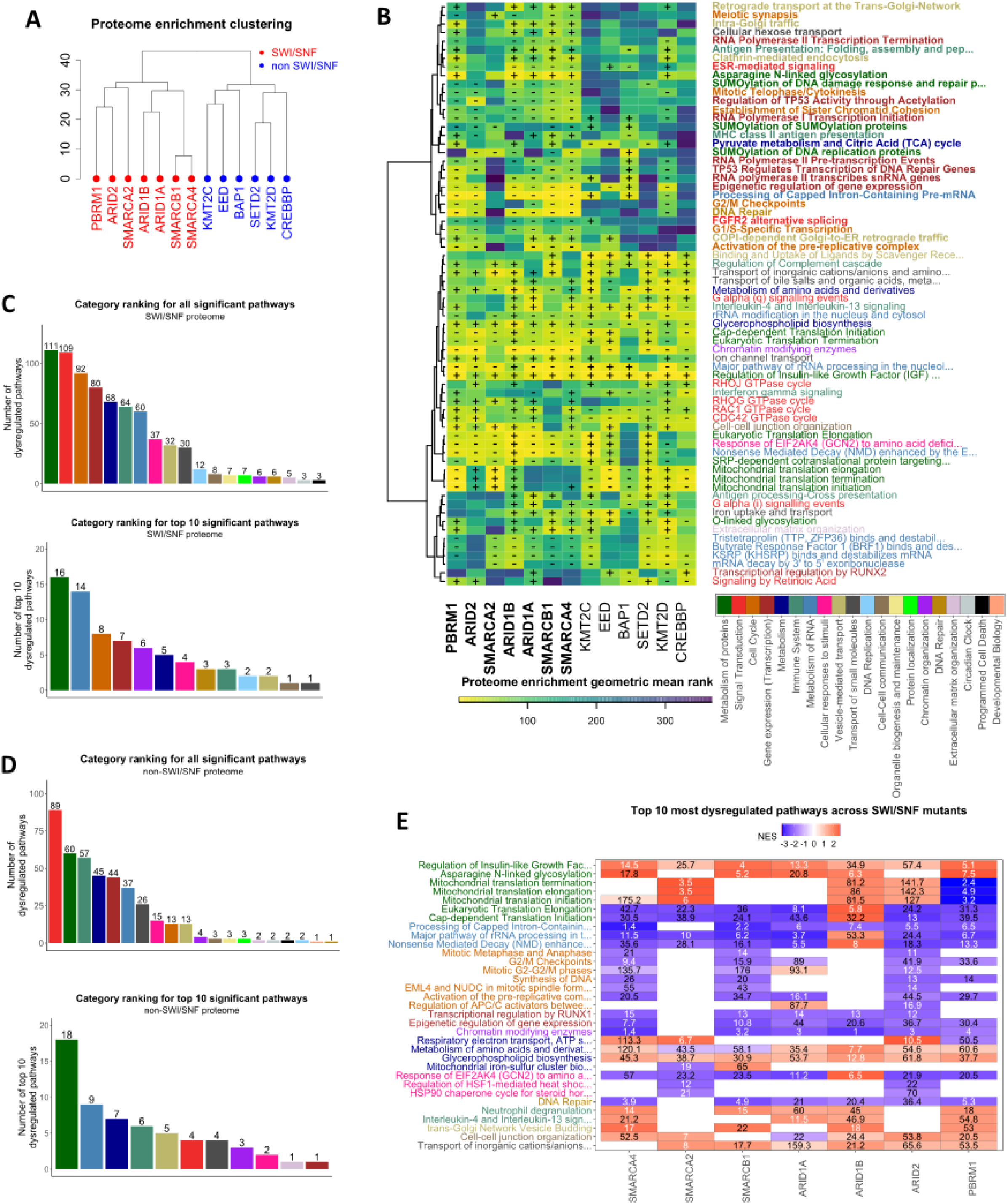
*Metabolism of proteins* is the most largely dysregulated category in the proteomics of SWI-SNF mutants and other enriched pathways are largely mutant-specific. **A)** Hierarchical clustering of mutant cell lines according to pathway ranking from proteomics enrichment analysis. **B)** Enriched pathways in at least five SWI/SNF mutants or at least four non SWI/SNF mutants. Symbols in each cell indicate if the pathway is significantly upregulated (+), downregulated (-) or not significant (empty), according to the NES score of GSEA. Pathway names are colored according to pathway categories. Pathways dysregulated in SWI/SNF mutants and not in non-SWI/SNF KO cell lines are highlighted in bold. Full pathways names are reported in Supplementary Table 7. **C)** Most dysregulated pathway categories in SWI/SNF mutants. Rankings correspond to the number of significantly enriched pathways for at least one SWI/SNF mutant, considering the whole pathway database (upper line) or only the Top10 most enriched pathways in each SWI/SNF mutant (lower line). **D)** Most dysregulated pathway categories in non-SWI/SNF mutants. Rankings correspond to the number of significantly enriched pathways for at least one non SWI/SNF mutant, considering the whole pathway database (upper line) or only the Top10 most enriched pathways in each SWI/SNF mutant (lower line). **E)** Top10 enriched pathways in SWI/SNF mutants. The cell colors indicate the pathway up- or downregulation (NES score of GSEA). The numbers inside the cells indicate the pathway ranking, in white, if the pathway is in the Top10 pathways of the mutant, and in black, otherwise. On the y-axis, pathways are arranged following the category rankings of panel B, and, within each category, according to the number of mutants for which the pathway appeared in the Top10. Full pathways names are reported in Supplementary Table 7.

To broadly describe enriched signaling pathways in SWI/SNF mutants, we analyzed them by pathway categories, defined as the largest pathway sets of the Reactome database. The most dysregulated category was *Metabolism of proteins*, which encompasses mechanisms related to protein folding and translation, peptide hormone metabolism, or post-translational protein modifications. It was followed by *Signal transduction*, *Cell cycle*, *Transcription, Metabolism, Immune system,* and *Metabolism of RNA* (Fig. 4B-C). When focusing only on the Top10 most dysregulated pathways for each SWI/SNF mutant, *Metabolism of proteins* remained the most frequently enriched category, followed by traditional SWI/SNF-regulated functions, i.e., *Metabolism of RNA*, *Cell Cycle Transcription*, *Chromatin organization,* and *Metabolism*. Interestingly, the four most broadly dysregulated pathways across mutants, within the *Metabolism of proteins* category, were related to post-translational modifications, including SUMOylation and glycosylation, potentially uncovering novel direct or indirect SWI/SNF functions (Fig. 4B). In the non-SWI/SNF group, *Signal transduction* was the most enriched category when accounting for all significant pathways, followed by *Metabolism of proteins* and *Immune system*. When using only the Top10 dysregulated pathways, M*etabolism of proteins* and *Metabolism of RNA* were the most enriched, followed by *Metabolism* and *Immune system* (Fig. 4D). *Cell cycle* and *DNA repair* were not enriched, and *Gene expression* has just one significant pathway, in contrast to the SWI/SNF group.

We next investigated the Top10 enriched pathways in each SWI/SNF mutant (Fig. 4E). As a quality check for the enrichment pipeline, *Chromatin-modifying enzymes* was the most dysregulated pathway (i.e. with the highest ranking) in each SWI/SNF mutant except for SMARCA2, in line with the very low expression of this subunit in HAP-1 cells. Three others of the Top10 dysregulated pathways were enriched in at least five SWI/SNF mutants, including potential non-canonical (*Major pathway of rRNA processing in the nucleolus and cytosol*) and novel (*Asparagine N-linked glycosylation*; *Nonsense Mediated Decay enhanced by the Exon Junction Complex*) SWI/SNF functions.

The most populated category was *Metabolism of proteins*, encompassing seven Top10 pathways (Fig. 4C, E, Supplementary Table 7). The most frequently dysregulated one was *Regulation of Insulin-like Growth Factor transport and uptake by insulin growth factor binding proteins*, which was consistently upregulated in all SWI/SNF mutants, with 13 (out of 124) genes being common to the GSEA leading edges of all mutants (Supplementary Fig. 3F, Supplementary Table 7). The second most dysregulated pathway, *Asparagine N-linked glycosylation*, was upregulated in all SWI/SNF mutants except *ARID2*- and *SMARCA2*-KO, with 61 (out of 305) genes being common to the GSEA leading edges across mutants. Intriguingly, this process, which corresponds to the addition of glycans to peptides folded in the endoplasmic reticulum (ER) and subsequent transport to the Golgi apparatus, had not been previously related to SWI/SNF functions. The other five dysregulated pathways of the *Metabolism of proteins* category were directly linked to protein translation, with two pathways occurring in the cytoplasm (*Cap-dependent translation initiation* and *Eukaryotic Translation Elongation*), and three of them related to mitochondrial protein metabolism (*Mitochondrial translation initiation, elongation, and termination*), reminiscent of the role of SWI/SNF subunits in OxPhos^42^. Discrepancies between subunit defects could also be observed: for example, mitochondrial metabolism-related pathways were all upregulated in the *PBRM1*-KO cell line, whereas it was the opposite in the *SMARCA2*-KO; *Cap-dependent translation initiation* and *Eukaryotic Translation Elongation* were both upregulated in the *ARID1B*-KO but downregulated in all other SWI/SNF mutants; and pathways related to mitochondrial translation were downregulated in *PBRM1*-KO and upregulated in the *SMARCA2*-, *ARID1B*- and *ARID2*-KO cell lines.

The following three most enriched Top10 pathways belonged to the *Metabolism of RNA* category and included: (i) *Processing of Capped Intron-Containing Pre-mRNA* , (ii) *Major pathway of rRNA processing in the nucleolus and cytosol*, and (iii) *Nonsense Mediated Decay enhanced by the Exon Junction Complex* which were downregulated in all SWI/SNF mutants, but SMARCA2 for the first pathway, and except for *ARID1B* where both latter ones were upregulated. The next seven most enriched Top10 pathways were part of the *Cell Cycle* category: six of them were linked to activation of the G1/S, G2/M checkpoints or mitosis, and notably identified in the *ARID1A*- and *ARID2*-KO cell lines, in line with the known role of these DNA-binding subunits in cell cycle control^19,43^, as well as in the *SMARCA4*-KO mutant, consistent with the described increased replication stress and ATR inhibitor sensitivity when *SMARCA4* is lost^17^. Both subsequent Top10 pathways belonged to the *Gene Expression* category and included *Transcriptional Regulation by RUNX1* (most likely HAP1 cell type-specific considering the role of the master transcriptional regulator RUNX1 in hematopoiesis and acute myeloid leukemia) and *Epigenetic regulation of gene expression*, most likely cell type-independent. They were dysregulated in all cell lines but the SMARCA2 KO one. Finally, the last four most enriched Top10 pathways belonged to the *Metabolism* category and notably included the *Respiratory electron chain* which, intriguingly, was upregulated in the *ARID2*-, *SMARCA2*- and *SMARCA4*-KO cell lines, and downregulated in the *PBRM1*-KO. In the non-SWI/SNF group, the most enriched Top10 pathways belonged to the *Metabolism of Proteins*, *Metabolism of mRNAs* and *Metabolism* categories (Fig. 4D)

Pathway enrichment analysis was further performed for the transcriptome dataset (Supplementary Fig. 5). When comparing category rankings for SWI-SNF mutants only, *Signal Transduction* was the most dysregulated category, followed by *Immune System, Metabolism of proteins, Gene Expression* and *Cellular response to stimuli* (Supplementary Fig. 5C), which were distinct from the proteomic enrichment results.

Surprisingly, none of the pathways of the *Chromatin organization* category were found enriched in any mutants (Supplementary Fig. 5B-E). Contrary to the modest overlap across enriched pathways which was found between proteomic and transcriptomic data when considering all significantly dysregulated pathways (Fig. 3E), focusing on the Top10 most dysregulated pathways for each SWI/SNF mutant allowed us to consistently identify *Metabolism of proteins* as the most frequently enriched category in both datasets (Fig. 4C, Supplementary Fig. 5C). By contrast, while *Metabolism of RNA* was the 2^nd^ most enriched category in the proteome Top10 category ranking, it occupied the 7^th^ position in the transcriptome dataset, with only three significantly enriched pathways (Supplementary Fig. 5C). When comparing the proteome and transcriptome Top10 enriched pathways in each SWI/SNF-mutants, we could identify 14 common pathways, out of 33 in the proteome and 40 in the transcriptome datasets (Fig. 4E and Supplementary Fig. 5E). Those included: *Asparagine N-linked glycosylation*, *Regulation of Insulin-like Growth Factor Receptors* and *Mitochondrial translation* (*Metabolism of proteins* category); *Processing of capped intron-containing pre-mRNA* and *Major pathway of rRNA processing* (*Metabolism of RNA*); *Glycerophospholipid biosynthesis* and *Respiratory electron transport* (*Metabolism*), and *Neutrophil degranulation* and *Insulin-4 and 13 signaling* (*Immune system*).

Overall, our new enrichment pipeline applied to the proteomic and transcriptomic data of our SWI/SNF-mutant isogenic panel allowed us not only to recapitulate known SWI/SNF-dependent pathways (e.g., cell cycle regulation and transcriptional regulation) but also to identify novel ones, such as *Metabolism of proteins*-, *Metabolism of RNA-* and *Mitochondrial metabolism*-related pathways.

### Drug screening identifies inhibitors of BRD9, protein synthesis, and histone modifiers as being selectively synthetic lethal with defects in SWI/SNF subunits

To further identify targetable vulnerabilities associated with SWI/SNF defects, we used a functional, orthogonal approach and performed high-throughput drug screening on the HAP1 panel using two libraries: (i) the Prestwick library (1200 off-patent small molecule inhibitors, suitable for drug repurposing), and (ii) the SelleckChem Epigenetics library (186 compounds), selected based on the hypothesis that synthetic lethalities may mostly occur in-between epigenetic pathways. We first performed a unidose screening, where cell viability was assessed after exposure to 10 μM of each drug for five days for the Prestwick library, and for 5 and 11 days for the epigenetic library, considering the longer time required for these drugs to rewire signaling and impact cell fitness (Fig. 5A). Hits were identified based on their ability to selectively kill SWI/SNF- or chromatin remodeling-mutant HAP1 cells while sparing the HAP1 parental (HAP1-WT) cell line (see Methods). The cytotoxicity of compounds of the Prestwick library was highly mutant-dependent, with only the estrogen antagonist Danazol affecting two or more SWI/SNF mutant cell lines (Fig. 5B, Supplementary Table 8). By contrast, in the Epigenetics library, four small molecule inhibitors selectively killed SWI/SNF mutants (Supplementary Table 9). As a benchmarking and quality evaluation, we could identify previously described synthetic lethalities: for example, the P300 inhibitor C646 was synthetic lethal in the *CREBBP*-KO and *ARID1A*-KO cell lines (with 86% and 74% reductions in survival fraction (SF) as compared to the parental HAP1-WT cell line)^44,45^. Interestingly, C646 was also selectively toxic to all other SWI/SNF mutants (*SMARCA2/4, ARID1B/2, PBRM1*-KO), as well as *KMT2C/2D-, SETD2-, EED-* and *BAP1*-KO, with reductions in SF ranging from 50% to 78%, thereby representing potential novel potent synthetic lethal interactions. Resveratrol, a flavonoid used as an anti-tumoral agent^46,47^, also selectively killed the *ARID1B*-, *PBRM1*-, and *SMARCA2*-KO cell lines, with reductions in SFs ranging from 48 to 84%.

**Figure 5.**
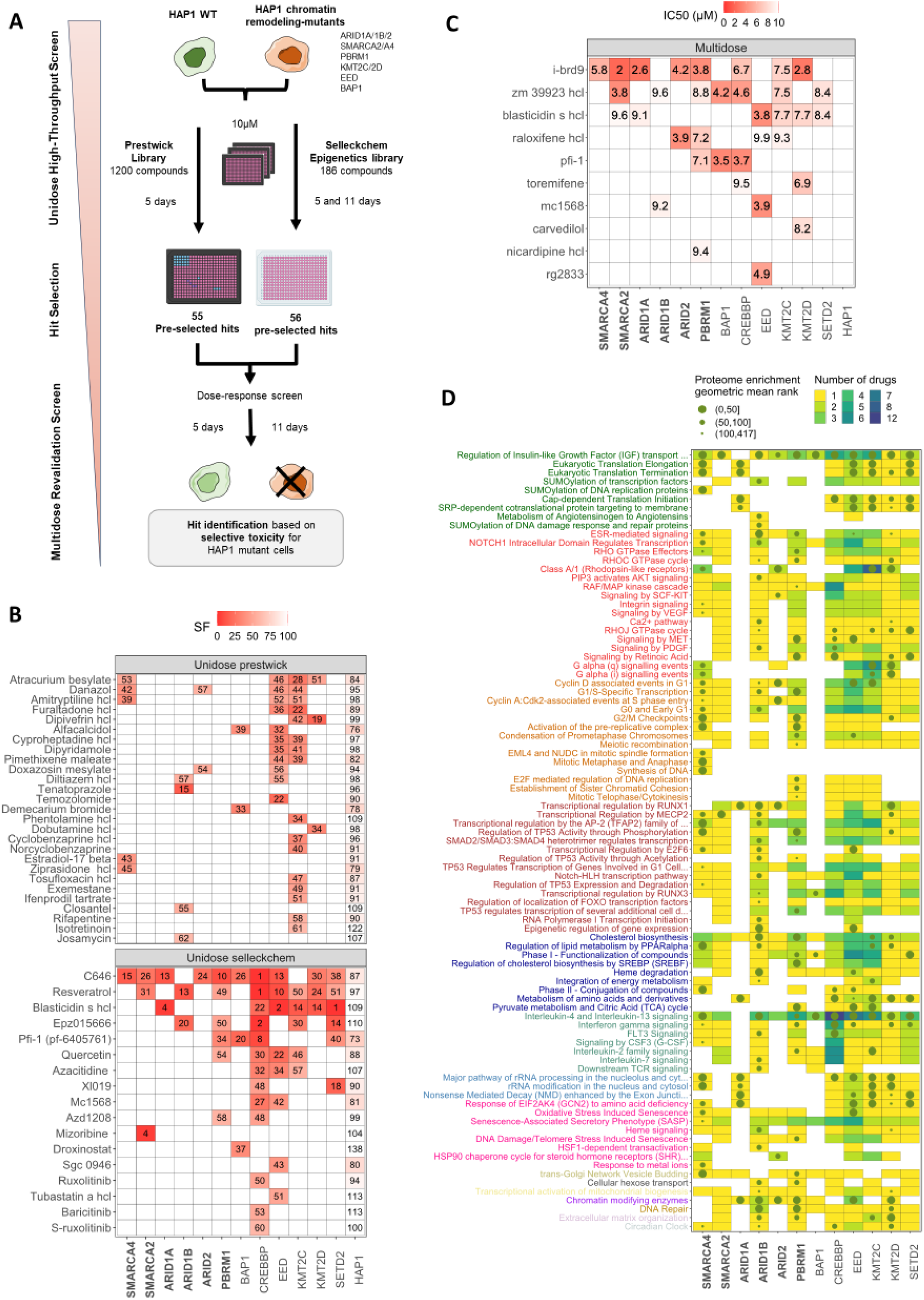
Drug screening reveals potential drug targets in SWI/SNF-KO cell lines. **A)** Drug screening strategy. Cell line sensitivity to a unidose drug screening of 1200 compounds (Prestwick Chemical Library) and 186 small molecule inhibitors (SelleckChem Epigenetic Library) was assessed; 55 and 56 hits were selected, respectively, for revalidation in multidose assay (see Methods). **B)** Unidose drug screening. Survival fractions (SF), normalized on control, are depicted for HAP1-mutant or HAP1-WT cell lines. Drugs are ranked by increasing number of sensitive cell lines and average SF across cell lines. **C)** Multidose drug screening. Inhibitory Concentration (IC)50 values of drugs that were selectively toxic towards HAP1-mutant cell lines are depicted; white squares indicate that IC50 was not reached. **D)** Enriched pathways in genes interacting with at least one HAP1-mutant selective drug of the unidose or multidose analyses for each cell line individually. Cell colors refer to the number of drugs interacting with genes of the pathway. Green dots represent the rank of the pathway in the proteome enrichment analysis whenever predicted as dysregulated. Pathways are colored based on Reactome categories and categories are arranged on the y-axis according to their enrichment in SWI/SNF-KO cell line proteomics (Fig. 4C).

The most promising compounds were further selected for revalidation in dose-response (“multi-dose”) medium throughput screening, based on: (i) the selective toxicity towards SWI/SNF- or other chromatin remodeling-mutant HAP1 cells; (ii) redundancy of hits targeting the same pathway; and (iii) potential for subsequent clinical development (see Methods; Supplementary Table 10). Using these criteria, 55 and 56 compounds were selected for revalidation from the Prestwick and Epigenetic drug libraries, respectively. In that dose-response screen, we defined “hits” as chemicals whose IC_50_ was smaller than the highest dose screened (10 µM) in a SWI/SNF or non-SWI/SNF KO cell line and were not toxic towards the HAP1-WT cell line (SF > 70% for any drug concentration in both replicates). Four main compounds reached these criteria (Fig. 5C). The top hit was i-brd9 (GSK602, an inhibitor of the GBAF-specific subunit BRD9), which scored in all but one SWI/SNF-KO cell lines; interestingly, inhibition of BRD9 was shown to be synthetic lethal with cBAF and PBAF SWI/SNF defects concomitantly to our screen^48^, allowing us to use that top hit as a positive quality control of our screen, and supporting the ability of our HAP1 isogenic panel to serve as signal-searching model for cell type-independent targetable vulnerabilities. By contrast, the top second hit, the JAK inhibitor zm39923 HCL, was potentially cell-type-specific, considering the role of JAK/STAT signaling in cells of the myeloid lineage, such as HAP1. Interestingly, the third hit was the cytidine analog blasticidin, an antibiotic that acts as a protein synthesis inhibitor by inhibiting the termination step of protein translation and peptide bond formation by the ribosome, thereby making a potential link between the drug screen and the dysregulation previously identified in the *Metabolism of proteins* pathway category by proteomic profiling (Fig. 4B-C). The fourth hit, Raloxifene HCL, an estrogen receptor (ER) modulator (close to Danazol) that inhibits the human cytosolic aldehyde oxidase-catalyzed phthalazine oxidation activity, was selectively toxic to the PBAF-mutant *ARID2*- and *PBRM1*-KO cell lines. Raloxifene has ER-dependent and ER-independent mechanisms of action, including modulation of cell metabolism and mitochondrial function, potentially underlying its selectivity in SWI/SNF-KO HAP1 cells, and in line with the dysregulation of *Mitochondrial Metabolism* previously identified by proteomic profiling (Fig. 4B)^49^. The fifth hit was the *BRD4* inhibitor PFI-1, which had previously been described as synthetic lethal with *SMARCA4* defects^50^, and also scored as hit in *PBRM1*-, *BAP1*-, and *CREBBP*-KO clones on our screen (Fig 5B, 5C). This novel synthetic lethality occurring at a similar magnitude in the *BAP1*- and *CREBBP*-KO clones was in line with the newly described CREBBP-like role of *BAP1* as a transcriptional activator in the *BAP1* complex^51^.

We next sought to explore drug-targeted pathways by using drug-gene interaction databases^52–55^ to identify genes interacting with hits of our drug screens (see Methods). Enriched pathways targeted by drug hits belonged to various categories, notably *Metabolism of proteins*, *Signal transduction*, *Cell Cycle*, *Gene expression*, *Metabolism,* and *Immune system* (Fig. 5D). These pathway categories were broadly concordant with the ones identified by proteomics profiling (Fig 4). Still, there was only a partial overlap between individual pathways predicted as targetable by drug screening and as dysregulated in proteomics (22%; SD = 17%), highlighting the complementarity of both molecular profiling (Fig. 2,4) and functional (Fig. 5) approaches. Among pathways that were both drug-targeted and enriched in proteomics, we could identify *Interleukin-4 and -13 signaling*, in line with the frequent identification of JAK/STAT inhibitors within the top hits of both drug screens, *Regulation of Insulin-like Growth Factor (IGF) transport* (dysregulated in 11 out of 12 mutant cell lines), and *Chromatin-modifying enzymes* (targetable in 10 mutant cell lines, and significant in the proteome enrichment of six of them), potentially favored by the enrichment in epigenetic drugs in our initial and revalidation screen.

In conclusion, high throughput drug screening enabled to confirm known synthetic lethalities with SWI/SNF defects, and to newly identify compounds selectively targeting SWI/SNF subunit-KO cell lines, including several epigenetic drugs, compounds interfering with mitochondrial metabolism, and small molecule inhibitors of ribosomal translation and protein metabolism.

### CRISPR DepMap project analysis identifies EP300 and mitochondrial targets as novel genetic vulnerabilities in a non-isogenic panel of cell lines with low expression of SWI/SNF subunits

To next investigate whether our findings could be extended to large, independent datasets of non-isogenic cancer cell lines from various cell-of-origins, we used the publicly available resource of the DepMap project^56^ and developed an algorithm to infer synthetic lethal interactions between defects in SWI/SNF subunits and any other gene in the DepMap whole genome CRISPR KO high-throughput screen (see Methods, Fig. 6A). To that end, we compared CRISPR effect scores between cell line groups selected based on the level of expression of genes encoding the SWI/SNF subunit or chromatin remodeler of interest^57^. Briefly, we: (i) estimated the distribution of the expression of each gene encoding a SWI/SNF subunit across the CCLE panel using transcriptomic data (Supplementary Fig. 7), since proteomics were only available for a subset of the cell lines ; (ii) defined two groups corresponding to cell lines with the 10% highest or lowest expression for the gene of interest; and (iii) compared between both groups the CRISPR score of each gene knocked out within the genome-wide CRISPR screen. We defined gene hits as the ones with adjusted p-value < 1e-3 and Wilcox displacement > 0.15, and super hits as those with p-adj < 1e-4 and Wilcox displacement > 0.15. Overall, gene hits and superhits were largely subunit defect-specific: 10 superhits and 28 hits were identified for two or more SWI/SNF subunits, whereas 49 superhits and 93 hits were private to one unique SWI/SNF subunit defect (Fig. 6B). As a quality control of our newly developed method, we could identify SMARCA4 as the top hit for cell lines with low *SMARCA2* expression (Supplementary Table 11). Thirty-one super hits and six hits were shared between SWI/SNF and non-SWI/SNF defective cell lines, while eight superhits and 41 hits were detected in a single non-SWI/SNF defect (Fig. 6C, Supplementary Table 11). The most represented super hit across SWI/SNF lines was E1A Binding Protein P300 (EP300), which was predicted as being synthetic lethal in six SWI/SNF-defective (*ARID1A/1B/2*, *PBRM1*, *SMARCA4/B1*) and three non-SWI/SNF-defective (*BAP1*, *KMT2D/2C*) populations. EP300 synthetic lethality was also close to significant for CREBBP low expression (Wilcox displacement>0.15 but p-adj=0.1, Supplementary Table 11). EP300 is a histone acetyltransferase that belongs to the same family and has a similar structure and function to its cognate co-activator CREBBP; both activate transcription through interactions with transcription factors and histone acetylation relaxing chromatin structure at gene promoters and enhancers. Interestingly and consistent with this data, our high-throughput drug screen previously identified the C646 EP300 inhibitor as one of the top hits, causing selective cell death in all SWI/SNF and non-SWI/SNF-KO mutants, with the strongest synthetic lethal interaction being – as expected – in the *CREBBP*-KO cell line (Fig. 5B and Supplementary Table 11). This match between our high-throughput drug screen on isogenic HAP1 cell lines, and the DepMap CRISPR screen on a non-isogenic panel of high/low chromatin remodeling expression cell lines from diverse genetic backgrounds, reinforced the relevance of our HAP1 model for signal-searching.

**Figure 6.**
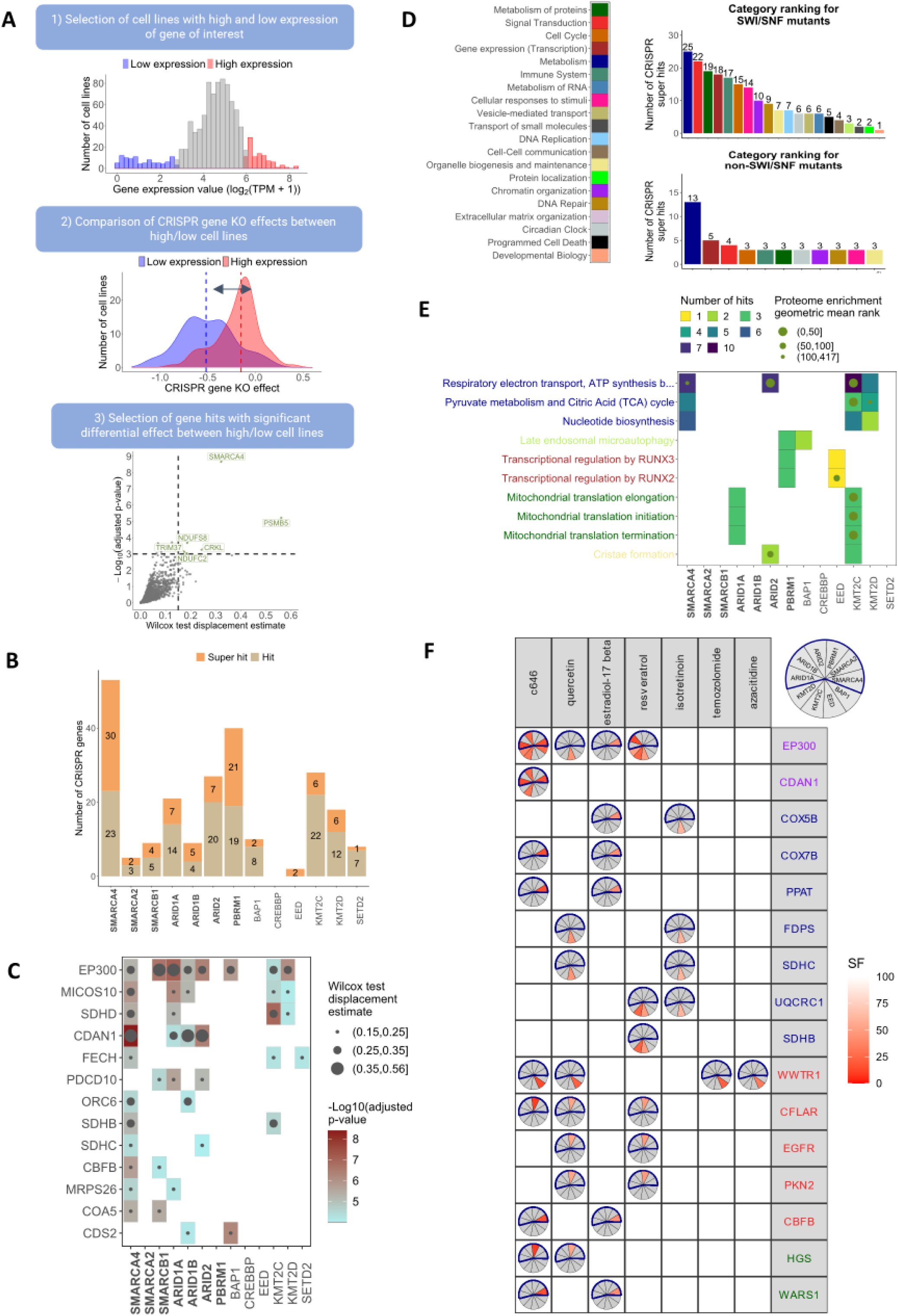
CRISPR DepMap project analysis identifies potential synthetic lethalities associated with SWI/SNF deficiencies. **A)** Scheme of the algorithm developed to predict synthetic lethal interactions. 1) A mutant-like gene of interest is chosen (e.g.: *SMARCA2*) and cell lines in the highest and lowest 10% quantiles are selected to form the high and low expression groups, respectively. 2) Given a gene KO by CRISPR (e.g. *EP300*), a Wilcoxon statistical test is run to evaluate the difference in the EP300 CRISPR score between the low and high expression groups of cell lines. If the test is significant and if the displacement between CRISPR score distributions of high and low expression groups is in the desired direction (i.e. *SMARCA2* low expressing cell lines are more sensitive to the deletion of *EP300* as compared to the high expression group), a synthetic lethal interaction is found. Significance is defined as Wilcox displacement estimate being > 0.15 and p-adj < 1e-3 for “hits” and p-adj < 0.0001 for “super-hits”. 3) This procedure was iteratively performed for the 13 chromatin remodelers of interest, set as the mutant-like gene, explored CRISPR genes corresponding to those of the Reactome database. Results can be displayed in a volcano plot where the dotted lines represent significance thresholds for hits. **B)** Number of super hits and hits for each cell line. **C)** Super-hit CRISPR genes for each cell line. Genes are ranked in decreasing order with respect to the number of affected mutants. **D)** Top dysregulated Reactome pathway categories considering the super-hits predicted across SWI/SNF mutants (upper line) or non-SWI/SNF mutants (lower line). **E)** Enriched pathways in the hit gene list for each cell line. The color code indicates the number of hits found in each pathway. The green dots represent the rank of the pathway in the proteome enrichment analysis whenever it was predicted as dysregulated. Pathways are ranked in decreasing order of the number of cell lines where they are found significant. **F)** Comparison between CRISPR analysis and drug screening. Genes identified as hits in the CRISPR study were compared with genes interacting with drugs identified as efficient in the drug screening, for each cell line individually. Drug-gene interactions were obtained from public databases (see Methods). Only genes being CRISPR hits for at least two cell lines or being targeted by at least two efficient drugs are depicted. Drugs are ordered based on the number of targeted CRISPR hits. Genes are colored according to the Reactome category they belong to (see panel D).

Strikingly, five out of the ten following superhits that scored in at least two SWI/SNF-KO cell lines (namely MICOS10, SDHD, SDHB, SDHC and MRSP26), were related to mitochondrial metabolism. MICOS10 is a protein that plays a key role in the formation of mitochondrial cristae, which are folds of the inner mitochondrial membrane where energy production occurs^58^. SDHB/C/D are subunits of the Succinate Dehydrogenase Complex which is part of the mitochondrial inner membrane protein of complex II of the respiratory chain, responsible for the oxidation of succinate in the oxidative phosphorylation and the TCA cycle. Finally, the *MRPS26* nuclear gene encodes Mitochondrial Ribosomal Protein S26, which participates in protein synthesis within the mitochondria. Additionally, *FECH* and *CDS2*, two genes related to mitochondrial metabolic pathways (*Heme biosynthesis* and *Glycerophospholipid biosynthesis*, respectively), were predicted as synthetic lethal with *SMARCA4* and *ARID1B* KO respectively (Fig. 6C). Altogether, this pointed towards a novel genetic vulnerability of SWI/SNF-defective cancer cells, which would be driven by an altered mitochondrial respiration and protein metabolism, as previously identified in our proteomics dataset.

Superhits for both SWI/SNF and non-SWI/SNF KO lines were mapped to their respective Reactome pathways to allow comparison with proteomics enrichment results in HAP1 cell lines. *Metabolism* was the most targetable category across all defects. In the SWI/SNF group, it was followed by *Signal transduction*, *Metabolism of proteins*, *Gene expression* and *Immune system* (Fig. 6D). In the non-SWI/SNF group, the subsequent targetable genes were almost evenly spread over 10 different categories. At the pathway level, processes related to mitochondrial respiration and protein metabolism (*Respiratory electron transport, TCA cycle*, *Mitochondrial translation*, *Cristae formation*) were the most enriched across SWI/SNF and *KMT2C/2D*-low expressing cell lines, in line with our proteomics data of the HAP1 isogenic panel (Fig. 6E).

To next explore whether genes predicted as targetable by the CRISPR DepMap analysis would be involved in pathways targeted by compounds identified as hits of the drug screening, we compared, for every clone, the list of CRISPR hits and the list of genes known to interact with hit compounds (Fig. 6F, Supplementary Figures 8-20, Supplementary Tables 12,13). This enabled us to identify 16 genes that interacted with seven drugs. The first one was *EP300*, the main target of the c646 drug, which was predicted as a super hit in the *ARID1A*-, *ARID2*-, *BAP1*-, *KMT2D*- and *SMARCA4*-low expressing cell lines (Fig. 6C). Interestingly, EP300 is also known to interact with other hits of the drug screen, including flavonoids quercetin, resveratrol, as well as estradiol-17 beta. The following seven genes were involved in cellular metabolism, of which five (*COX5B*, *COX7B*, *SDHB*, *SDHC,* and *UQCRC1*) were part of the mitochondrial respiratory chain. Two additional genes were involved in *Metabolism of proteins*: *HGS*, which plays a role in protein deubiquitination and cell signaling by Receptor Tyrosine Kinases, specifically EGFR and Rho GTPases, and *WARS1* which encodes a tryptophanyl-tRNA synthetase, an enzyme involved in cytosolic protein translation.

In summary, analysis of the DepMap project genome-wide CRISPR screen allowed us to identify multiple SWI/SNF defects-associated genetic vulnerabilities that corresponded either to pharmacologic vulnerabilities identified in our high-throughput drug screening on the HAP1-isogenic panel (e.g., *EP300* and C646), or to pathways dysregulated in proteomics profiling (e.g. *MICOS10*, *SDHB/C/D* and mitochondrial metabolism), overall pointing towards “orthogonal” synthetic lethal dependencies that may be the most robust and independent from the cellular genetic background.

### SMARCA4-defective cells are vulnerable to inhibition of EP300 and mitochondrial respiration

We next thought to explore whether the above-identified “orthogonal” synthetic lethal dependencies would also operate in isogenic histotype-relevant models and decided to focus on deficiencies of SMARCA4, a pivotal SWI/SNF catalytic subunit frequently mutated across multiple solid tumors. To do so, we created two *SMARCA4* isogenic models in cancer cell types where *SMARCA4* is found mutated: non-small cell lung cancer (H358 cell line), and sarcoma (U2OS cell line). Each model was constituted of one *SMARCA4*-WT parental cell line, and one *SMARCA4*-KO cell line where *SMARCA4* had been disabled by CRISPR-Cas9 gene editing (Fig. 7A). For both cell lines, knocking out *SMARCA4* resulted in an increased protein expression of ARID1A/1B and ARID2 and a decreased PBRM1 expression, as observed in the HAP1 model for the latter (Fig. 7B). We next assessed the sensitivity of each model to compounds identified as “orthogonal” hits: the selective CBP/EP300 inhibitor CPI-637^59^ and two mitochondrial respiratory chain poisons: antimycin and oligomycin (which inhibit complexes III and V, respectively). Short-term drug sensitivity assays did not show any significant therapeutic window, in line with our previous unidose drug screen and the need for longer exposure to epigenetic drugs to rewire the cell epigenome and impact cellular fitness (Supplementary Fig. 21A). By contrast, long-term colony formation assays confirmed that the *SMARCA4*-KO cell line of both models was selectively sensitive to CPI-637, with IC_50_ values respectively 2.9-fold and 7.8-fold smaller in H358 and U2OS SMARCA4-KO cells as compared to corresponding parental cells (two-way ANOVA p<0.001, Fig. 7C, Supplementary Fig 21). This suggested that CBP/EP300 inhibition could represent a novel synthetic lethal therapeutic strategy for SMARCA4-defective tumors. Similarly, antimycin was selectively toxic towards the *SMARCA4*-KO cell line of both models in short-term exposure, consistent with the more rapid action of drugs targeting cell metabolism (IC_50_ values respectively 43-fold and 1.1-fold smaller in H358 and U2OS *SMARCA4*-KO than in parental cells, two-way ANOVA p<0.001, Fig. 7D). Intriguingly, the *SMARCA4*-KO H358 cell line was profoundly more sensitive to oligomycin than the parental cell line (short-term IC_50_ : 2.63 µM for WT *vs* 0.0647 nM for *SMARCA4*-KO; long-term IC_50_ : 1.65 nM for WT *vs* 0.14 nM for *SMARCA4*-KO; two-way ANOVA p<0.001, Fig. 7E-F), while a therapeutic window was inconstantly observed in the U2OS cell line (Supplementary Fig. 21), suggesting a potential variability and the contribution of additional factors for this latter synthetic lethal interaction.

**Figure 7.**
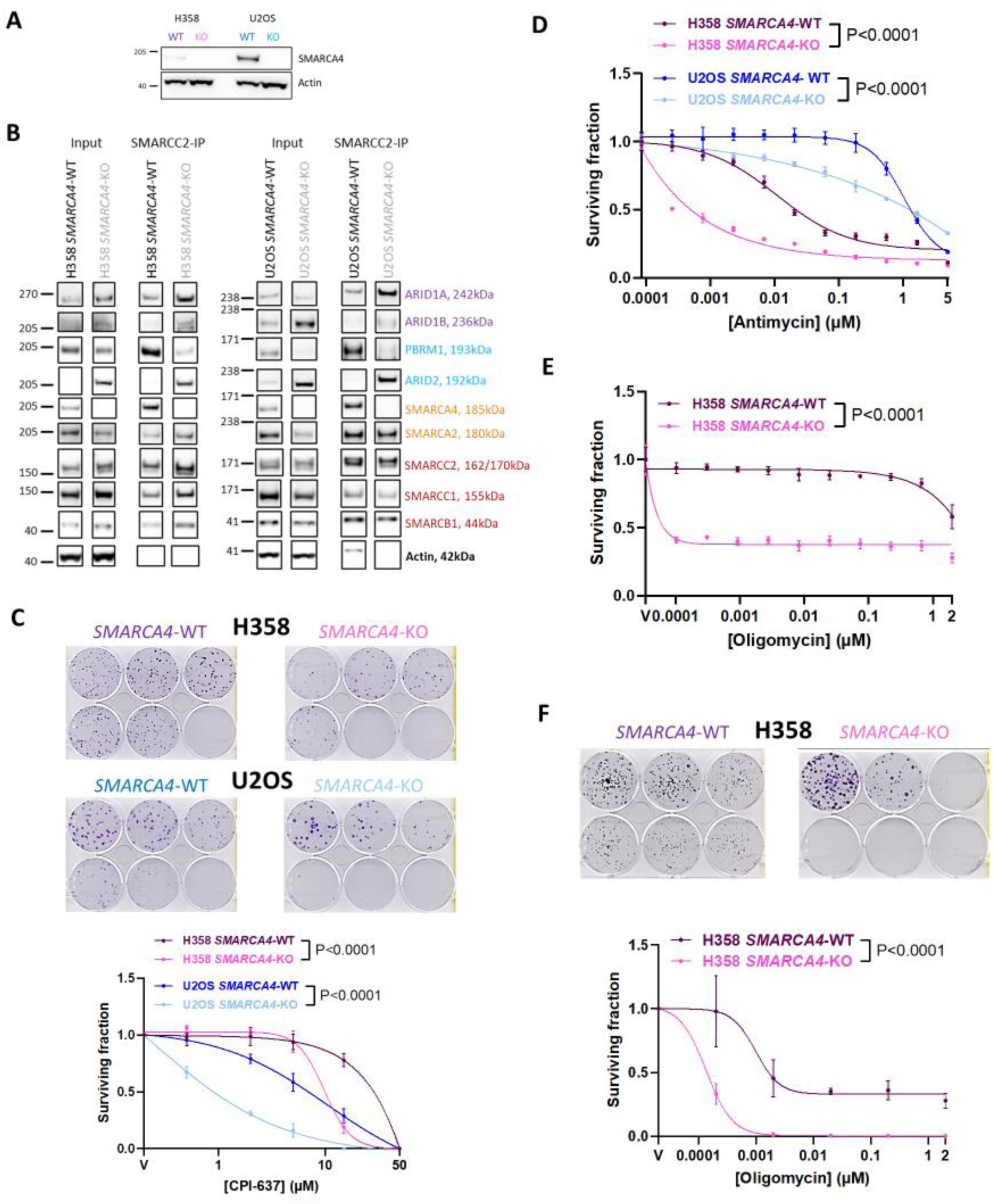
SMARCA4-defective cells are vulnerable to inhibition of EP300 and mitochondrial respiration. **A)** Western blot of SMARCA4 expression on total cell extracts of H358 *SMARCA4*-WT or -KO, U2OS *SMARCA4*-WT or -KO. **B)** Western blot on total cell extracts of H358 *SMARCA4*-WT or -KO, U2OS *SMARCA4*-WT or -KO of selected SWI/SNF subunits following immunoprecipitation (IP) of the SMARCC2 core subunit. **C)** Dose-response clonogenic survival curves and representative pictures of *SMARCA4*-isogenic models exposed to increasing concentrations of CPI-637 for 10 days (U2OS cells) or 13 days (H358 cells). Mean ± SD, number of replicates per data point *n* = 3; two-way ANOVA and post hoc Sidak’s test. **D)** Dose-response survival curves of *SMARCA4*-isogenic models exposed to increasing concentrations of mitochondrial respiration complex III inhibitor Antimycin A for 7 days. Mean ± SD, n = 3; two-way ANOVA and post hoc Sidak’s test. **E)** Dose-response survival curves of H358 *SMARCA4*-isogenic cells exposed to increasing concentrations of mitochondrial respiration complex V inhibitor Oligomycin A for 7 days. Mean ± SD, n = 3; two-way ANOVA and post hoc Sidak’s test. **F)** Dose-response clonogenic survival curves and representative pictures of H358 *SMARCA4*-isogenic cells exposed to increasing concentrations of Oligomycin A for 17 days. Mean ± SD, *n* = 3; two-way ANOVA and post hoc Sidak’s test.

Overall, the SMARCA4-EP300 and -mitochondrial respiratory chain complexes synthetic lethal dependencies could be revalidated in independent cancer-relevant models, highlighting their robustness and potential clinical translatability.

## DISCUSSION

Mutations in subunits of the SWI/SNF chromatin complex occur in 20% of solid tumors and still represent a highly unmet need. In this study, we integrated multi-omics profiling and drug screening of HAP1 isogenic cell lines mutated for individual SWI/SNF subunits or other chromatin modifying enzymes, with CRISPR screens from the DepMap project to investigate the consequences of SWI/SNF mutations on gene expression and intracellular pathway dysregulation, and subsequently identify targetable genetic vulnerabilities. We uncovered that the defects in several SWI/SNF subunits, including SMARCA4, led to a dependency on the protein acetyltransferase CBP/EP300 and adequate mitochondrial respiration, which we revalidated in independent disease-relevant SWI/SNF-defective tumor models.

Targeting tumor loss-of-function is often more challenging than targeting oncogene addiction, and identification of synthetic lethal relationships or genetic vulnerabilities is key to uncovering novel therapeutic approaches. The choice of the model in which such discoveries are performed is key: while clean isogenic models allow to confidently link the observed phenotype with the genetic alteration, they do not enable to explore the cell type-dependency of genetic vulnerabilities and non-isogenic models better recapitulate the tumor reality and heterogeneity. Here, we show that isogenic HAP1 models, despite being haploid and originating from chronic myeloid leukemia - a histology where SWI/SNF is usually not mutated - represent a useful tool for the initial identification of genetic dependencies that are cell type-independent. Indeed, we could confidently revalidate several genetic dependencies identified using multi-omics profiling and high-throughput drug screening on our HAP1 isogenic panel, not only in the DepMap project CRISPR screen, performed on independent non-isogenic cell lines, but also in additional disease-relevant models. Among them, our strongest uncovered genetic dependency was the synthetic lethality between EP300 pharmacological inhibition (or *EP300* genetic depletion) and SWI/SNF subunit defects, including SMARCB1/A4 and ARIDA1/1B/2. Interestingly, such synthetic lethality was also extremely recently found to operate between SMARCB1 or ARID1A defects and the dual inhibition of CBP/p300 in unrelated models^60–62^, thereby reinforcing the robustness of our findings. Several CBP or EP300 small molecule inhibitors are currently being tested in patients with advanced solid tumors or hematological malignancies (e.g., NCT06433947, NCT05488548), opening the way to clinical translation of our findings.

Differential expression analysis between SWI/SNF-mutated *versus* -WT cell lines and pathway enrichment both concluded on a weak correlation between alterations of the transcriptome and proteome, suggesting that transcript levels were only moderately predictive of protein expression, as reported in the literature for various species and tissues^63–68^. Still, the correlation between RNA and protein expression may vary depending on the cellular process in which genes are involved^69^. In this study, the *Chromatin modifying enzymes* pathway, together with several DNA damage response and Cell Cycle processes, were found enriched in SWI/SNF mutants in the proteomics analysis whereas they were not identified in the previous transcriptomics studies (ARID1A^20,70,71^, ARID1B^71^, ARID2^72,73^, PBRM1^74–76^, SMARCA2/SMARCA4^77,78^ and SMARCB1^79^). These results implicate the role of post transcriptional regulation underscoring the weaker predictive capacity of the latter dataset for pathway functionality.

Optimal integration of large-scale multi-omics datasets and functional high-throughput screening requires the design and use of robust bioinformatics pipelines. Here, we developed an optimized pathway enrichment strategy based on two improvements: (i) a biologically-sound reduction of the number of pathways tested for enrichment, and (ii) an optimization of the enrichment statistical algorithms (see Methods). By applying this new pipeline to proteomics datasets, we uncovered novel dysregulations in SWI/SNF-defective cell line models, which were linked to *Metabolism of proteins*, including protein sumoylation, glycosylation, and mitochondrial translation. Such pathway alterations were further supported by the selective toxicity of blasticidin - a drug that acts as a protein synthesis inhibitor - and mitochondrial respiration inhibition in SWI/SNF-defective cell lines, observed in independent drug and CRISPR screens. Whether this directly results from the loss of a SWI/SNF subunit function which may be involved in protein or mitochondrial metabolism, or whether a SWI/SNF-dependent transcriptional dysregulation of genes involved in such processes secondarily induces these phenotypes, remains to be explored. In this context, the recently described cytoplasmic location of some SWI/SNF subunits is intriguing, and whether some SWI/SNF subunits also have cytoplasmic or mitochondrial functions might require further investigation^34,80^.

In conclusion, our data uncover several previously undescribed targetable synthetic lethal vulnerabilities with defects in SWI/SNF subunits and provide a large muti-omics, drug and CRISPR screen-based resource to the scientific community for further identification of signaling and genetic dependencies in chromatin remodeling-defective cancers.

## Acknowledgments

We thank Prof. Raphaël Margueron (Institut Curie, UMR934, Paris) and Prof. Eric Pasmant (Institut Cochin, Paris) for kindly providing the SMARCB1, BAP-1 or EED KO HAP1 cell lines.

## Funding

JBS was funded by a PhD scholarship from Université Paris-Saclay (France), Institut Gustave Roussy and Institut Curie Foundations.

This work was funded by program grants to SPV:

- Fondation ARC PGA1-RF20190208576
- European Research Council ERC TargetSWitch – 101077864,
- INSERM ATIP-Avenir / La Ligue Contre le Cancer 2018
- Agence Nationale pour la Recherche ANR tremplin-ERC(9)2020
- Gustave Roussy Foundation for the Programme Emergent “Epigénétique” (no grant number).

and program grants to Gustave Roussy:

- INCa-DGOS-Inserm_12551 SIRIC2,
- INCa-DGOS-Inserm-ITMO Cancer_18002 SIRIC EpiCURE.

This work was funded by program grants awarded to AB:

- INSERM ATIP-Avenir/ Plan Cancer (2018-2023)
- Institut Curie Foundation
- INSERM ITMO Cancer « Mathématiques & Informatique contre le cancer » grant (2022-2025)
- INSERM FC3R (2024-2025)

This work was funded by grants awarded to JC:

- BBSCR BB/Y004477/1
- MRC MR/W001276/1

## Author contributions

Conceptualization, SPV, AB; Methodology, JBS, CA, MG, LCD, TIR, MP, CN, DM, RC, MC, JC, PG, EA, EDN, SPV, AB; Software, JBS, PG; Validation, JBS, CA, MG, MC, PG, EDN, SPV, AB; Formal Analysis, JBS, CA, MG, LCD, TIR, MP, PG, SPV, AB; Resources, JC, SPV, AB; Data curation, JBS, CA, MG, LCD, MC, EA, EDN, SPV, AB; Writing – Original Draft, JBS, CA, SPV, AB ; Writing –Review & Editing, JBS, CA, SPV, AB ; Visualization, JBS, CA, MG, RC; Supervision, SPV, AB; Project administration, SPV, AB; Funding Acquisition, SPV, AB ;

## SUPPLEMENTARY MATERIAL

Supplementary material is available at https://github.com/SyspharmaCurie/Bretones-et-al2024

Document S1. Supplementary Figures 1-21

Supplementary file S1. GMT file for pruned Reactome pathway database

Supplementary Table 1. Transcriptomics differential gene expression analysis

Supplementary Table 2. Proteomics differential gene expression analysis

Supplementary Table 3. RIME dataset differential analysis

Supplementary Table 4. Reactome pathways discarded by top-down pruning step.

Supplementary Table 5. Reactome pathways discarded by bottom-up pruning step.

Supplementary Table 6. Transcriptomics enrichment pathway analysis

Supplementary Table 7. Proteomics enrichment pathway analysis

Supplementary Table 8. Unidose Prestwick drug-screening analysis Supplementary

Table 9. Unidose SelleckChem drug-screening analysis

Supplementary Table 10. Multidose Prestwick and SelleckChem drug-screening analyses

Supplementary Table 11. DepMap CRISPR results for HAP1 KO cell lines

Supplementary Table 12. Comparison of CRISPR with Prestwick unidose drug screening based on drug-gene interaction databases

Supplementary Table 13. Comparison of CRISPR with SelleckChem unidose drug screening based on drug-gene interaction databases

## METHODS

### Cell lines

HAP1 parental and KO cell lines for ARID1A, ARID1B, ARID2, PBRM1, SMARCA2, SMARCA4, KMT2C, KMT2D, CREBBP or SETD2 were purchased from Horizon Discovery. HAP-1 KO cell lines for SMARCB1, BAP-1 or EED were obtained from Drs. Raphaël Margueron (Institut Curie, UMR934, Paris) and Eric Pasmant’s teams (Institut Cochin, Paris). NCI-H358 and U2OS cell lines were purchased from ATCC.

SMARCA4 gene knockout was performed in U2OS and NCI-H358 cell lines using a CRISPR/Cas9-based gene editing approach. Cells were targeted using the Edit-R™ CRISPR/Cas9 gene engineering protocol (Horizon), according to the supplier’s instructions. The following sgRNA sequence was used (5′-TTGTCCTGAGGGTACCCTCC-3′) to generate a frameshift deletion in exon 1 of the SMARCA4 gene. Cells were transfected in T25 flasks with sgRNA and Cas9 plasmid, using Lipofectamine 2000 (Thermo Fisher). Several rounds of transfection were performed to obtain an optimal knockout efficiency. SMARCA4 expression was monitored on the transfected pool at each transfection cycle by western blot, and when sufficient depletion of the protein was observed, cells were plated in 96-well plates for clonal isolation using the classical limiting dilution method. Colonies were recovered and profiled for SMARCA4 expression by western blot.

HAP-1 cells were cultured in Iscove’s Modified Dulbecco’s Medium (IMDM, Gibco) supplemented with 10% FBS (Sigma), 1% Penicillin-Streptomycin (Gibco), 1% Sodium Pyruvate (Gibco), 1% Sodium Bicarbonate (Gibco) and 1% Non-Essential Amino Acids (Gibco). U2OS cells were cultured in in high glucose-Dulbecco’s Modified Eagle Medium (DMEM, Gibco) supplemented with 10% FBS (Sigma), 1% Penicillin-Streptomycin (Gibco), 1% Sodium Bicarbonate (Gibco), 1% Non-Essential Amino Acids (Gibco) and 1% HEPES (Gibco). NCI-H358, cells were cultured in Roswell Park Memorial Institute-1640 (RPMI-1640) medium supplemented with 2mM L-glutamine (Gibco), 10% FBS (Sigma) and 1% Penicillin-Streptomycin (Gibco). All cells were grown and maintained at 37°C and 5% CO2. Cells were controlled for mycoplasma-free status using the Venor®GeM Classic Kit (Minerva Biolabs).

### Co-immunoprecipitation SMARCC2 immunoblotting

For co-immunoprecipitation (co-IP), cells were lysed in custom lysis buffer (1% NP-40, 50 mM Tris-HCl, 137 mM NaCl, 10% glycerol, supplemented with 1% Halt™ protease and phosphatase inhibitor cocktail). Lysates were generated on ice and centrifuged for 30 min at 16,900 g before supernatant collection. Lysates containing 300g proteins were incubated O/N at 4°C on a rotating wheel with 50 µL Dynabeads protein G (ThermoFisher, 10004A) and 0.25µg of SMARCC2 antibody (Cell Signaling #12760) or equivalent rabbit polyclonal IgG isotype (Cell Signaling, #2729). Flow-through and IP fractions were collected and subjected to electrophoresis using NuPAGE™ 4-12% Bis-Tris or NuPAGE™ 3-8% Tris-Acetate precast gels (Invitrogen, Carlsbad, CA, USA). After migration, proteins were transferred to a nitrocellulose membrane (GE Healthcare). 3% bovine serum albumin (BSA) in TBS buffer supplemented with 0.1% Tween 20 (TBST 0.1%) was used to block the membrane, at room temperature (RT) for 1 h. Primary antibodies were diluted in 3% BSA in TBST 0.1%, and incubated at 4°C O/N. The next day, the membrane was washed three times with TBST 0.1%, each for 10 min, followed by incubation with horseradish-peroxidase (HRP)-conjugated secondary antibodies at RT for 1 h, in 5% milk in TBST 0.1%. The membrane was washed again three times with TBST 0.1% and incubated with Amersham ECL prime detection reagent (GE Healthcare) or Clarity Max ECL substrate (Biorad). The membrane was then imaged with a BioRad ChemiDoc XRS+ chemiluminescent detection system. Antibodies used: PBRM1 (A301-591A-M) from Bethyl Laboratories (Montgomery, TX, USA); SMARCC1 (#11956S) and SMARCC2 (#12760S) from Cell Signaling Technology (Danvers, MA, USA); ARID1B (ab57461), SMARCA2 (ab15597) from Abcam (Cambridge, UK); ARID1A (sc-32761), ARID2 (sc-166117) SMARCA4 (sc17796), SMARCB1 (sc-166165) from Santa Cruz (Dallas, TX, USA);β-Actin (A1978) from Sigma Aldrich (Gillingham, UK); Goat anti-Rabbit IgG (H+L) Secondary Antibody, HRP, 31460, Thermofisher Scientific; Goat anti-Mouse IgG (H+L) Secondary Antibody, HRP, 31430, Thermofisher Scientific.

### Transcriptome analysis

Transcriptomic analysis was performed on three independent replicates for HAP1 parental cell lines and for all isogenic mutants except SMARCB1, BAP1 and EED. 70-80% confluent cells were harvested, and total RNA was extracted using Rneasy Mini Kit (Qiagen, 74104) with DNAse treatment, according to the manufacturer’s instructions. Every RNA sample was quantified with a Qubit Fluorometer and evaluated for quality controls using 2100 Bioanalyzer (Agilent Technologies). After RNA Integrity Number (RIN) quality control, cDNA libraries were generated using the NEBNext Ultra II RNA Library Prep Kit (NEB #E7775) on Bravo Liquid Handler (Agilent). Subsequent indexed RNA sequencing of cDNA libraries with paired-end reads was performed according to the standard Illumina protocol using NovaSeq 6000 S2 system, with a target of 100Gb (X million reads) per sample. Pre-processing of reads included quality controls with FASTQC and adapter trimming with TrimGalore using the nf-core RNA-seq pipeline. Following alignment to GRCh38 with STAR, the quality of RNAseq data was evaluated with RseQC (BAM stat, junction saturation, RPKM saturation, read duplication, Inner distance). FeatureCounts was used to quantify expression relative to the transcriptome.

### Proteome analysis

#### Samples preparation

Proteomic analysis was performed on 3 independent replicates of HAP1 parental cell lines and for all isogenic mutants except CREBBP, KMT2C, KMT2D, and SETD2, for which 2 biological replicates were processed. Cells were harvested at 70% confluence and pellets were dissolved in 150µL lysis buffer containing 1% sodium deoxycholate (SDC), 100mM triethylammonium bicarbonate (TEAB), 10% isopropanol, 50mM NaCl and Halt protease and phosphatase inhibitor cocktail (100X) (Thermo Fisher, #78442). Pellets were pulsed with probe sonication for 15 sec (on ice), followed by boiling at 90°C for 5 min and re-sonication for 5 sec. Protein concentration was measured with the Coomassie Plus Bradford Protein Assay (Pierce) according to the manufacturer’s instructions. Protein aliquots of 100 μg were reduced with 5 mM tris-2-carboxyethyl phosphine (TCEP) for 1 h at 60 °C and alkylated with 10 mM Iodoacetamide (IAA) for 30 min. Proteins were finally digested with trypsin (Pierce) at 75 ng/μL O/N. The peptides were labelled with the TMT-11plex reagents (Thermo Fisher) according to the manufacturer’s instructions. Peptides were fractionated with the XBridge C18 column (2.1 x 150 mm, 3.5 μm, Waters) on a Dionex Ultimate 3000 HPLC system at high pH. Mobile phase A was 0.1% ammonium hydroxide and mobile phase B was acetonitrile, 0.1% ammonium hydroxide. The TMT labelled peptide mixture was fractionated using a multi-step gradient elution at 0.2 mL/min. The separation method was: for 5 minutes isocratic at 5% B, for 35 min gradient to 35% B, gradient to 80% B in 5 min, isocratic for 5 minutes, and re-equilibration to 5% B. Fractions were collected every 30 sec and vacuum dried.

#### LC-MS/MS

On-line LC-MS/MS analysis was performed on the Dionex Ultimate 3000 system coupled with the Orbitrap Lumos Mass Spectrometer (Thermo Scientific). Peptide fractions were reconstituted in 40 μL 0.1% formic acid and 10 μL were first loaded and desalted on a PepMap C18 Nanotrap trapping column (100 μm × 2 cm C18, 5 μm, 100 Å) at 10 μL/min flow rate. The samples were then analyzed with the Acclaim PepMap RSLC (75 μm × 50 cm, 2 μm, 100 Å) C18 capillary column at 45 °C. Mobile phase A was 0.1% formic acid and mobile phase B was 80% acetonitrile, 0.1% formic acid. The gradient method at flow rate 300 nL/min was: for 90 min gradient from 5%-38% B, for 10 min up to 95% B, for 5 min isocratic at 95% B, re-equilibration to 5% B in 5 min, for 10 min isocratic at 10% B. Precursor ions within 375-1,500 m/z were selected at mass resolution of 120 k in top speed mode (3 sec cycle) and were isolated for CID fragmentation with quadrupole isolation width 0.7 Th, collision energy 35% and max IT 50 ms. MS3 spectra were obtained with further HCD fragmentation of the top 5 most abundant CID fragments isolated with Synchronous Precursor Selection (SPS). Collision energy was applied at 65% with 105 ms IT and 50 k resolution. Targeted precursors were dynamically excluded for further activation for 45 seconds with 7 ppm mass tolerance.

#### MS data analysis

The mass spectra were submitted to SequestHT for database search in Proteome Discoverer 2.2 (Thermo Scientific) using reviewed UniProt human protein entries. The precursor mass tolerance was 20 ppm and the fragment ion mass tolerance was 0.5 Da for fully tryptic peptides. TMT6plex at N-terminus/K and Carbamidomethyl at C were selected as static modifications. Dynamic modifications were oxidation of M, and deamidation of N/Q. Peptide confidence was estimated with the Percolator node, and peptides were filtered for q-value to be used for protein quantification.

Firstly, a differential expression analysis was performed to generate lists of genes/proteins deregulated in a mutant as compared to the wild-type cell line. Differential expression analyses were performed using limma-voom^81,82^ after Trimmed Mean of M values (TMM) normalization^83^. For differential analysis, a gene was assumed to be dysregulated if its p-adj was below 0.05. DE was performed for HAP1 transcriptomics and proteomics datasets and for transcriptomics data from Schick et al. To compare both transcriptomics studies, only common mutant and WT HAP1 cell lines were selected. Whole transcriptome comparison was done by taking all common genes between datasets in each mutant and computing the Pearson correlation on the list of log_2_FCs. Similarly, the correlation between HAP1 transcriptome and the proteome differential expression analysis was assessed by Pearson’s test performed on gene log_2_(FC) values^84^. For Gene Set Analysis, differentially expressed genes were ranked using the absolute value of the t-statistic computed by limma.

Next, Gene Set Analysis was conducted to find biological pathways overrepresented in each differentially expressed gene list. Pathway definitions were based on a pruned version of the Reactome database V82^85^ (see next paragraph) and specified in the form of a Gene Matrix Transposed (GMT) file.

Regarding pathway enrichment, an optimized pipeline was developed from the combined study of three methods: ^(1)^ Overrepresentation Analysis (ORA, Gprofiler2 R package^86^), pre-ranked Gene Set Enrichment Analysis (GSEA, fgsea R package^87,88^), and ROntoTools^27^. ORA is a first-generation method that uses only differentially expressed genes as input and consists of a hypergeometric test to compare to a background set of human genes. GSEA is a second-generation method (Functional Class Sorting) and it employs a Kolmorogov-Smirnoff running sum using the whole gene list ranked by a statistic, in this study, the t-test statistic given by limma. ROntoTools is a third-generation method (topology-based) that computes a pathway perturbation score using a pathway graph representation where the nodes (proteins) are weighted based on the p-adj of differential expression analysis using the formula -log_10_(p-adj/max(p-adj)). For all three methods, Benjamini-Hochberg multiple testing correction was applied to compute p-adj values for each pathway which served to rank them.

A benchmark of 13 methods, based on 4 different metrics, concluded on the superiority of ROntoTools, followed by GSEA, and highlighted a considerable difference in performance between both algorithms and the rest of the methods^28^. ORA was included in that study and was found to be the worst-performing method among all evaluated ones. Nonetheless, a Nature Protocol study published in 2019 indicated that ORA was the most utilized method across all Gene Set Analysis algorithms and was included here as a negative control^25^. Of note, in their study, Nguyen et al. used GSEA with phenotype permutations, which requires at least seven biological replicates per condition, a condition rarely met in high-throughput data analysis^89^. Having at most three replicates per condition, we had to use the pre-ranked version of GSEA. Being aware that it has an expected higher false positive rate as compared to GSEA algorithm with phenotype permutations, we lowered the p-adj threshold from 0.25 to 0.05, as proposed by Subramanian in the GSEA documentation (https://www.gsea-msigdb.org/gsea/doc/GSEAUserGuideFrame.html). Next, given the rather low sensitivity (i.e.: true positive rate) reported for enrichment methods, we explored the combination of existing algorithms as associating several methods may potentially increase the number of discoveries. Furthermore, ROntoTools can only be run on pathways for which the topology is documented, and does not report the sign of pathway dysregulation, as GSEA does with the Normalized Enrichment Scores (NES). To reduce the number of false positives when combining methods, the results of each algorithm were merged by computing the geometric mean of pathway ranks^90^.

### RIME

#### Immunoprecipitation

Cells were fixed with 0.1mM DSP (dithiobis[succinimidylpropionate], Thermo Fisher Scientific, 22585) for 30 minutes at RT, washed and lysed using the following lysis buffer: 50 mM Tris-HCl, 137 mM NaCl, 10% glycerol, 1% NP40 and protease inhibitor cocktail (Roche, 11836153001). Lysates were generated on ice and incubated with 1u/mL Benzonase Nuclease (Sigma, E1014-5KU) and 2mM MgCl2 for 30 minutes on a rotating wheel, then centrifuged 15 min at 16,900 g before supernatant collection. Lysates were incubated with 1µg SMARCC2 antibody (Cell Signaling, #12760) or rabbit polyclonal IgG isotype (Cell Signaling, #2729) on a rotating wheel for 2 hours at 4°C and 1 hour more incubation with 1.5mg Dynabeads protein G (Thermo Fisher Scientific, 10004A). IP fractions were washed four times with lysis buffer, then washed twice with 50 mM ammonium bicarbonate, before trypsin digestion (1µg Trypsin Sequencing Grade, Merck, 11418475001) O/N at 37°C with shaking. Peptides were purified by filtering throught Millipore Multiscreen HTS plate before mass spec analysis.

#### LC-MS/MS Analysis

On-line LC-MS/MS analysis was performed on an Orbitrap Fusion Lumos hybrid mass spectrometer coupled with an Ultimate 3000 RSLCnano UPLC system. Samples were first loaded and desalted on a PepMap C18 nano trap (100 µm i.d. x 20 mm, 100 Å, 5µ), then peptides were separated on a PepMap C18 column (75 µm i.d. x 500 mm, 2 µm) over a linear gradient of 4–32% CH3CN/0.1% FA in 90 min, cycle time at 120 min at a flow rate of 300 nl/min. All instruments and HPLC columns were from Thermo Fisher. The MS acquisition used the standard DDA method with Top Speed 3s cycle time. Briefly, the Orbitrap full MS survey scan was m/z 375 – 1500 with resolution 120,000 at m/z 200, with AGC (Automatic Gain Control) set at 40,000 and maximum injection time at 50 sec. Multiply charged ions (z = 2 – 5) with intensity above 8,000 counts were fragmented in HCD (higher collision dissociation) cell at 30% collision energy, and the isolation window was set at 1.6 Th. The fragment ions were detected in the ion trap with AGC at 10,000 and maximum injection time at 50 ms. The dynamic exclusion time was set at 30 s with ±10 ppm.

#### MS data analysis

Raw data were analysed using MaxQuant v1.6.1.43^91^. Protein identification was performed by database searches against the SwissProt human database (August 2019) and quantification used the MaxLFQ algorithm^91^ and the iBAQ measure^92^. The following search and quantification parameters were used: Trypsin was set as digestion mode with a maximum of two missed cleavages allowed. The main search peptide tolerance was set to 20 ppm, and MS/MS match tolerance set to 0.5 Da. Acetylation at the N-terminus, oxidation of methionine, and deamidation of asparagine or glutamine were set as variable modifications. Peptide and protein identifications were set at 1% FDR.

Data analysis was performed with the Perseus software (v1.6.2.3). Contaminants, reverse hits,proteins identified only by site, and proteins identified in only one of the biological replicates for each KO line were removed before further analysis. For each sample, the iBAQ values were normalised by scaling to the iBAQ of the bait protein (SMARCC2) in the corresponding sample. Normalised iBAQ values were transformed and missing values were imputed from a normal distribution representing lowest abundance values to enable statistical analysis. Welch’s t-tests were performed for each KO line against the wild type. Averaged scaled relative iBAQs were hierarchically clustered using Pearson correlation in Phantasus^93^.

### Cell line clustering based on gene expression or pathway enrichment

Cell line clustering was done for the transcriptome and proteome based on the t-statistic of differential expression analysis. Cell line clustering for transcriptome and proteome enrichment results was based on the geometric mean of pathway rank. Both analyses were done with the Ward hierarchical clustering algorithm and the Euclidean distance. In both cases, but especially for the genes and proteins, our data was subject to the “Curse of dimensionality” problem, as we tried to cluster ∼10 cell lines using thousands of dimensions (∼17000 in transcriptome, ∼ 8000 in proteome, ∼ 400 in enrichment). For this reason, the Principal Components Analysis (PCA) dimensionality reduction method was applied before clustering ^94^. In every analysis, choosing the number of principal components that explained between 85 and 90% of the variance seemed to yield the best results. Choosing a lower or higher number of components did not allow a biologically meaningful classification. In the former case, PCA did not retain enough signal or variance explained to conduct a proper clustering. In the latter case, PCA likely kept components that include noise or that are redundant with other components, therefore worsening the clustering.

### Reduction of Reactome pathway database

Reactome database was used for pathway definition given its very comprehensive nature (Reactome V85, 2629 pathways, 29 pathway categories)^85^. A two-step pruning algorithm was developed to obtain a minimum set of Reactome pathways of interest. First, a top-down pruning was performed taking advantage of the tree structure of Reactome and keeping only pathways with lengths between minimal and maximal values to be specified. It is accepted that pathways smaller than 10 genes may not be informative enough and that pathways larger than 500 genes may be too general and should be discarded^25^. Our procedure starts at the top of the tree, from the pathway corresponding to the whole Reactome category, and goes down. In every iteration, the current pathway (parent) is checked for correct size. If its size is not within bounds, it is ruled out. If its size is correct, its children are checked as well for correct size and eliminated if their size is out of bounds. Then, the union of the genes of the remaining children is taken and compared to the parent gene set. If at least 1 gene is lost in the children’s union set, the parent is kept, as its gene set cannot be reproduced by merging the child pathways. On the opposite, if the parent is included in its children, it can be discarded. This procedure is repeated iteratively until reaching every leaf of the tree (i.e.: pathways that do not have any children). It is done independently for each of the 28 categories in Reactome, each regarded as a different pathway tree. Secondly, a bottom-up pruning step was conducted for pathways remaining from step 1. A given pathway is compared to the rest and, if it includes other smaller pathways in their entirety, those pathways are eliminated. For the present study aiming at identifying novel drug targets in the context of SWI/SNF deficiencies, we discarded nine irrelevant Reactome pathway categories which were: *drug ADME*, *Hemostasis*, *Digestion and absorption*, *Reproduction*, *Sensory Perception*, *Neuronal System*, *Developmental Biology*, *Muscle contraction*, and *Disease*.

### Unidose and multidose drug screening

High-throughput drug screening assays were performed in the HAP1 WT and 11 mutant cell lines (*SMARCA2, SMARCA4, ARID1A, ARID1B, ARID2, PBRM1, CREBBP, KMT2D, KMT2C, SETD2, EED* and *BAP1*-KO). Cells were cultured in Iscove’s Modified Dulbecco’s Medium (IMDM - Gibco Life Technologies - 31980048), supplemented with 10% fetal bovine serum (Gibco Life Technologies – 10270106), 1% of Penicillin/Streptomycin (Gibco Life Technologies - 15140122), Sodium Pyruvate (Gibco Life Technologies - 11360039), Sodium Bicarbonate 7.5% solution (Gibco Life Technologies - 25080094) and Non-Essential Amino Acids Solution (Gibco Life Technologies - 11140035), in a 37°C incubator with 5% CO2. These cells were cultured in flasks and expanded as 2D monolayers using the same media as described above. For cell passages, cells were washed with PBS and detached with trypsin-EDTA (Gibco Life Technologies - 25300054) for 5 minutes at 37°C. All cell lines were tested mycoplasma-free by the MycoAlert™ Mycoplasma Detection Kit (Lonza).

The unidose dataset comprised 1,200 chemicals from the Prestwick Chemical Library and 186 compounds from the SelleckChem Epigenetic Library. Cells were counted using a T4 Cellometer (Nexcelom) to obtain the desired number of cells for the screen and seeded in 384-well plates (ViewPlate-384 Black, Perkin Elmer) using MultiDrop combi (Thermo Fisher Scientific), in 40µL of media. For the Prestwick library, cell line densities were empirically determined: 250 cells per well for *SMARCA2* and *KMT2-*KO; 300 cells per well for *ARID1A, ARID2, PBRM1, SMARCA4, KMT2C*-KO, and HAP1; 400 cells per well for ARID1B, *BAP1* and *EED-*KO; 800 cells per well for *CREBBP* and *SETD2*-KO. For the SelleckChem library, cell line densities were as follows: 100 cells per well for all cell lines except for the *SMARCA4*- (150 cells per well) and *KMT2C-KO* (200 cells per well). Twenty-four hours after cell seeding, compounds or control DMSO-vehicle were transferred to the cell plates using the MultiChannel Arm™ 384 (MCA 384) (TECAN), to a final concentration of 10µM and 0.5% of DMSO. Cells were then incubated for 5 days before cell viability assessment for both libraries. For the Epigenetic Library, cells were incubated with the chemical compounds for two exposure durations, 5 days and 11 days, before cell viability assessment. CellTiter-Glo 2.0 Assay kit (Promega Inc., Madison, USA) was used to determine the cell viability in compound-treated wells. The CellTiter-Glo 2.0 reagent was equilibrated to room temperature for 30 min before use. A volume of 25µL per well of CellTiter-Glo 2.0 reagent was robotically added using MultiDrop combi (Thermo Fisher Scientific). The contents were mixed for 2 minutes at 300 rpm on an orbital shaker (Titramax 100, Dutscher) and plates were further incubated for 10 min at room temperature to stabilize luminescent signals. Units of luminescent signal generated by a thermos-stable luciferase are proportional to the amount of ATP present in viable cells. Luminescence was recorded using a CLARIOStar (BMG Labtech) (gain = 3600). In the unidose study, the survival fraction (SF) was calculated as the ratio between raw luminescence in a treated well and the median luminescence of DMSO-treated wells of the same replicate. Hits were subsequently identified based on three criteria: 1) efficacy (SF for both replicates < 70%), 2) specificity (WT-vs-mutant difference of SF for both replicates > 30%), and 3) non-toxicity (SF for both replicates > 70% in WT HAP1).

From this unidose study, the 55 compounds most cytotoxic for the mutant cell lines while not harming the WT cells, were selected for a multi-dose assay (see details below), together with 56 epigenetic drugs chosen from the SelleckChem Epigenetic Library. All compounds were received already diluted in Dimethyl Sulfoxide (DMSO) as 10 mM stock solution and reformatted in-house into 384-well source plates for automatic dose-response robotic screening. DMSO was added in the remaining wells as internal plate solvent controls. Cells were seeded with the same density as in the unidose experiments, as described above. Twenty-four hours after cell seeding, cell plates were treated with compounds and vehicle (0.5% DMSO maximum) titrated in 8-point, three-fold dilutions starting at a concentration of 10 µM. For the Prestwick library, cell viability was assessed after 5 days of compound incubation (6 days after cell seeding). For the epigenetic drugs, half of the media was removed, fresh media was added after a 120h incubation period, and the assay plates were incubated at 37°C. Cell viability was assessed after 144h (12 days after cell seeding). For each condition, cell viability was assessed by measuring cellular ATP through luminescence using the Cell-titer Glo technology as described previously. The screen was performed at the same early cell passages (±1) for the three biological replicates when the cells had been passaged four times after thawing from liquid nitrogen. The survival fraction (SF) was computed as the ratio between raw luminescence in a treated well and the median luminescence of wells containing no cells from the same plaque. Compounds were subsequently sorted by molecular targets, and manually selected for multi-dose (dose-response) revalidation, based on the frequency of targets identified in a specific class, the # of compounds of that specific class within the Prestwick library, and the subsequent potential for repurposing or clinical drug development.

For the multi-dose medium-throughput screen, a log-logistic curve was fitted to each concentration-SF dataset using the drc R package and further used for compound selection. Firstly, any chemical killing more than 30% of the HAP1 WT cell population at any concentration was considered too toxic and was eliminated for further analyses. Next, compounds showing increasing or flat log-logistic curves for the mutant cell lines were considered ineffective and were discarded (NoEffect function of the drc package, p<0.05, and visual inspection)^95^. Similarly, chemicals for which the fitted curve did not drop under 50% survival at the highest concentration (10 µM) were considered as not cytotoxic enough and were eliminated. In summary, the compounds of interest were defined as the ones reaching an IC_50_ between 0 and 10 µM in a mutant cell line while not reaching an IC_30_ in HAP1 WT cells.

### Assessing pathway targetability from drug screening data

Drug-gene interaction databases contain qualitative drug-gene interactions based on curated scientific evidence. Three databases were used to map working chemicals from the unidose and multi-dose analyses to their interacting genes: drug-gene interaction database (DGIdb)^52^, Comparative Toxicogenomics Database (CTD)(Davis et al., 2021),and DrugBank^54^. An extra manual mapping was added for chemicals unable to map to any of the three databases. In case unmapped chemicals remain after the manual curation, PubChem Bioassays *in vitro* data was used to add interacting genes to such drugs. For the multi-dose analysis, only cytotoxic chemicals with an IC_50_ < 5 µM were selected for mapping to avoid the non-specific effects of drugs working only at very high concentrations.

In the previous step, every cytotoxic chemical in a mutant cell line was successfully mapped to the set of interacting genes based on drug-gene interaction databases. Such gene list was used as input to an Overrepresentation Analysis (ORA) against the pruned Reactome database (the same version used to conduct enrichment for transcriptomics and proteomics data). The test results in a list of pathways ranked in increasing order based on the p-adj. For every pathway list corresponding to every cytotoxic chemical in a mutant, only significant pathways were selected (p-adj < 0.05). A heatmap representing the pathways targeted in at least one SWI/SNF KO cell line and targeted based on the proteome enrichment was built by merging ORA results of cytotoxic drugs of the unidose and multidose analyses (Fig. 5D).

### Estimating gene targetability using the CRISPR Cancer Dependency Map (DepMap) project

The DepMap project is a public database containing hundreds of cancer cell line models profiled for genomic information and sensitivity to genetic and small molecule perturbations^23^. Here, we used the DepMap transcriptomics (RNA-seq) and CRISPR datasets available in 1100 cell lines. The CRISPR technology was used to knock out approximately 17000 genes and compute a CRISPR dependency score for every gene. The more negative this score is, the more deleterious the elimination of that gene is for the cellular phenotype^96^.

An algorithm was developed to predict synthetic lethal interactions between HAP1 chromatin remodeling mutant genes (E.g.: *SMARCA2*) and any other gene from the Reactome pruned database (called here CRISPR gene). The method has 2 steps. (1) First, the definition of high and low mutant expression cell populations. We obtain the expression distribution of the HAP1 mutated gene across all cell lines in the database that contain measured expression for this gene. Then, the first and last 10% quantiles were selected as the low and high expression groups, respectively. (2) The second step consists of testing a displacement between CRISPR score distributions of the high and low mutant gene expression cell populations. Because of the non-normality of the CRISPR score distribution, the nonparametric Wilcox rank sum test was used to test for a difference in CRISPR gene scores between both cell line groups. If the score is significantly more negative in the low expression group (p-adj < 0.05), the algorithm would predict a possible synthetic lethality between the mutant and the CRISPR gene (I.e.: knocking out the CRISPR gene on a cell line already deficient for the mutant gene would lead to cell death). The algorithm was run for every mutant independently and for all genes in the pruned Reactome database. The results are shown in form of a volcano plot indicating the Wilcox test displacement and the -log_10_(p-adj) (Supplementary Fig. 6 to 18). A Benjamini-Hochberg multiple testing correction was applied. Genes with a p-adj < 1e-3 and a Wilcox displacement estimate > 0.15 were considered as “hits” (enough evidence of a synthetic lethal interaction). Hits with a p-adj < 1e-4 were regarded as “super hits” (strong evidence of a synthetic lethal interaction). For each mutant cell line, the list of CRISPR gene hits was used as input for ORA against the pruned Reactome database to obtain the set of pathways enriched in the hit list. Significant pathways (p-adj < 0.05) were named as targetable pathways for a particular mutant cell line, as they are overrepresented in the hit gene list.

### Revalidation proliferation assays

The p300 inhibitor C646, the CBP inhibitor CPI-637, and the mitochondrial respiration complex V inhibitor Oligomycin A were purchased from Selleck Chemicals. The mitochondrial respiration complex III inhibitor Antimycin A was kindly gifted by Dr. Catherine Brenner’s team (Institut Gustave Roussy, UMR9018, France). Short-term survival assays were performed in 96-well plates. Exponentially-growing cells were plated in triplicates at a density of 150-1000 cells per well. Drug/vehicle was added 24 h after seeding and cells were continuously exposed to the drug for 7 days, after which cell viability was estimated using CellTiter-Glo® Luminescent Cell Viability Assay (Promega, Madison, WI, USA) on a Spark® Multimode Microplate Reader (Tecan, Switzerland). Long-term survival assays were performed in 6-well plates. Exponentially growing cells were plated in triplicates at a density of 150-300 cells/well. Drug/vehicle was added 24 h after seeding and cells were exposed to the drug for at least 10 days, with replenishment every 2 days by freshly made dilutions of drug or vehicle. Cells were fixed and stained with 0.5% crystal violet in 25% methanol for 30 min and washed three times with deionized water. Colonies of > 50 cells were counted manually. Survival fractions (SF) were calculated compared to the DMSO-treated control and dose-response curves were generated using Prism 10 (GraphPad Software®) after Log transformation of the drug concentration. Two-way ANOVA testing the effect of WT/KO conditions and of varied concentrations of drug exposure on SF and post hoc Sidak’s tests were performed to assess the significance of the drug efficacy.

## REFERENCES

1. Hanahan, D. Hallmarks of Cancer: New Dimensions. Cancer Discov 12, 31–46 (2022).

2. Shain, A. H. & Pollack, J. R. The Spectrum of SWI/SNF Mutations, Ubiquitous in Human Cancers. PLoS One 8, (2013).

3. Kadoch, C. et al. Proteomic and bioinformatic analysis of mammalian SWI/SNF complexes identifies extensive roles in human malignancy. Nat Genet 45, 592–601 (2013).

4. Savas, S. & Skardasi, G. The SWI/SNF complex subunit genes: Their functions, variations, and links to risk and survival outcomes in human cancers. Crit Rev Oncol Hematol 123, 114–131 (2018).

5. Chabanon, R. M., Morel, D. & Postel-Vinay, S. Exploiting epigenetic vulnerabilities in solid tumors: Novel therapeutic opportunities in the treatment of SWI/SNF-defective cancers. Semin Cancer Biol 61, 180–198 (2020).

6. Mittal, P. & Roberts, C. W. M. The SWI/SNF complex in cancer — biology, biomarkers and therapy. Nature Reviews Clinical Oncology 2020 17:7 17, 435–448 (2020).

7. Schick, S. et al. Systematic characterization of BAF mutations provides insights into intracomplex synthetic lethalities in human cancers. Nature Genetics 2019 51:9 51, 1399–1410 (2019).

8. Mashtalir, N. et al. Modular Organization and Assembly of SWI/SNF Family Chromatin Remodeling Complexes. Cell 175, 1272–1288.e20 (2018).

9. Morel, D., Almouzni, G., Soria, J. C. & Postel-Vinay, S. Targeting chromatin defects in selected solid tumors based on oncogene addiction, synthetic lethality and epigenetic antagonism. Annals of Oncology 28, 254–269 (2017).

10. Helming, K. C. et al. ARID1B is a specific vulnerability in ARID1A-mutant cancers. Nat Med 20, 251–254 (2014).

11. Hoffman, G. R. et al. Functional epigenetics approach identifies BRM/SMARCA2 as a critical synthetic lethal target in BRG1-deficient cancers. Proceedings of the National Academy of Sciences 111, 3128–3133 (2014).

12. Gounder, M. et al. Tazemetostat in advanced epithelioid sarcoma with loss of INI1/SMARCB1: an international, open-label, phase 2 basket study. Lancet Oncol 21, 1423–1432 (2020).

13. Italiano, A. et al. Tazemetostat, an EZH2 inhibitor, in relapsed or refractory B-cell non-Hodgkin lymphoma and advanced solid tumours: a first-in-human, open-label, phase 1 study. Lancet Oncol 19, 649–659 (2018).

14. Chabanon, R. M., Morel, D. & Postel-Vinay, S. Exploiting epigenetic vulnerabilities in solid tumors: Novel therapeutic opportunities in the treatment of SWI/SNF-defective cancers. Semin Cancer Biol 61, 180–198 (2020).

15. Ngo, C. & Postel-Vinay, S. Immunotherapy for SMARCB1-Deficient Sarcomas: Current Evidence and Future Developments. Biomedicines 10, (2022).

16. MorphoSys receives U.S. FDA fast track designation for tulmimetostat in endometrial cancer. News release. MorphoSys AG. September 12, 2023. . *Accessed from:* https://tinyurl.com/2p9rfvfb https://tinyurl.com/2p9rfvfb.

17. Kurashima, K., et al. SMARCA4 deficiency-associated heterochromatin induces intrinsic DNA replication stress and susceptibility to ATR inhibition in lung adenocarcinoma. NAR Cancer 2, zcaa005 (2020).

18. Smith-Roe, S. L. et al. SWI/SNF complexes are required for full activation of the DNA-damage response. Oncotarget 6, 732–745 (2015).

19. Williamson, C. T. et al. ATR inhibitors as a synthetic lethal therapy for tumours deficient in ARID1A. Nat Commun 7, 13837 (2016).

20. Shen, J. et al. ARID1A Deficiency Impairs the DNA Damage Checkpoint and Sensitizes Cells to PARP Inhibitors. Cancer Discov 5, 752–767 (2015).

21. Wang, X. et al. BRD9 defines a SWI/SNF sub-complex and constitutes a specific vulnerability in malignant rhabdoid tumors. Nat Commun 10, (2019).

22. Sabnis, R. W. Novel Compounds for Targeted Degradation of BRD9 and Their Use for Treating Cancer. ACS Med Chem Lett 13, (2022).

23. Tsherniak, A. et al. Defining a Cancer Dependency Map. Cell 170, 564–576.e16 (2017).

24. Dempster, J. M., et al. Extracting Biological Insights from the Project Achilles Genome-Scale CRISPR Screens in Cancer Cell Lines. bioRxiv 720243 (2019) doi:10.1101/720243.

25. Reimand, J. et al. Pathway enrichment analysis and visualization of omics data using g:Profiler, GSEA, Cytoscape and EnrichmentMap. Nat Protoc 14, 482–517 (2019).

26. Simillion, C., Liechti, R., Lischer, H. E. L., Ioannidis, V. & Bruggmann, R. Avoiding the pitfalls of gene set enrichment analysis with SetRank. BMC Bioinformatics 18, 151 (2017).

27. Voichita, C., Donato, M. & Draghici, S. Incorporating gene significance in the impact analysis of signaling pathways. Proceedings - 2012 11th International Conference on Machine Learning and Applications, ICMLA 2012 1, 126–131 (2012).

28. Nguyen, T. M., Shafi, A., Nguyen, T. & Draghici, S. Identifying significantly impacted pathways: A comprehensive review and assessment. Genome Biol 20, 1–15 (2019).

29. Nakayama, R. T. et al. SMARCB1 is required for widespread BAF complex–mediated activation of enhancers and bivalent promoters. Nat Genet 49, 1613–1623 (2017).

30. Roumeliotis, T. I. et al. Genomic Determinants of Protein Abundance Variation in Colorectal Cancer Cells. Cell Rep 20, 2201–2214 (2017).

31. Batsché, E., Yaniv, M. & Muchardt, C. The human SWI/SNF subunit Brm is a regulator of alternative splicing. Nat Struct Mol Biol 13, 22–29 (2006).

32. Gañez-Zapater, A. et al. The SWI/SNF subunit BRG1 affects alternative splicing by changing RNA binding factor interactions with nascent RNA. Molecular Genetics and Genomics 297, 463–484 (2022).

33. Sahu, R. K. et al. SWI/SNF chromatin remodelling complex contributes to clearance of cytoplasmic protein aggregates and regulates unfolded protein response in Saccharomyces cerevisiae. FEBS J 287, 3024–3041 (2020).

34. Ulicna, L. et al. The Interaction of SWI/SNF with the Ribosome Regulates Translation and Confers Sensitivity to Translation Pathway Inhibitors in Cancers with Complex Perturbations. Cancer Res 82, 2829–2837 (2022).

35. Yan, Z. et al. PBAF chromatin-remodeling complex requires a novel specificity subunit, BAF200, to regulate expression of selective interferon-responsive genes. Genes Dev 19, 1662–1667 (2005).

36. Zhou, M., Yuan, J., Deng, Y., Fan, X. & Shen, J. Emerging role of SWI/SNF complex deficiency as a target of immune checkpoint blockade in human cancers. Oncogenesis 10, 3 (2021).

37. Li, J. et al. Epigenetic driver mutations in ARID1A shape cancer immune phenotype and immunotherapy. Journal of Clinical Investigation 130, 2712–2726 (2020).

38. Maxwell, M. B. et al. ARID1A suppresses R-loop-mediated STING-type I interferon pathway activation of anti-tumor immunity. Cell 187, 3390–3408.e19 (2024).

39. Bakr, A. et al. ARID1A regulates DNA repair through chromatin organization and its deficiency triggers DNA damage-mediated anti-tumor immune response. Nucleic Acids Res 52, 5698–5719 (2024).

40. Miao, D. et al. Genomic correlates of response to immune checkpoint therapies in clear cell renal cell carcinoma. Science (1979) 359, 801–806 (2018).

41. Yu, N.-K. et al. Interactome analysis illustrates diverse gene regulatory processes associated with LIN28A in human iPS cell-derived neural progenitor cells. iScience 24, 103321 (2021).

42. Lissanu Deribe, Y., et al. Mutations in the SWI/SNF complex induce a targetable dependence on oxidative phosphorylation in lung cancer. Nat Med 24, 1047–1057 (2018).

43. Zhang, L. et al. Identifying Cell Cycle Modulators That Selectively Target ARID1A Deficiency Using High-Throughput Image-Based Screening. SLAS Discovery 22, 813–826 (2017).

44. Wilson, M. R. et al. ARID1A Mutations Promote P300-Dependent Endometrial Invasion through Super-Enhancer Hyperacetylation. Cell Rep 33, (2020).

45. Nie, M. et al. Genome-wide CRISPR screens reveal synthetic lethal interaction between CREBBP and EP300 in diffuse large B-cell lymphoma. Cell Death Dis 12, 419 (2021).

46. Varoni, E. M., Lo Faro, A. F., Sharifi-Rad, J. & Iriti, M. Anticancer Molecular Mechanisms of Resveratrol. Front Nutr 3, (2016).

47. Rauf, A. et al. Resveratrol as an anti-cancer agent: A review. Crit Rev Food Sci Nutr 58, 1428–1447 (2018).

48. Gatchalian, J. et al. A non-canonical BRD9-containing BAF chromatin remodeling complex regulates naive pluripotency in mouse embryonic stem cells. Nat Commun 9, 5139 (2018).

49. Obach, R. S. POTENT INHIBITION OF HUMAN LIVER ALDEHYDE OXIDASE BY RALOXIFENE. Drug Metabolism and Disposition 32, 89–97 (2004).

50. Shorstova, T. et al. SWI/SNF-Compromised Cancers Are Susceptible to Bromodomain Inhibitors. Cancer Res 79, 2761–2774 (2019).

51. Campagne, A. et al. BAP1 complex promotes transcription by opposing PRC1-mediated H2A ubiquitylation. Nat Commun 10, 348 (2019).

52. Freshour, S. L. et al. Integration of the Drug–Gene Interaction Database (DGIdb 4.0) with open crowdsource efforts. Nucleic Acids Res 49, D1144–D1151 (2021).

53. Davis, A. P. et al. Comparative Toxicogenomics Database (CTD): Update 2021. Nucleic Acids Res 49, D1138–D1143 (2021).

54. Wishart, D. S. et al. DrugBank 5.0: a major update to the DrugBank database for 2018. Nucleic Acids Res 46, D1074–D1082 (2018).

55. Kim, S. et al. PubChem in 2021: new data content and improved web interfaces. Nucleic Acids Res 49, D1388–D1395 (2021).

56. DepMap Broad. DepMap 24Q2 Public. https://depmap.org/portal (2024).

57. Wappett, M. et al. SynLeGG: analysis and visualization of multiomics data for discovery of cancer ‘Achilles Heels’ and gene function relationships. Nucleic Acids Res 49, W613–W618 (2021).

58. Joubert, F. & Puff, N. Mitochondrial Cristae Architecture and Functions: Lessons from Minimal Model Systems. Membranes (Basel) 11, (2021).

59. Taylor, A. M. et al. Fragment-Based Discovery of a Selective and Cell-Active Benzodiazepinone CBP/EP300 Bromodomain Inhibitor (CPI-637). ACS Med Chem Lett 7, 531–536 (2016).

60. Sasaki, E. et al. Structure and assembly of scalable porous protein cages. Nat Commun 8, 14663 (2017).

61. Blümli, S. et al. Acute depletion of the ARID1A subunit of SWI/SNF complexes reveals distinct pathways for activation and repression of transcription. Cell Rep 37, 109943 (2021).

62. Geng, H. et al. Leveraging synthetic lethality to uncover potential therapeutic target in gastric cancer. Cancer Gene Ther 31, 334–348 (2024).

63. Schenk, S. et al. Combined transcriptome and proteome profiling reveals specific molecular brain signatures for sex, maturation and circalunar clock phase. Elife 8, e41556 (2019).

64. Casas-Vila, N. et al. The developmental proteome of Drosophila melanogaster. Genome Res 27, 1273–1285 (2017).

65. Grün, D. et al. Conservation of mRNA and protein expression during development of C.elegans. Cell Rep 6, 565–577 (2014).

66. Tops, B. B. J., Gauci, S., Heck, A. J. R. & Krijgsveld, J. Worms from Venus and Mars: Proteomics profiling of sexual differences in Caenorhabditis elegans using in vivo15N isotope labeling. J Proteome Res 9, 341–351 (2010).

67. Ghazalpour, A. et al. Comparative Analysis of Proteome and Transcriptome Variation in Mouse. PLoS Genet 7, 1001393 (2011).

68. Gry, M. et al. Correlations between RNA and protein expression profiles in 23 human cell lines. BMC Genomics 10, 1–14 (2009).

69. Shankavaram, U. T. et al. Transcript and protein expression profiles of the NCI-60 cancer cell panel: an integromic microarray study. Mol Cancer Ther 6, 820–832 (2007).

70. Mandal, J., Mandal, P., Wang, T.-L. & Shih, I.-M. Treating ARID1A mutated cancers by harnessing synthetic lethality and DNA damage response. J Biomed Sci 29, 71 (2022).

71. Watanabe, R. et al. SWI/SNF Factors Required for Cellular Resistance to DNA Damage Include ARID1A and ARID1B and Show Interdependent Protein Stability. Cancer Res 74, 2465–2475 (2014).

72. Oba, A. et al. ARID2 modulates DNA damage response in human hepatocellular carcinoma cells. J Hepatol 66, 942–951 (2017).

73. Moreno, T. et al. ARID2 deficiency promotes tumor progression and is associated with higher sensitivity to chemotherapy in lung cancer. Oncogene 40, 2923–2935 (2021).

74. Chabanon, R. M. et al. PBRM1 Deficiency Confers Synthetic Lethality to DNA Repair Inhibitors in Cancer. Cancer Res 81, 2888–2902 (2021).

75. Gu, D. et al. PBRM1 Deficiency Sensitizes Renal Cancer Cells to DNMT Inhibitor 5-Fluoro-2’-Deoxycytidine. Front Oncol 12, (2022).

76. Feng, H. et al. PBAF loss leads to DNA damage-induced inflammatory signaling through defective G2/M checkpoint maintenance. Genes Dev 36, 790–806 (2022).

77. Rother, M. B. & van Attikum, H. DNA repair goes hip-hop: SMARCA and CHD chromatin remodellers join the break dance. Philos Trans R Soc Lond B Biol Sci 372, (2017).

78. Ribeiro-Silva, C. et al. DNA damage sensitivity of SWI/SNF-deficient cells depends on TFIIH subunit p62/GTF2H1. Nat Commun 9, 4067 (2018).

79. Kim, K. H. & Roberts, C. W. M. Mechanisms by which SMARCB1 loss drives rhabdoid tumor growth. Cancer Genet 207, 365–372 (2014).

80. Yu, S. et al. SWI/SNF interacts with cleavage and polyadenylation factors and facilitates pre-mRNA 3′ end processing. Nucleic Acids Res 46, 8557–8573 (2018).

81. Law, C. W., Chen, Y., Shi, W. & Smyth, G. K. voom: precision weights unlock linear model analysis tools for RNA-seq read counts. Genome Biol 15, R29 (2014).

82. Smyth, G. K. Linear Models and Empirical Bayes Methods for Assessing Differential Expression in Microarray Experiments. 3, (2004).

83. Robinson, M. D. & Oshlack, A. A scaling normalization method for differential expression analysis of RNA-seq data. Genome Biol 11, R25 (2010).

84. Cañas, R. A. et al. Transcriptome-wide analysis supports environmental adaptations of two Pinus pinaster populations from contrasting habitats. BMC Genomics 16, 909 (2015).

85. Jassal, B. et al. The reactome pathway knowledgebase. Nucleic Acids Res 48, D498–D503 (2020).

86. Kolberg, L., Raudvere, U., Kuzmin, I., Vilo, J. & Peterson, H. gprofiler2 – an R package for gene list functional enrichment analysis and namespace conversion toolset g:Profiler [version 2; peer review: 2 approved] . F1000Res 9, (2020).

87. Korotkevich, G., Sukhov, V. & Sergushichev, A. Fast gene set enrichment analysis. bioRxiv 060012 (2016) doi:10.1101/060012.

88. Subramanian, A. et al. Gene set enrichment analysis: a knowledge-based approach for interpreting genome-wide expression profiles. Proc Natl Acad Sci U S A 102, 15545–15550 (2005).

89. Yoon, S., Kim, S.-Y. & Nam, D. Improving Gene-Set Enrichment Analysis of RNA-Seq Data with Small Replicates. PLoS One 11, e0165919- (2016).

90. Italiani, P. et al. Profiling the Course of Resolving vs. Persistent Inflammation in Human Monocytes: The Role of IL-1 Family Molecules. Front Immunol 11, (2020).

91. Cox, J. & Mann, M. MaxQuant enables high peptide identification rates, individualized p.p.b.-range mass accuracies and proteome-wide protein quantification. Nat Biotechnol 26, 1367–1372 (2008).

92. Schwanhäusser, B. et al. Global quantification of mammalian gene expression control. Nature 473, 337–342 (2011).

93. Kleverov, M. et al. Phantasus, a web application for visual and interactive gene expression analysis. Elife 13, (2024).

94. Murtagh, F. & Legendre, P. Ward’s Hierarchical Agglomerative Clustering Method: Which Algorithms Implement Ward’s Criterion? J Classif 31, 274–295 (2014).

95. Ritz, C., Baty, F., Streibig, J. C. & Gerhard, D. Dose-Response Analysis Using R. PLoS One 10, e0146021 (2015).

96. Dempster, J. M. et al. Chronos: a cell population dynamics model of CRISPR experiments that improves inference of gene fitness effects. Genome Biol 22, 343 (2021).

