## Supplementary Figures for "Integrated multiomic profiling reveals SWI/SNF subunit-specific pathway alterations and targetable vulnerabilities"

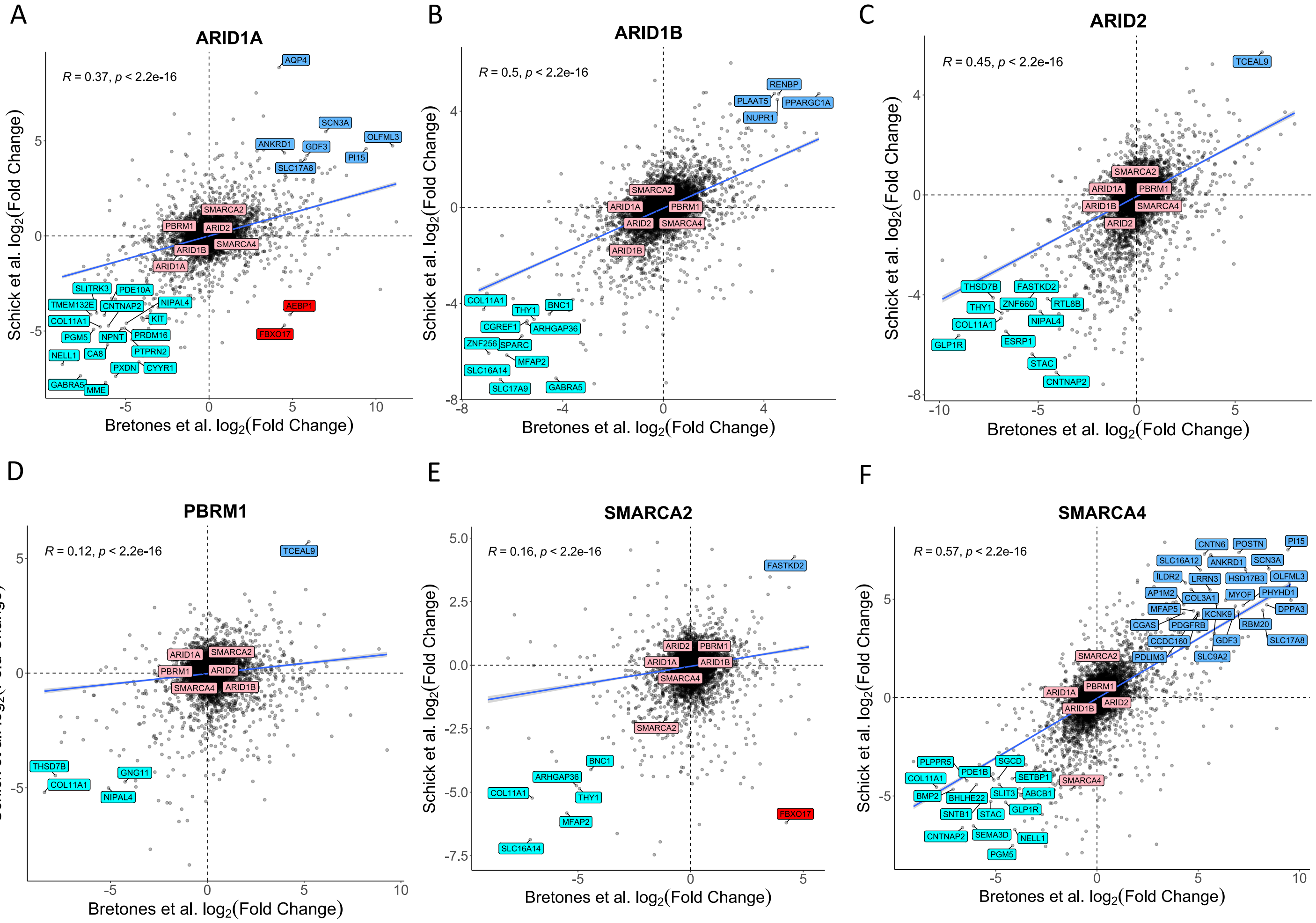


**Supplementary Fig. 1. Comparison of the transcriptome in HAP1 KO cell lines between our study and Schick et al.**[7] .Comparison of the transcriptome differential expression in *ARID1A-*, *ARID1B-*, *ARID2-*, *PBRM1-*, *SMARCA2-*, and *SMARCA4*-KO cell lines. The color code indicates the quadrant where the most differentially expressed genes ($|log_2FC| > 4$) lie (blue = upregulation in both datasets, cyan = downregulation in both datasets, red = upregulation in our study and downregulation in Schick's, beige = upregulation in Schick's and downregulation in our study, pink = SWI/SNF related genes). The blue line represents the Pearson correlation line, with indicated parameters.

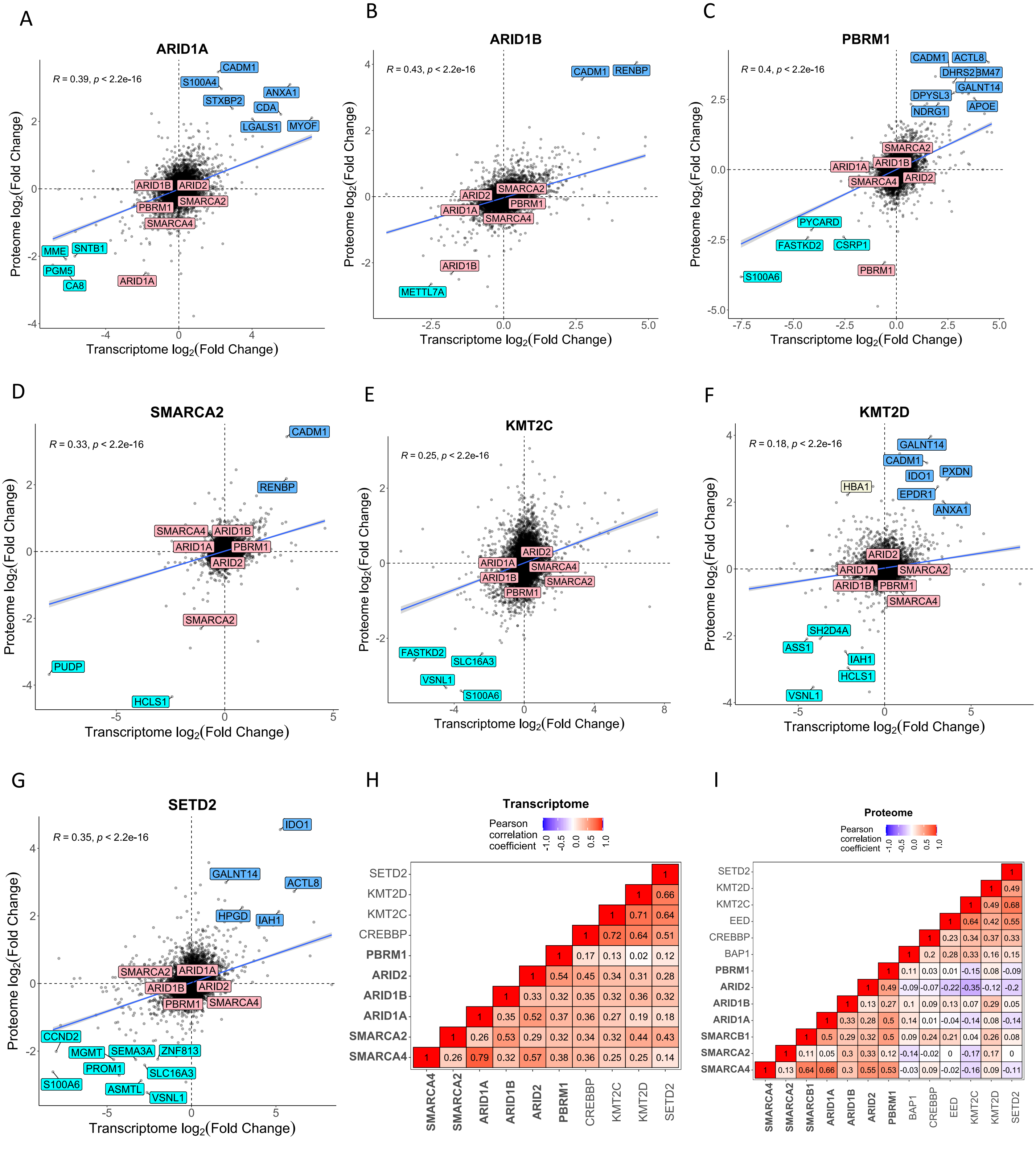


**Supplementary Fig. 2. Correlation of transcriptome and proteome analysis across the HAP1 isogenic panel. A-G)** Comparison of transcriptome and proteome differential expression in the *ARID1B*-, *PBRM1*-, *SMARCA4*-, *SMARCA2*-, *KMT2C*-, *KMT2D*-, and *SETD2*-KO cell lines. The color code indicates the quadrant where the most differentially expressed genes ($|\log_2 FC| > 2$) lie (blue = upregulation in both transcriptome and proteome, cyan = downregulation in both transcriptome and proteome, beige = upregulation in transcriptome and downregulation in proteome, pink = SWI/SNF related genes). The blue line represents the Pearson correlation line, with indicated parameters. **H,I)** Pearson correlation between isogenic cell lines of the transcriptome (H) and the proteome (I).

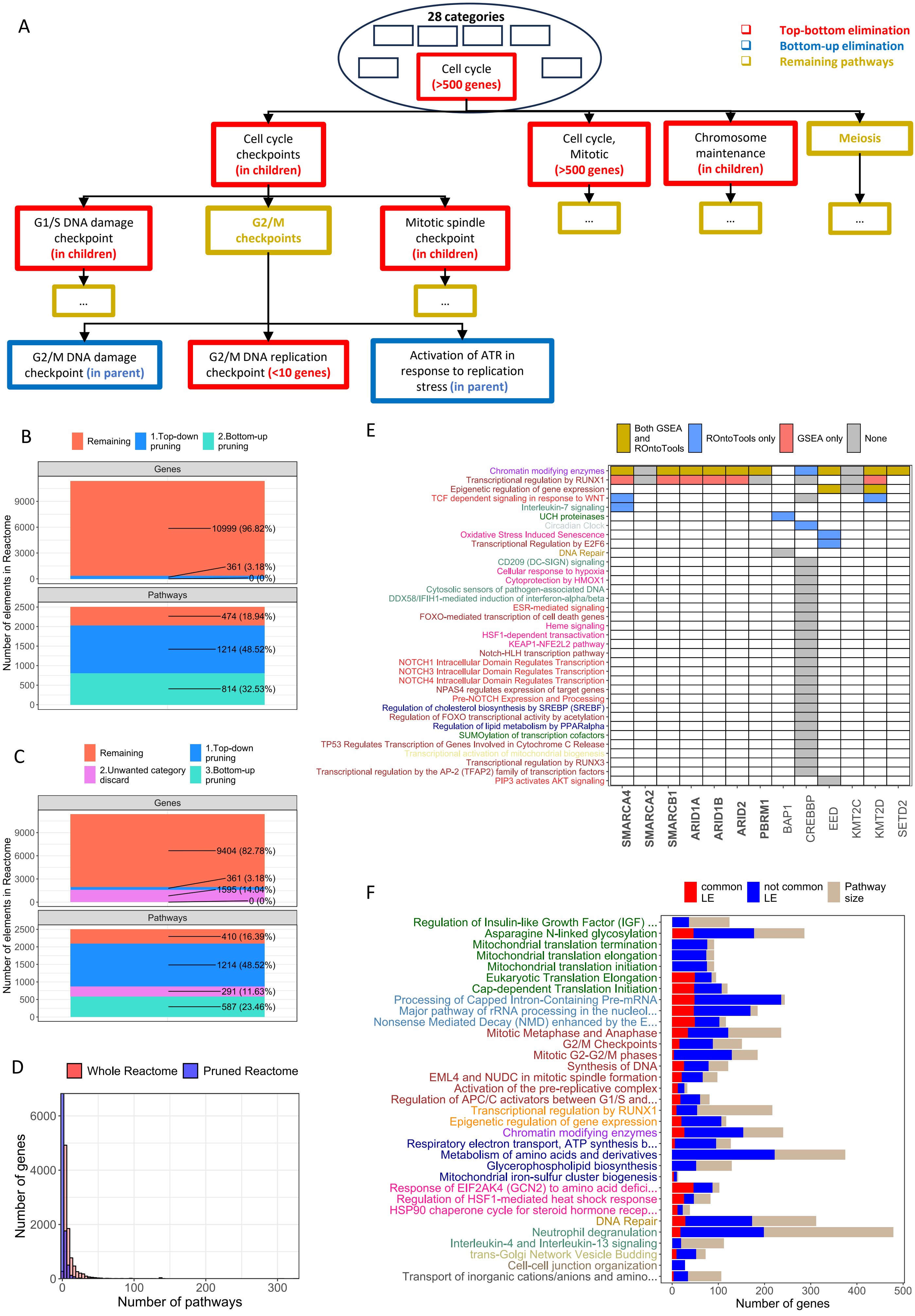

A
28 categories
Cell cycle
(>500 genes)
Top-bottom elimination
Bottom-up elimination
Remaining pathways
Cell cycle checkpoints (in children)
Cell cycle, Mitotic (>500 genes)
Chromosome maintenance (in children)
Meiosis
G1/S DNA damage checkpoint (in children)
G2/M checkpoints
Mitotic spindle checkpoint (in children)
...
G2/M DNA damage checkpoint (in parent)
G2/M DNA replication checkpoint (<10 genes)
Activation of ATR in response to replication stress (in parent)
B
Remaining
1.Top-down pruning
2.Bottom-up pruning
Genes
Pathways
Number of elements in Reactome
10999 (96.82%)
361 (3.18%)
0 (0%)
474 (18.94%)
1214 (48.52%)
814 (32.53%)
C
Remaining
1.Top-down pruning
2.Unwanted category discard
3.Bottom-up pruning
Genes
Pathways
Number of elements in Reactome
9404 (82.78%)
361 (3.18%)
1595 (14.04%)
0 (0%)
410 (16.39%)
1214 (48.52%)
291 (11.63%)
587 (23.46%)
D
Whole Reactome
Pruned Reactome
Number of genes
Number of pathways
E
Both GSEA and ROntoTools
ROntoTools only
GSEA only
None
Chromatin modifying enzymes
Transcriptional regulation by RUNX1
Epigenetic regulation of gene expression
TCF dependent signaling in response to WNT
Interleukin-7 signaling
UCH proteinases
Circadian Clock
Oxidative Stress Induced Senescence
Transcriptional Regulation by E2F6
DNA Repair
CD209 (DC-SIGN) signaling
Cellular response to hypoxia
Cytoprotection by HMOX1
Cytosolic sensors of pathogen-associated DNA
DDX58/IFIH1-mediated induction of interferon-alpha/beta
ESR-mediated signaling
FOXO-mediated transcription of cell death genes
Heme signaling
HSF1-dependent transactivation
KEAP1-NFE2L2 pathway
Notch-HLH transcription pathway
NOTCH1 Intracellular Domain Regulates Transcription
NOTCH3 Intracellular Domain Regulates Transcription
NOTCH4 Intracellular Domain Regulates Transcription
NPAS4 regulates expression of target genes
Pre-NOTCH Expression and Processing
Regulation of cholesterol biosynthesis by SREBP (SREBF)
Regulation of FOXO transcriptional activity by acetylation
Regulation of lipid metabolism by PPARalpha
SUMOylation of transcription cofactors
TP53 Regulates Transcription of Genes Involved in Cytochrome C Release
Transcriptional activation of mitochondrial biogenesis
Transcriptional regulation by RUNX3
Transcriptional regulation by the AP-2 (TFAP2) family of transcription factors
PIP3 activates AKT signaling
SMARCA4
SMARCA2
SMARCB1
ARID1A
ARID1B
ARID2
PBRM1
BAP1
CREBBP
EED
KMT2C
KMT2D
SETD2
F
common LE
not common LE
Pathway size
Regulation of Insulin-like Growth Factor (IGF) ...
Asparagine N-linked glycosylation
Mitochondrial translation termination
Mitochondrial translation elongation
Mitochondrial translation initiation
Eukaryotic Translation Elongation
Cap-dependent Translation Initiation
Processing of Capped Intron-Containing Pre-mRNA
Major pathway of rRNA processing in the nucleol...
Nonsense Mediated Decay (NMD) enhanced by the E...
Mitotic Metaphase and Anaphase
G2/M Checkpoints
Mitotic G2-G2/M phases
Synthesis of DNA
EML4 and NUDC in mitotic spindle formation
Activation of the pre-replicative complex
Regulation of APC/C activators between G1/S and...
Transcriptional regulation by RUNX1
Epigenetic regulation of gene expression
Chromatin modifying enzymes
Respiratory electron transport, ATP synthesis b...
Metabolism of amino acids and derivatives
Glycerophospholipid biosynthesis
Mitochondrial iron-sulfur cluster biogenesis
Response of EIF2AK4 (GCN2) to amino acid defici...
Regulation of HSF1-mediated heat shock response
HSP90 chaperone cycle for steroid hormone recep...
DNA Repair
Neutrophil degranulation
Interleukin-4 and Interleukin-13 signaling
trans-Golgi Network Vesicle Budding
Cell-cell junction organization
Transport of inorganic cations/anions and amino...
Number of genes

**Supplementary Fig. 3. <u>Development of a novel Gene Set Analysis pipeline</u>. A)** Scheme of the pruning algorithm developed for the Reactome database, including: (1) a top-down and (2) a bottom-up pruning step. The former is applied to every category in Reactome independently, starting from the largest pathway that gives name to that category. Maximum and minimum pathway size are defined as hyperparameter (in this example, min_size = 10 and max_size = 500 genes). Iteratively, every pathway is checked for correct size, and for not being entirely included in its children. If the pathway size is not within the limits OR if the pathway is included in its children, it is discarded (red rectangle). The remaining pathways after running the pruning algorithm have golden rectangles. After the top-down pruning, the bottom-up approach is applied to the remaining pathways, regardless of their category. **B)** Number of remaining genes and pathways after running both pruning steps of the methods. The total number of genes and pathways correspond to the original Reactome database. **C)** Number of remaining genes and pathways after applying the pruning method and discarding unwanted categories for this project (see Methods). Category exclusion were performed after the top-down step and before the bottom-up pruning. **D)** Distribution of gene redundancy defined as the number of pathways where each gene appears, either in the original Reactome database or in the pruned version. In the whole database, the mean number of pathways per gene is 10.4, whereas it is only 2.79 in the pruned version. Similarly, in the whole database the median number of pathways per gene is 6, but only 1 in the pruned version. **E)** Proteome enrichment results for pathways containing the KO SWI/SNF gene in the current cell line. **F)** Leading edges (LE) of SWI/SNF mutants for the Top10 pathways represented in Fig. 4G, either in common (red) or unique to any cell line (blue). The total pathway size is represented in beige.

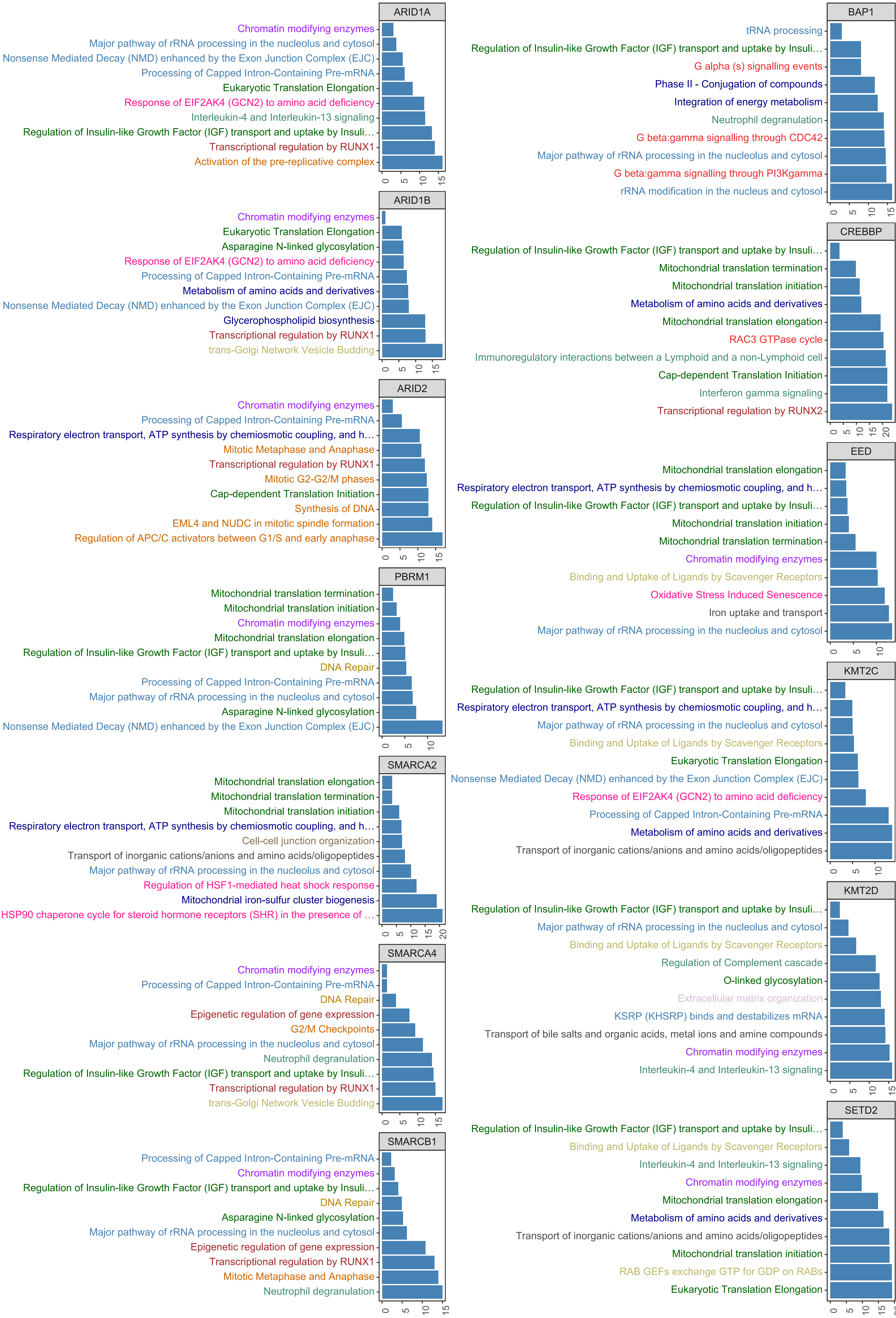
ARID1A
Chromatin modifying enzymes
Major pathway of rRNA processing in the nucleolus and cytosol
Nonsense Mediated Decay (NMD) enhanced by the Exon Junction Complex (EJC)
Processing of Capped Intron-Containing Pre-mRNA
Eukaryotic Translation Elongation
Response of EIF2AK4 (GCN2) to amino acid deficiency
Interleukin-4 and Interleukin-13 signaling
Regulation of Insulin-like Growth Factor (IGF) transport and uptake by Insuli…
Transcriptional regulation by RUNX1
Activation of the pre-replicative complex
0 5 10 15
ARID1B
Chromatin modifying enzymes
Eukaryotic Translation Elongation
Asparagine N-linked glycosylation
Response of EIF2AK4 (GCN2) to amino acid deficiency
Processing of Capped Intron-Containing Pre-mRNA
Metabolism of amino acids and derivatives
Nonsense Mediated Decay (NMD) enhanced by the Exon Junction Complex (EJC)
Glycerophospholipid biosynthesis
Transcriptional regulation by RUNX1
trans-Golgi Network Vesicle Budding
0 5 10 15
ARID2
Chromatin modifying enzymes
Processing of Capped Intron-Containing Pre-mRNA
Respiratory electron transport, ATP synthesis by chemiosmotic coupling, and h…
Mitotic Metaphase and Anaphase
Transcriptional regulation by RUNX1
Mitotic G2-G2/M phases
Cap-dependent Translation Initiation
Synthesis of DNA
EML4 and NUDC in mitotic spindle formation
Regulation of APC/C activators between G1/S and early anaphase
0 5 10 15
PBRM1
Mitochondrial translation termination
Mitochondrial translation initiation
Chromatin modifying enzymes
Mitochondrial translation elongation
Regulation of Insulin-like Growth Factor (IGF) transport and uptake by Insuli…
DNA Repair
Processing of Capped Intron-Containing Pre-mRNA
Major pathway of rRNA processing in the nucleolus and cytosol
Asparagine N-linked glycosylation
Nonsense Mediated Decay (NMD) enhanced by the Exon Junction Complex (EJC)
0 5 10
SMARCA2
Mitochondrial translation elongation
Mitochondrial translation termination
Mitochondrial translation initiation
Respiratory electron transport, ATP synthesis by chemiosmotic coupling, and h…
Cell-cell junction organization
Transport of inorganic cations/anions and amino acids/oligopeptides
Major pathway of rRNA processing in the nucleolus and cytosol
Regulation of HSF1-mediated heat shock response
Mitochondrial iron-sulfur cluster biogenesis
HSP90 chaperone cycle for steroid hormone receptors (SHR) in the presence of …
0 5 10 15 20
SMARCA4
Chromatin modifying enzymes
Processing of Capped Intron-Containing Pre-mRNA
DNA Repair
Epigenetic regulation of gene expression
G2/M Checkpoints
Major pathway of rRNA processing in the nucleolus and cytosol
Neutrophil degranulation
Regulation of Insulin-like Growth Factor (IGF) transport and uptake by Insuli…
Transcriptional regulation by RUNX1
trans-Golgi Network Vesicle Budding
0 5 10 15
SMARCB1
Processing of Capped Intron-Containing Pre-mRNA
Chromatin modifying enzymes
Regulation of Insulin-like Growth Factor (IGF) transport and uptake by Insuli…
DNA Repair
Asparagine N-linked glycosylation
Major pathway of rRNA processing in the nucleolus and cytosol
Epigenetic regulation of gene expression
Transcriptional regulation by RUNX1
Mitotic Metaphase and Anaphase
Neutrophil degranulation
0 5 10 15
BAP1
tRNA processing
Regulation of Insulin-like Growth Factor (IGF) transport and uptake by Insuli…
G alpha (s) signalling events
Phase II - Conjugation of compounds
Integration of energy metabolism
Neutrophil degranulation
G beta:gamma signalling through CDC42
Major pathway of rRNA processing in the nucleolus and cytosol
G beta:gamma signalling through PI3Kgamma
rRNA modification in the nucleus and cytosol
0 5 10 15
CREBBP
Regulation of Insulin-like Growth Factor (IGF) transport and uptake by Insuli…
Mitochondrial translation termination
Mitochondrial translation initiation
Metabolism of amino acids and derivatives
Mitochondrial translation elongation
RAC3 GTPase cycle
Immunoregulatory interactions between a Lymphoid and a non-Lymphoid cell
Cap-dependent Translation Initiation
Interferon gamma signaling
Transcriptional regulation by RUNX2
0 5 10 15 20
EED
Mitochondrial translation elongation
Respiratory electron transport, ATP synthesis by chemiosmotic coupling, and h…
Regulation of Insulin-like Growth Factor (IGF) transport and uptake by Insuli…
Mitochondrial translation initiation
Mitochondrial translation termination
Chromatin modifying enzymes
Binding and Uptake of Ligands by Scavenger Receptors
Oxidative Stress Induced Senescence
Iron uptake and transport
Major pathway of rRNA processing in the nucleolus and cytosol
0 5 10
KMT2C
Regulation of Insulin-like Growth Factor (IGF) transport and uptake by Insuli…
Respiratory electron transport, ATP synthesis by chemiosmotic coupling, and h…
Major pathway of rRNA processing in the nucleolus and cytosol
Binding and Uptake of Ligands by Scavenger Receptors
Eukaryotic Translation Elongation
Nonsense Mediated Decay (NMD) enhanced by the Exon Junction Complex (EJC)
Response of EIF2AK4 (GCN2) to amino acid deficiency
Processing of Capped Intron-Containing Pre-mRNA
Metabolism of amino acids and derivatives
Transport of inorganic cations/anions and amino acids/oligopeptides
0 5 10
KMT2D
Regulation of Insulin-like Growth Factor (IGF) transport and uptake by Insuli…
Major pathway of rRNA processing in the nucleolus and cytosol
Binding and Uptake of Ligands by Scavenger Receptors
Regulation of Complement cascade
O-linked glycosylation
Extracellular matrix organization
KSRP (KHSRP) binds and destabilizes mRNA
Transport of bile salts and organic acids, metal ions and amine compounds
Chromatin modifying enzymes
Interleukin-4 and Interleukin-13 signaling
0 5 10 15
SETD2
Regulation of Insulin-like Growth Factor (IGF) transport and uptake by Insuli…
Binding and Uptake of Ligands by Scavenger Receptors
Interleukin-4 and Interleukin-13 signaling
Chromatin modifying enzymes
Mitochondrial translation elongation
Metabolism of amino acids and derivatives
Transport of inorganic cations/anions and amino acids/oligopeptides
Mitochondrial translation initiation
RAB GEFs exchange GTP for GDP on RABs
Eukaryotic Translation Elongation
0 5 10 15

**Supplementary Fig. 4.** **<u>Top10 enriched pathways in the proteomics of SWI/SNF (first column) and non SWI/SNF (second column) mutants in proteomics</u>**. Pathways are colored based on the Reactome category they belong to (see Fig. 3B for legend).

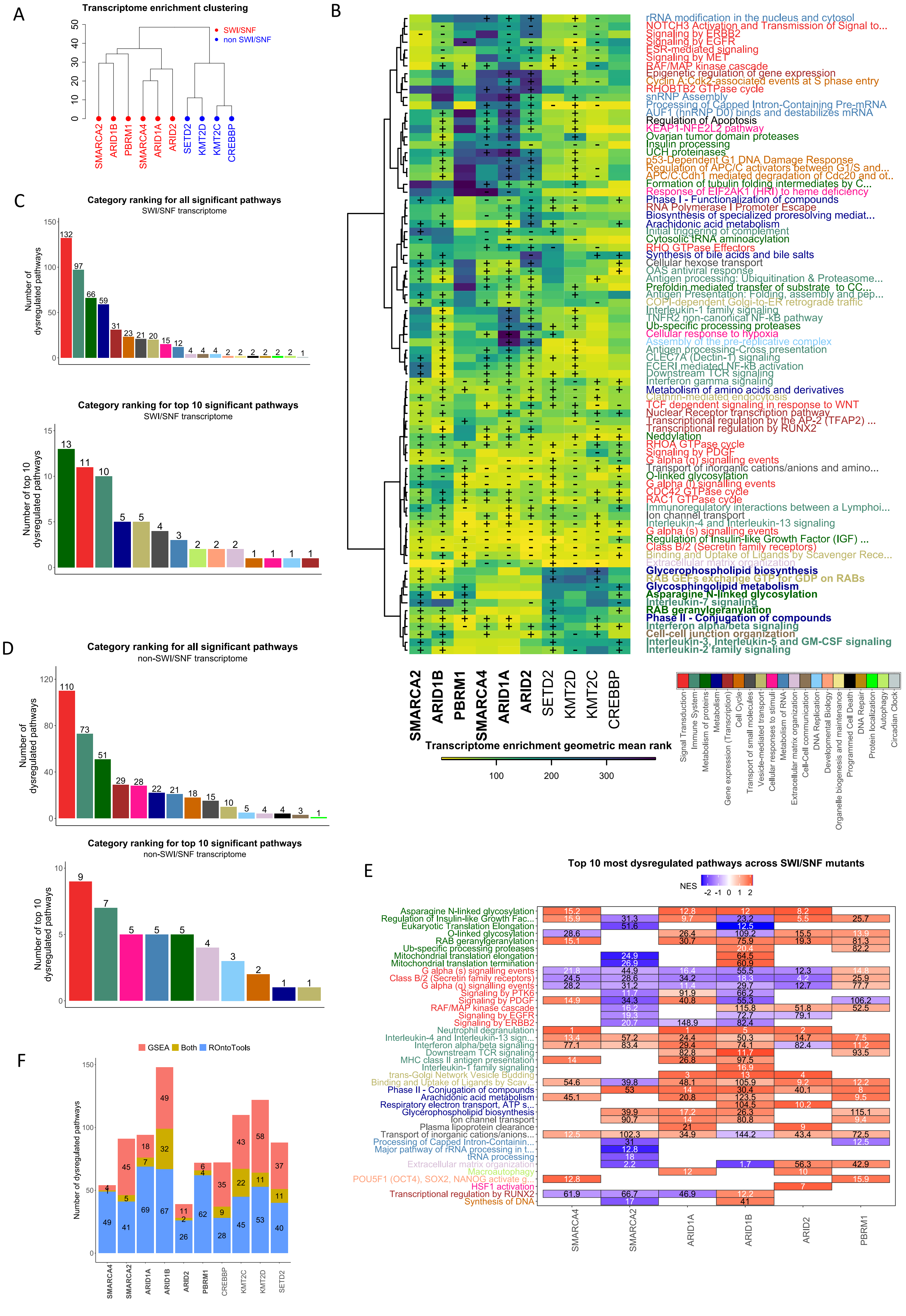
A
Transcriptome enrichment clustering
SWI/SNF
non SWI/SNF
SMARCA2
ARID1B
PBRM1
SMARCA4
ARID1A
ARID2
SETD2
KMT2D
KMT2C
CREBBP
B
rRNA modification in the nucleus and cytosol
NOTCH3 Activation and Transmission of Signal to...
Signaling by ERBB2
Signaling by EGFR
ESR-mediated signaling
Signaling by MET
RAF/MAP kinase cascade
Epigenetic regulation of gene expression
Cyclin A:Cdk2-associated events at S phase entry
RHOBTB2 GTPase cycle
snRNP Assembly
Processing of Capped Intron-Containing Pre-mRNA
AUF1 (hnRNP D0) binds and destabilizes mRNA
Regulation of Apoptosis
KEAP1-NFE2L2 pathway
Ovarian tumor domain proteases
Insulin processing
UCH proteinases
p53-Dependent G1 DNA Damage Response
Regulation of APC/C activators between G1/S and...
APC/C:Cdh1 mediated degradation of Cdc20 and ot...
Formation of tubulin folding intermediates by C...
Response of EIF2AK1 (HRI) to heme deficiency
Phase I - Functionalization of compounds
RNA Polymerase I Promoter Escape
Biosynthesis of specialized proresolving mediat...
Arachidonic acid metabolism
Initial triggering of complement
Cytosolic tRNA aminoacylation
RHO GTPase Effectors
Synthesis of bile acids and bile salts
Cellular hexose transport
OAS antiviral response
Antigen processing: Ubiquitination & Proteasome...
Prefoldin mediated transfer of substrate to CC...
Antigen Presentation: Folding, assembly and pep...
COPI-dependent Golgi-to-ER retrograde traffic
Interleukin-1 family signaling
TNFR2 non-canonical NF-kB pathway
Ub-specific processing proteases
Cellular response to hypoxia
Assembly of the pre-replicative complex
Antigen processing-Cross presentation
CLEC7A (Dectin-1) signaling
FCERI mediated NF-kB activation
Downstream TCR signaling
Interferon gamma signaling
Metabolism of amino acids and derivatives
Clathrin-mediated endocytosis
TCF dependent signaling in response to WNT
Nuclear Receptor transcription pathway
Transcriptional regulation by the AP-2 (TFAP2) ...
Transcriptional regulation by RUNX2
Neddylation
RHOA GTPase cycle
Signaling by PDGF
G alpha (q) signalling events
Transport of inorganic cations/anions and amino...
O-linked glycosylation
G alpha (i) signalling events
CDC42 GTPase cycle
RAC1 GTPase cycle
Immunoregulatory interactions between a Lymphoi...
Ion channel transport
Interleukin-4 and Interleukin-13 signaling
G alpha (s) signalling events
Regulation of Insulin-like Growth Factor (IGF) ...
Class B/2 (Secretin family receptors)
Binding and Uptake of Ligands by Scavenger Rece...
Extracellular matrix organization
Glycerophospholipid biosynthesis
RAB GEFs exchange GTP for GDP on RABs
Glycosphingolipid metabolism
Asparagine N-linked glycosylation
Interleukin-7 signaling
RAB geranylgeranylation
Phase II - Conjugation of compounds
Interferon alpha/beta signaling
Cell-cell junction organization
Interleukin-3, Interleukin-5 and GM-CSF signaling
Interleukin-2 family signaling
Transcriptome enrichment geometric mean rank
100
200
300
Signal Transduction
Immune System
Metabolism of proteins
Metabolism
Gene expression (Transcription)
Cell Cycle
Transport of small molecules
Vesicle-mediated transport
Cellular responses to stimuli
Metabolism of RNA
Extracellular matrix organization
Cell-Cell communication
DNA Replication
Developmental Biology
Organelle biogenesis and maintenance
Programmed Cell Death
DNA Repair
Protein localization
Autophagy
Circadian Clock
C
Category ranking for all significant pathways
SWI/SNF transcriptome
Number of dysregulated pathways
132 97 66 59 31 23 21 20 15 12 4 4 4 2 2 2 2 2 2 1
Category ranking for top 10 significant pathways
SWI/SNF transcriptome
Number of top 10 dysregulated pathways
13 11 10 5 5 4 3 2 2 2 1 1 1 1
D
Category ranking for all significant pathways
non-SWI/SNF transcriptome
Number of dysregulated pathways
110 73 51 29 28 22 21 18 15 10 5 4 4 3 1
Category ranking for top 10 significant pathways
non-SWI/SNF transcriptome
Number of top 10 dysregulated pathways
9 7 5 5 5 4 3 2 1 1
E
Top 10 most dysregulated pathways across SWI/SNF mutants
NES
-2 -1 0 1 2
Asparagine N-linked glycosylation
Regulation of Insulin-like Growth Fac...
Eukaryotic Translation Elongation
O-linked glycosylation
RAB geranylgeranylation
Ub-specific processing proteases
Mitochondrial translation elongation
Mitochondrial translation termination
G alpha (s) signalling events
Class B/2 (Secretin family receptors)
G alpha (q) signalling events
Signaling by PTK6
Signaling by PDGF
RAF/MAP kinase cascade
Signaling by EGFR
Signaling by ERBB2
Neutrophil degranulation
Interleukin-4 and Interleukin-13 sign...
Interferon alpha/beta signaling
Downstream TCR signaling
MHC class II antigen presentation
Interleukin-1 family signaling
trans-Golgi Network Vesicle Budding
Binding and Uptake of Ligands by Scav...
Phase II - Conjugation of compounds
Arachidonic acid metabolism
Respiratory electron transport, ATP s...
Glycerophospholipid biosynthesis
Ion channel transport
Plasma lipoprotein clearance
Transport of inorganic cations/anions...
Processing of Capped Intron-Containin...
Major pathway of rRNA processing in t...
tRNA processing
Extracellular matrix organization
Macroautophagy
POU5F1 (OCT4), SOX2, NANOG activate g...
HSF1 activation
Transcriptional regulation by RUNX2
Synthesis of DNA
SMARCA4
SMARCA2
ARID1A
ARID1B
ARID2
PBRM1
F
GSEA
Both
ROntoTools
Number of dysregulated pathways
SMARCA4
SMARCA2
ARID1A
ARID1B
ARID2
PBRM1
CREBBP
KMT2C
KMT2D
SETD2

**Supplementary Fig. 5.** **<u>Results of transcriptome enrichment for HAP1 KO cell lines</u>.** **A)** Hierarchical clustering on the PCA projection of the pathway ranking from enrichment analysis. The predicted clusters correspond to the SWI/SNF / non-SWI/SNF division, allowing a perfect separation of KO lines. **B)** Enriched pathways in at least four SWI/SNF mutants or at least three non SWI/SNF mutants. Row clustering separated pathways differentially dysregulated in SWI/SNF mutants (in bold), pathways dysregulated in all cell lines, and those only dysregulated in non-SWI/SNF mutants. Pathway names are colored according to pathway categories. The + and – signs represent the NES score from GSEA indicating if the pathway is significantly upregulated (+), downregulated (-) or not significant (empty). **C)** Top dysregulated Reactome pathway categories in SWI/SNF mutants, considering all enriched pathway (upper line) or only the Top10 most enriched ones (lower line) for each cell line. Rankings correspond to the number of significantly enriched pathways for at least one SWI/SNF mutant in each category. **D)** Top dysregulated Reactome pathway categories in non-SWI/SNF mutants, considering all enriched pathway (upper line) or only the Top10 most enriched ones (lower line) for each cell line. Rankings correspond to the number of significantly enriched pathways for at least one SWI/SNF mutant in each category. **G)** Enriched pathways in transcriptome, as detected by GSEA and ROntoTools. The number of significant pathways flagged by both methods represented a small proportion of the total number of significant pathways highlighting a modest agreement between both enrichment methods. **H)** Top10 enriched pathways in the transcriptome of SWI/SNF mutants. Pathways are ranked following the category rankings of panel C. Inside each category, pathways are ranked based on the number of mutant cell lines for which the pathway appears in the Top10. The numbers inside the cells indicate the geometric mean of ranks of GSEA and ROntoTools, in white, if the pathway is in the Top10 pathways of the mutant, and in black, otherwise.

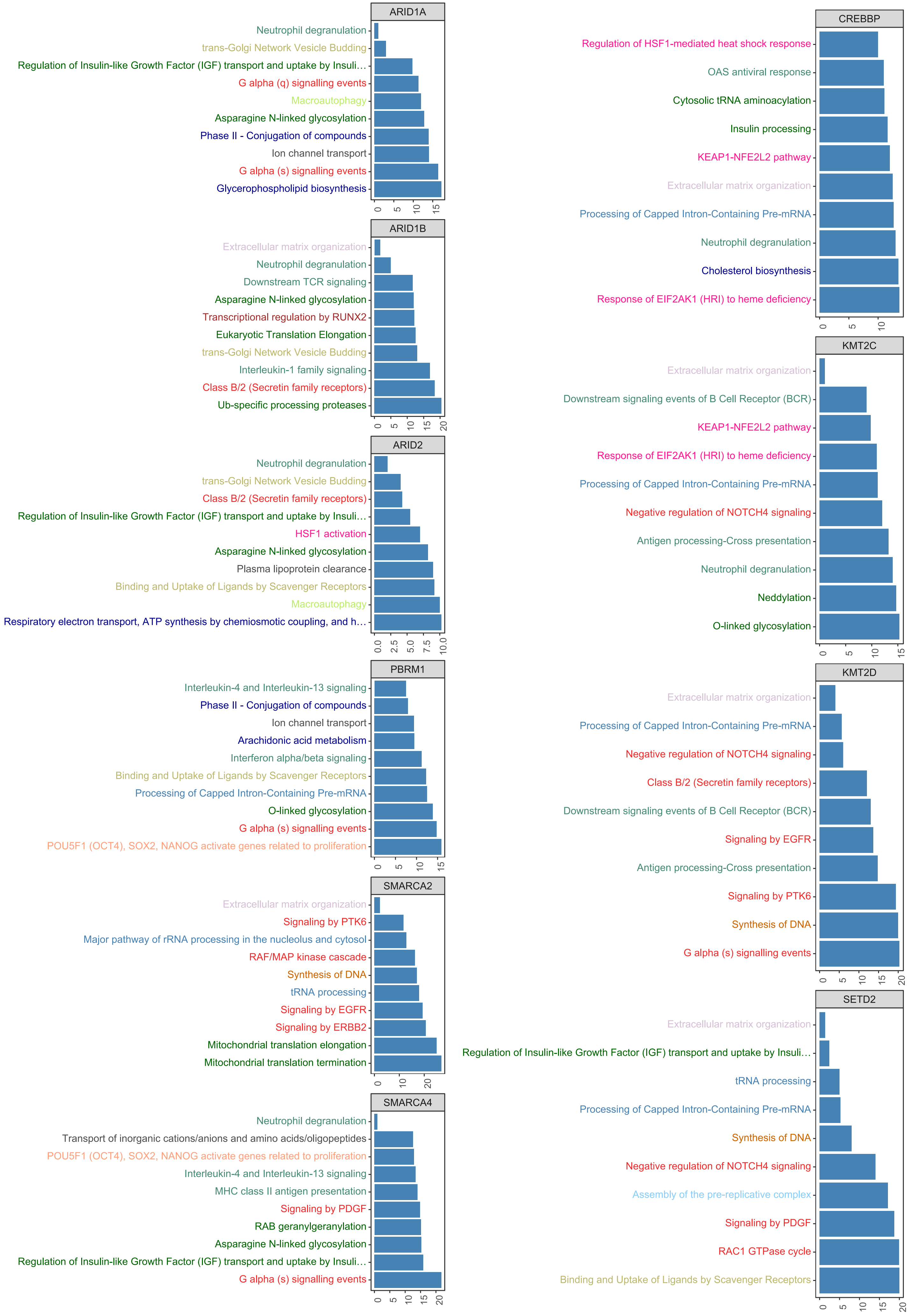
ARID1A
Neutrophil degranulation
trans-Golgi Network Vesicle Budding
Regulation of Insulin-like Growth Factor (IGF) transport and uptake by Insuli…
G alpha (q) signalling events
Macroautophagy
Asparagine N-linked glycosylation
Phase II - Conjugation of compounds
Ion channel transport
G alpha (s) signalling events
Glycerophospholipid biosynthesis
0 5 10 15
ARID1B
Extracellular matrix organization
Neutrophil degranulation
Downstream TCR signaling
Asparagine N-linked glycosylation
Transcriptional regulation by RUNX2
Eukaryotic Translation Elongation
trans-Golgi Network Vesicle Budding
Interleukin-1 family signaling
Class B/2 (Secretin family receptors)
Ub-specific processing proteases
0 5 10 15 20
ARID2
Neutrophil degranulation
trans-Golgi Network Vesicle Budding
Class B/2 (Secretin family receptors)
Regulation of Insulin-like Growth Factor (IGF) transport and uptake by Insuli…
HSF1 activation
Asparagine N-linked glycosylation
Plasma lipoprotein clearance
Binding and Uptake of Ligands by Scavenger Receptors
Macroautophagy
Respiratory electron transport, ATP synthesis by chemiosmotic coupling, and h…
0.0 2.5 5.0 7.5 10.0
PBRM1
Interleukin-4 and Interleukin-13 signaling
Phase II - Conjugation of compounds
Ion channel transport
Arachidonic acid metabolism
Interferon alpha/beta signaling
Binding and Uptake of Ligands by Scavenger Receptors
Processing of Capped Intron-Containing Pre-mRNA
O-linked glycosylation
G alpha (s) signalling events
POU5F1 (OCT4), SOX2, NANOG activate genes related to proliferation
0 5 10 15
SMARCA2
Extracellular matrix organization
Signaling by PTK6
Major pathway of rRNA processing in the nucleolus and cytosol
RAF/MAP kinase cascade
Synthesis of DNA
tRNA processing
Signaling by EGFR
Signaling by ERBB2
Mitochondrial translation elongation
Mitochondrial translation termination
0 10 20
SMARCA4
Neutrophil degranulation
Transport of inorganic cations/anions and amino acids/oligopeptides
POU5F1 (OCT4), SOX2, NANOG activate genes related to proliferation
Interleukin-4 and Interleukin-13 signaling
MHC class II antigen presentation
Signaling by PDGF
RAB geranylgeranylation
Asparagine N-linked glycosylation
Regulation of Insulin-like Growth Factor (IGF) transport and uptake by Insuli…
G alpha (s) signalling events
0 5 10 15 20
CREBBP
Regulation of HSF1-mediated heat shock response
OAS antiviral response
Cytosolic tRNA aminoacylation
Insulin processing
KEAP1-NFE2L2 pathway
Extracellular matrix organization
Processing of Capped Intron-Containing Pre-mRNA
Neutrophil degranulation
Cholesterol biosynthesis
Response of EIF2AK1 (HRI) to heme deficiency
0 5 10
KMT2C
Extracellular matrix organization
Downstream signaling events of B Cell Receptor (BCR)
KEAP1-NFE2L2 pathway
Response of EIF2AK1 (HRI) to heme deficiency
Processing of Capped Intron-Containing Pre-mRNA
Negative regulation of NOTCH4 signaling
Antigen processing-Cross presentation
Neutrophil degranulation
Neddylation
O-linked glycosylation
0 5 10 15
KMT2D
Extracellular matrix organization
Processing of Capped Intron-Containing Pre-mRNA
Negative regulation of NOTCH4 signaling
Class B/2 (Secretin family receptors)
Downstream signaling events of B Cell Receptor (BCR)
Signaling by EGFR
Antigen processing-Cross presentation
Signaling by PTK6
Synthesis of DNA
G alpha (s) signalling events
0 5 10 15 20
SETD2
Extracellular matrix organization
Regulation of Insulin-like Growth Factor (IGF) transport and uptake by Insuli…
tRNA processing
Processing of Capped Intron-Containing Pre-mRNA
Synthesis of DNA
Negative regulation of NOTCH4 signaling
Assembly of the pre-replicative complex
Signaling by PDGF
RAC1 GTPase cycle
Binding and Uptake of Ligands by Scavenger Receptors
0 5 10 15 20

**Supplementary Fig. 6.** **Top10 entiched pathways in the transcriptome of SWI/SNF (first column) and non-SWI/SNF (second column) mutants**. Pathways are colored based on the Reactome category they belong to (see Fig. 3B for legend).

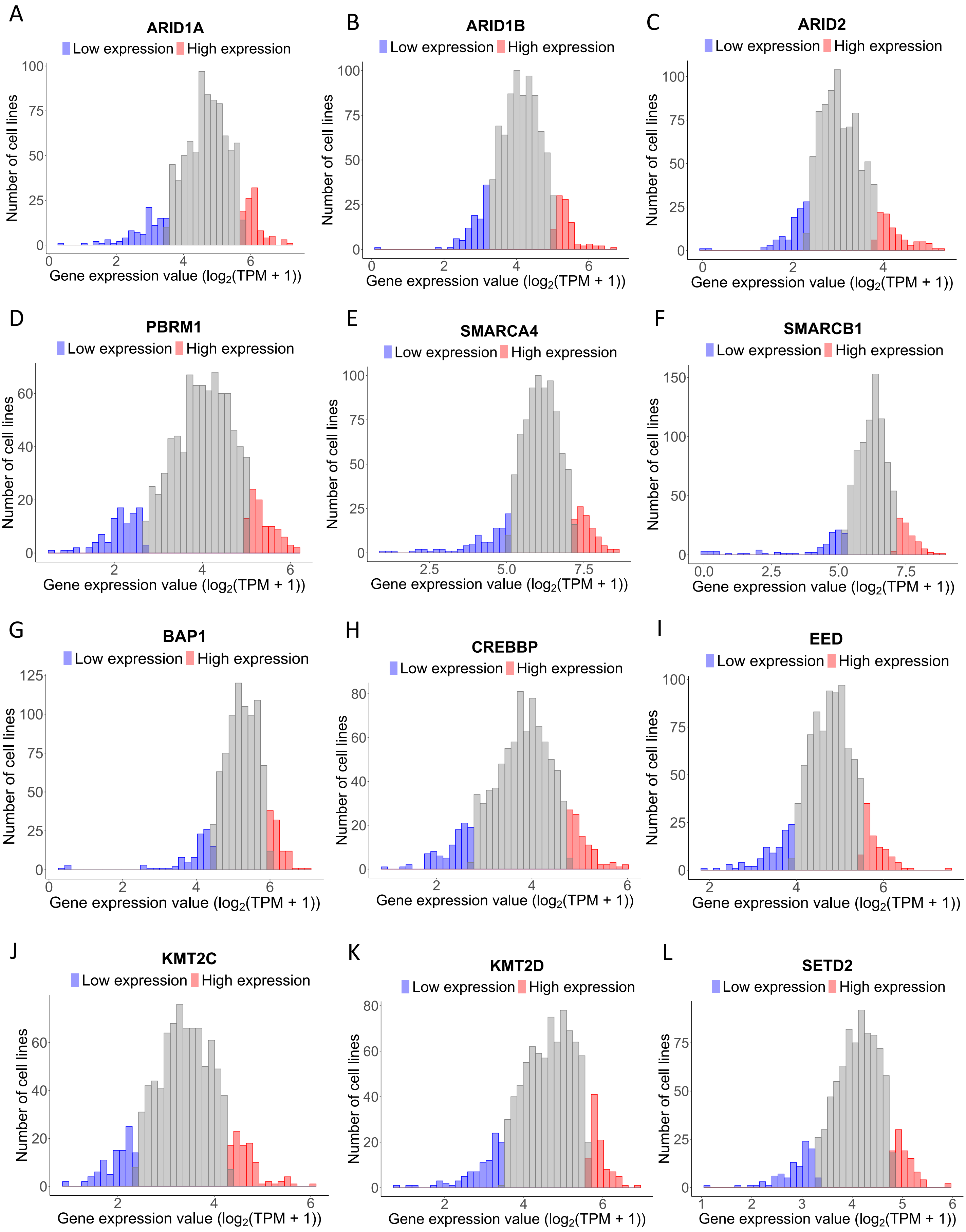


**Supplementary Fig 7. Distribution of mutant-like chromatin remodeling genes in the Depmap cancer cell line transcriptomics dataset.** Cell lines in the highest and lowest 10% quantiles are selected to form the high (red) and low (blue) expression groups, respectively.

### ARID1A

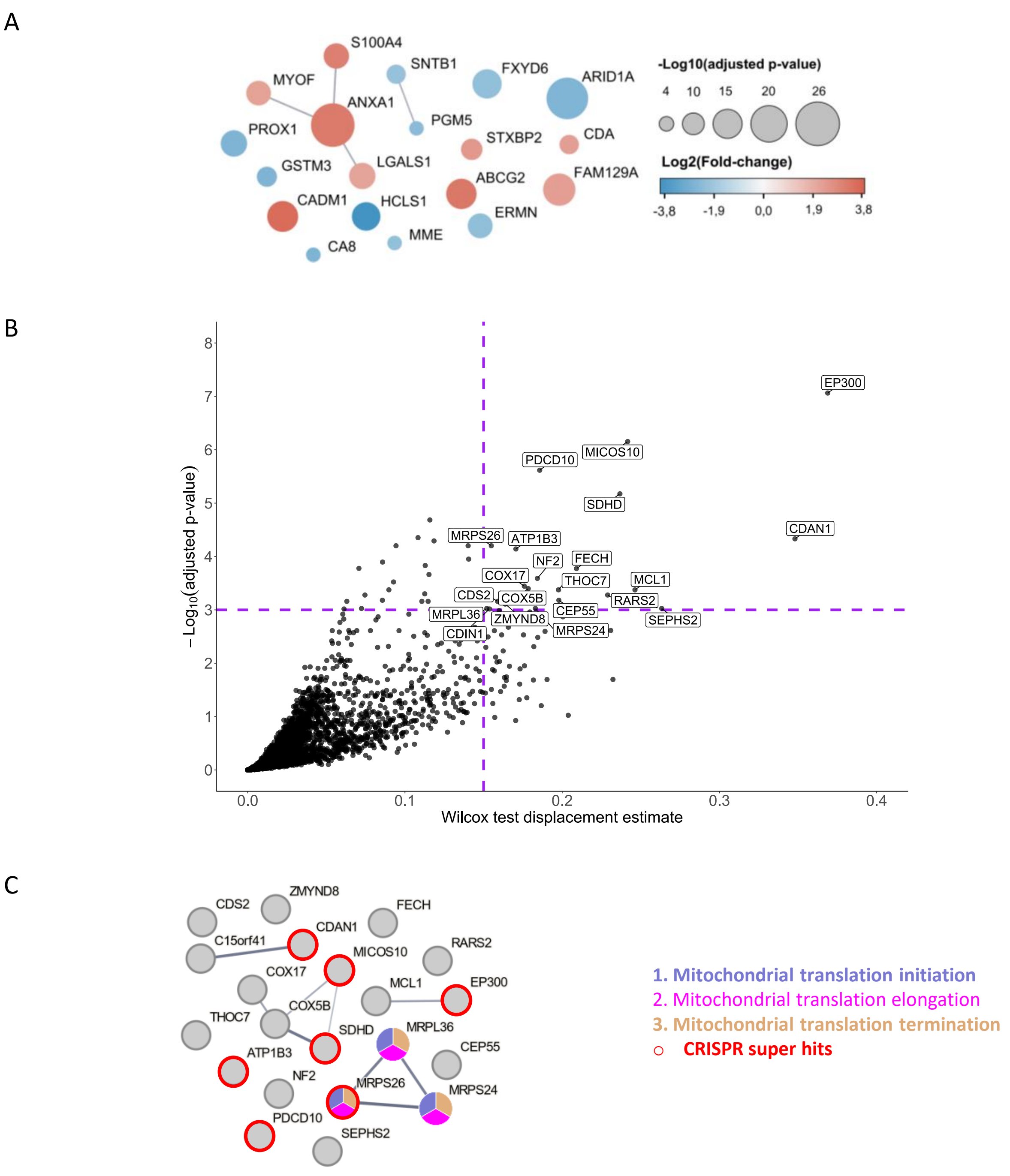


**Supplementary Fig. 8. <u>Differentially expressed proteins and predicted synthetic lethal targets in ARID1A-KO or low expression cell lines</u>. A)** STRING maps of proteins with $|\log_2(FC)| > 2$ and adjusted p-value < 0.05. Every node represents a protein. The node color represents the $\log_2(FC)$ and the node size is proportional to the significance of each protein expressed as -log10(p-adj). Edges represent functional interactions as predicted by STRING, which are inferred from text mining, databases, experiments, co-expression, neighborhoods, gene fusions and co-occurrence using a custom threshold of 0.5. The linewidth indicates STRING interaction confidence (from 0 to 1). **B)** Volcano plot showing synthetic lethality hits for the mutant-like gene (here ARID1A) from the Depmap analysis. The x-axis shows displacement estimate of the Wilcox test indicating the difference in the CRISPR score distributions between high and low expression groups for the mutant-like gene (here ARID1A). The dotted lines indicate the threshold for defining hits (p-adj <0.001 and Wilcox test displacement > 0.15). **C)** STRING map of hit genes as inferred from the Depmap analysis. The STRING parameters are similar to panel A. The red circles represent super hits (hits with p-adj < 1e-4). The node color code represents the pathway affiliation for CRISPR genes.

### ARID1B

A

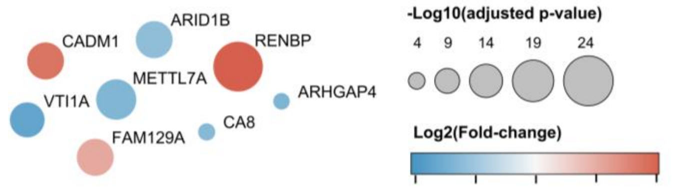


B

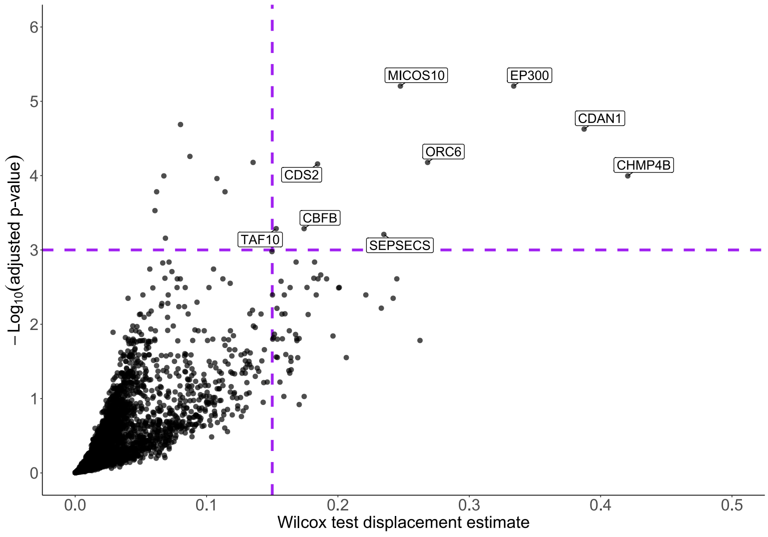


C

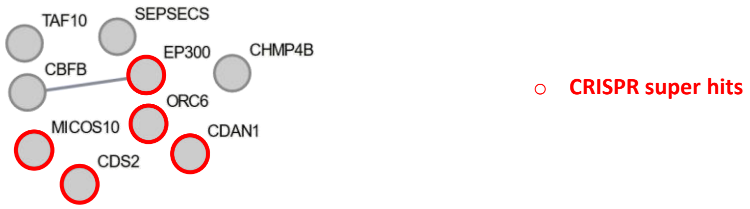


**Supplementary Fig. 9. <u>Differentially expressed proteins and predicted synthetic lethal targets in ARID1B_KO or low expression cell lines</u>.** See Supplementary Figure 7 for legend.

### ARID2

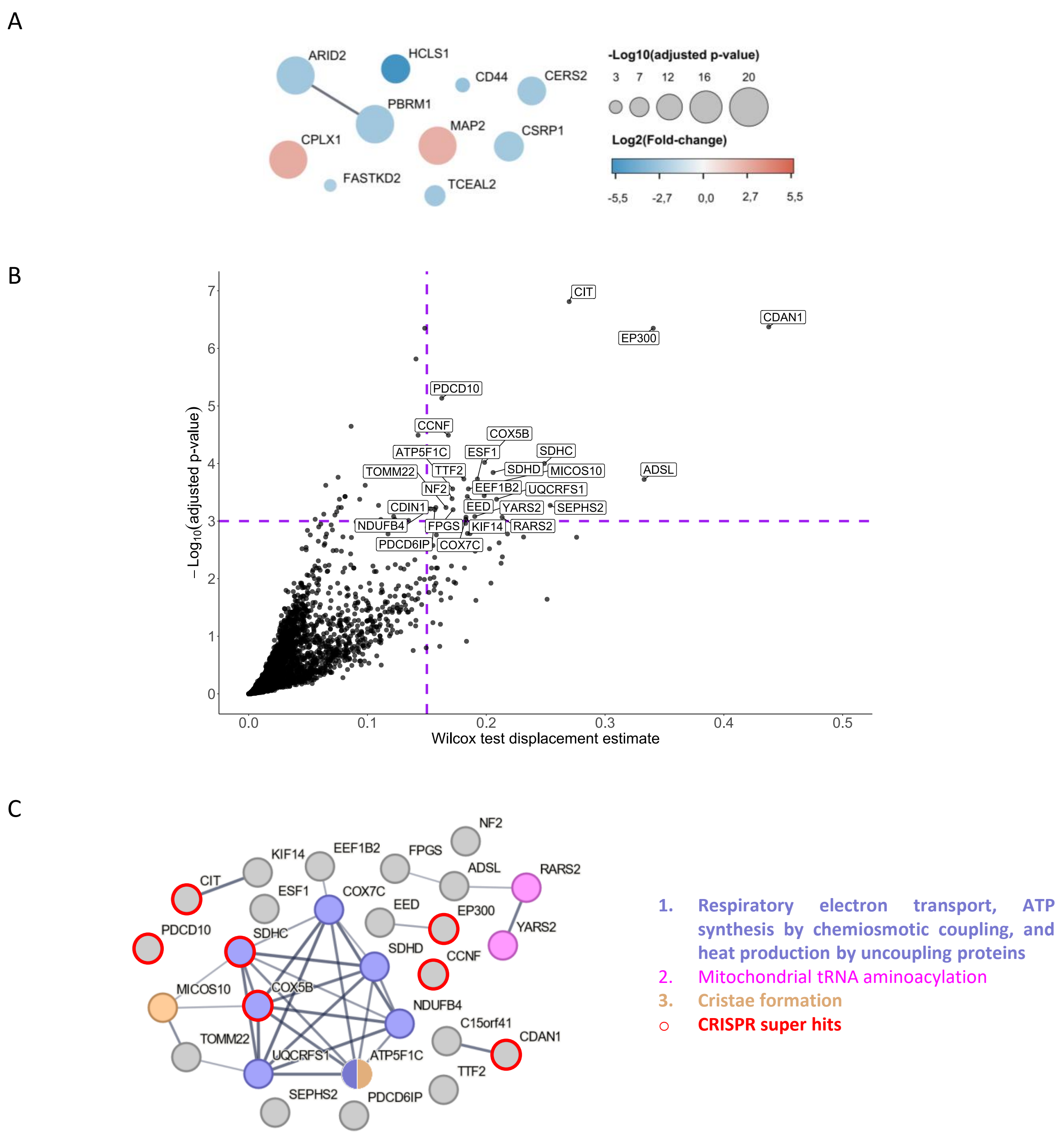


**Supplementary Fig. 10. <u>Differentially expressed proteins and predicted synthetic lethal targets in ARID2_KO or low expression cell lines</u>** See Supplementary Figure 7 for legend.

### PBRM1

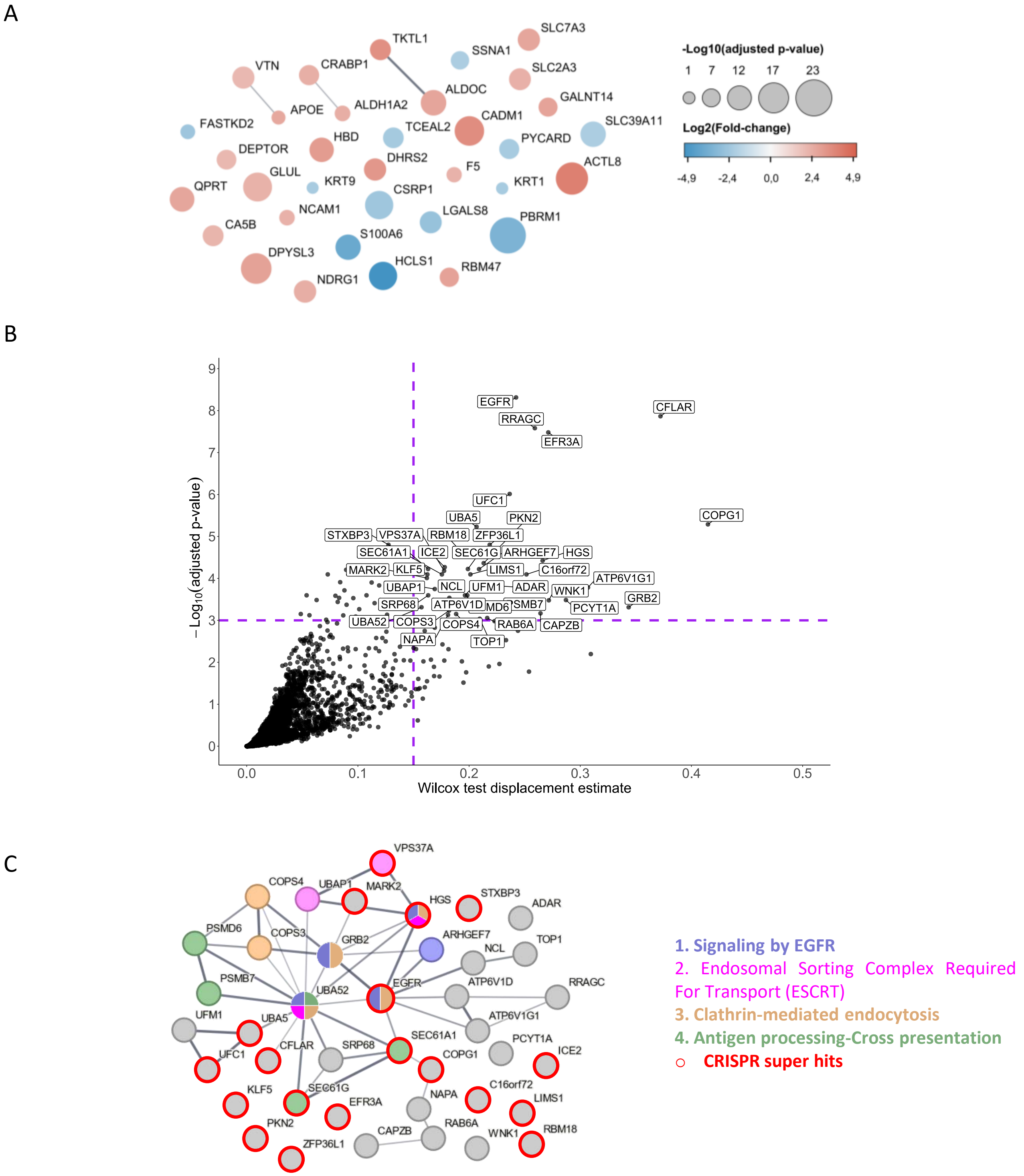


**Supplementary Fig. 11. <u>Differentially expressed proteins and predicted synthetic lethal targets in PBRM1-KO or low expression cell lines</u>** See Supplementary Figure 7 for legend.

### SMARCA2

A

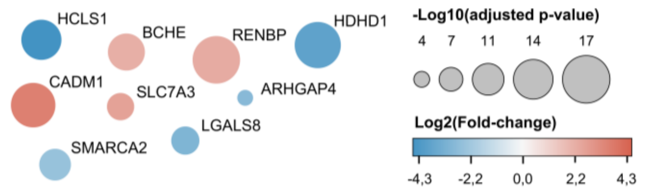


B

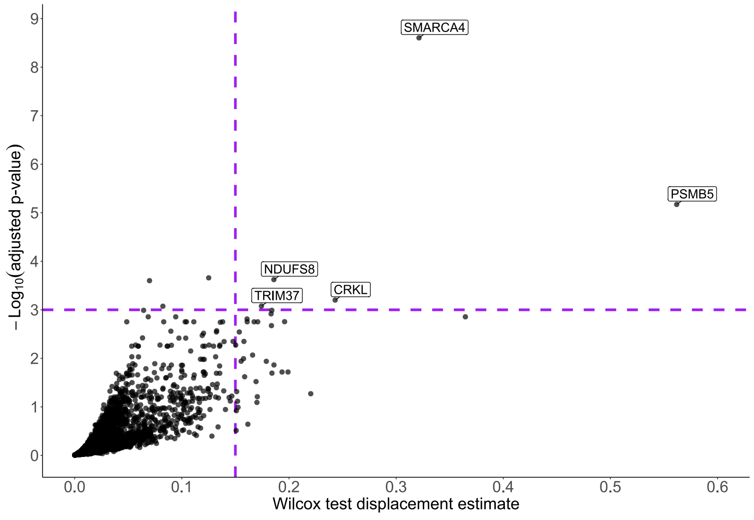


C

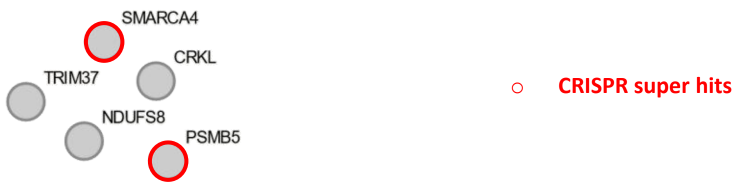


**Supplementary Fig. 12. <u>Differentially expressed proteins and predicted synthetic lethal targets in SMARCA2-KO or low expression cell lines.</u>** See Supplementary Figure 7 for legend.

### SMARCA4

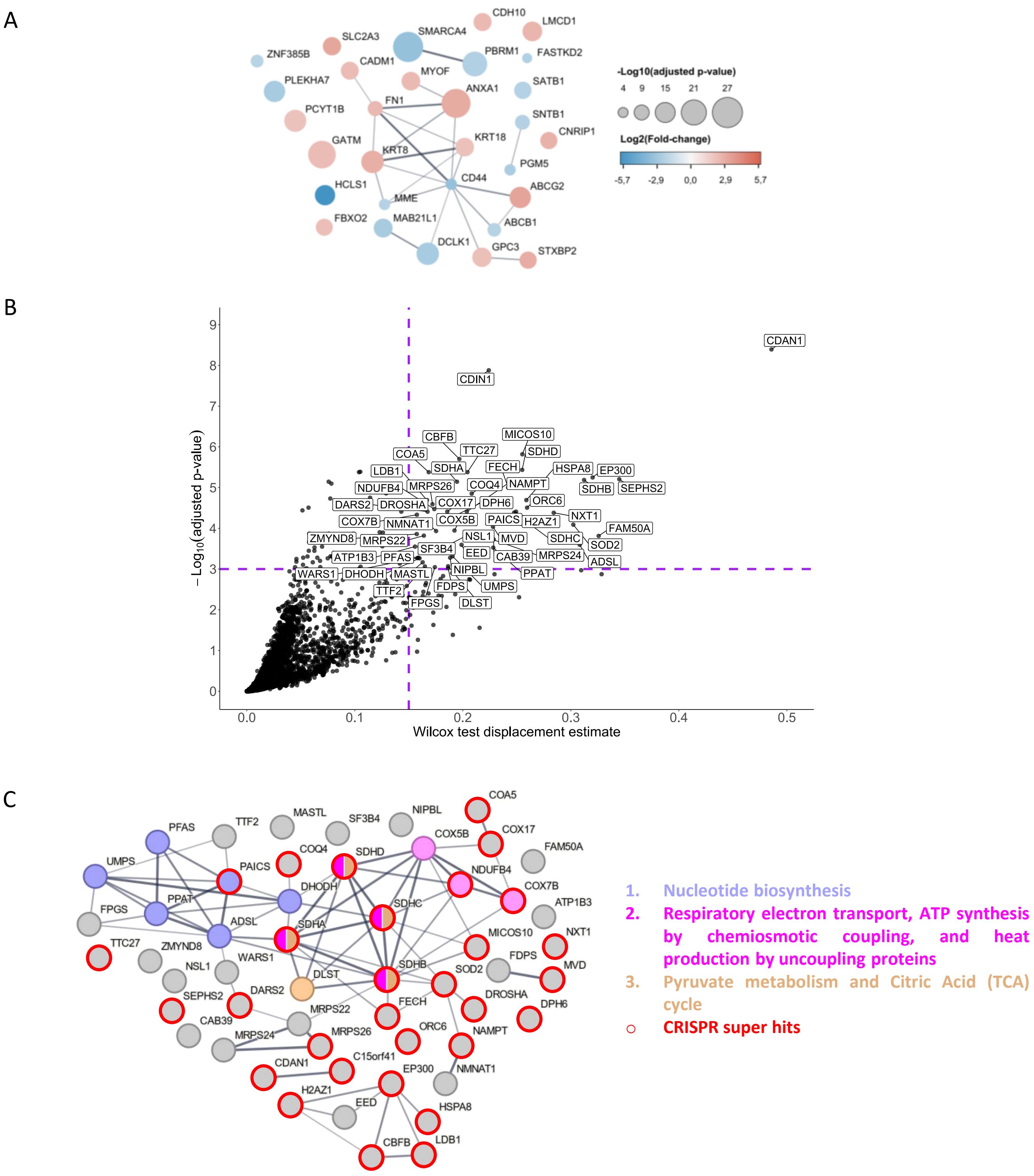


**Supplementary Fig. 13. Differentially expressed proteins and predicted synthetic lethal targets in SMARCA4-KO or low expression cell lines.** See Supplementary Figure 7 for legend.

SMARCB1

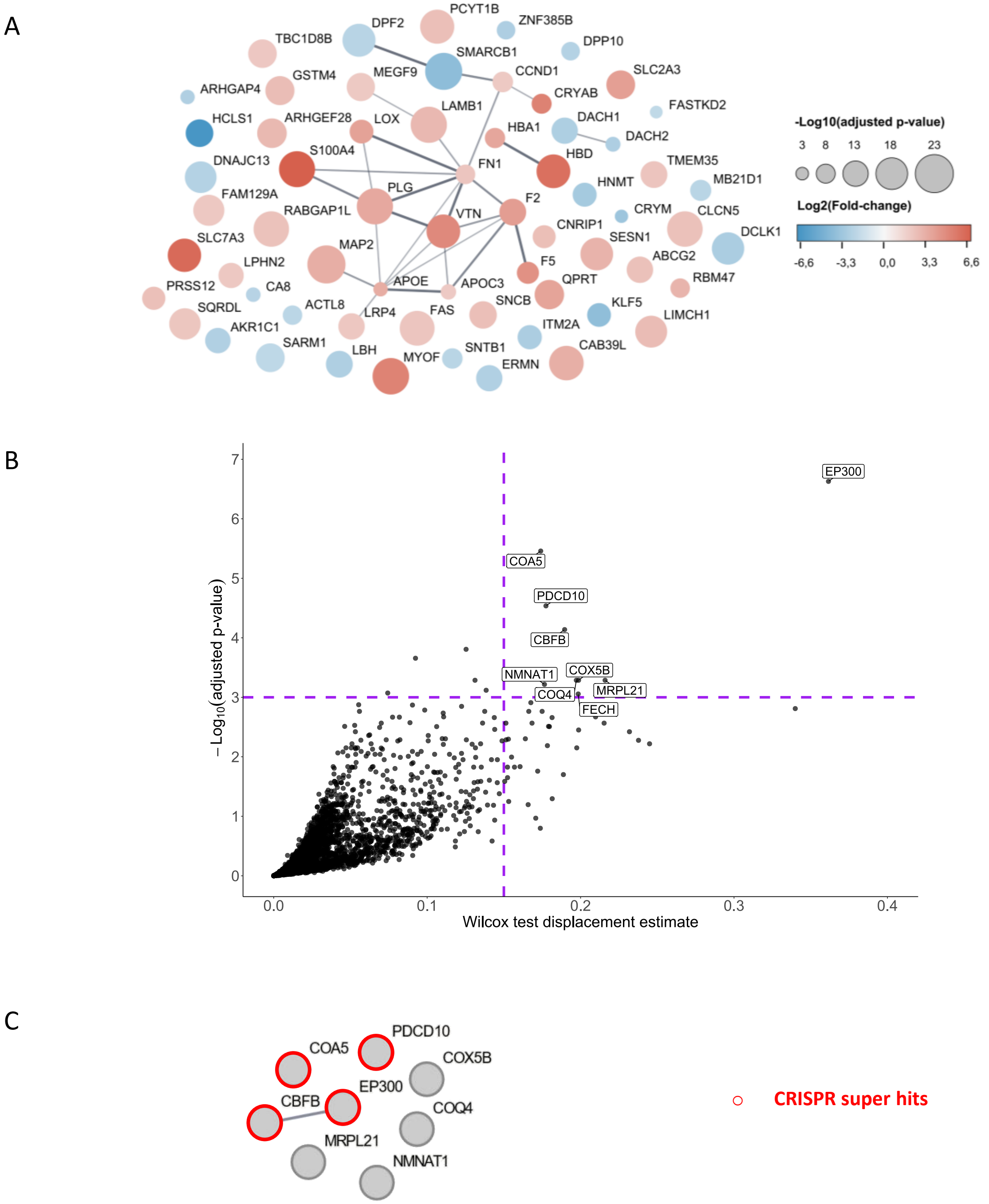


**Supplementary Fig. 14. Differentially expressed proteins and predicted synthetic lethal targets in SMARCB1-KO or low expression cell lines.** See Supplementary Figure 7 for legend.

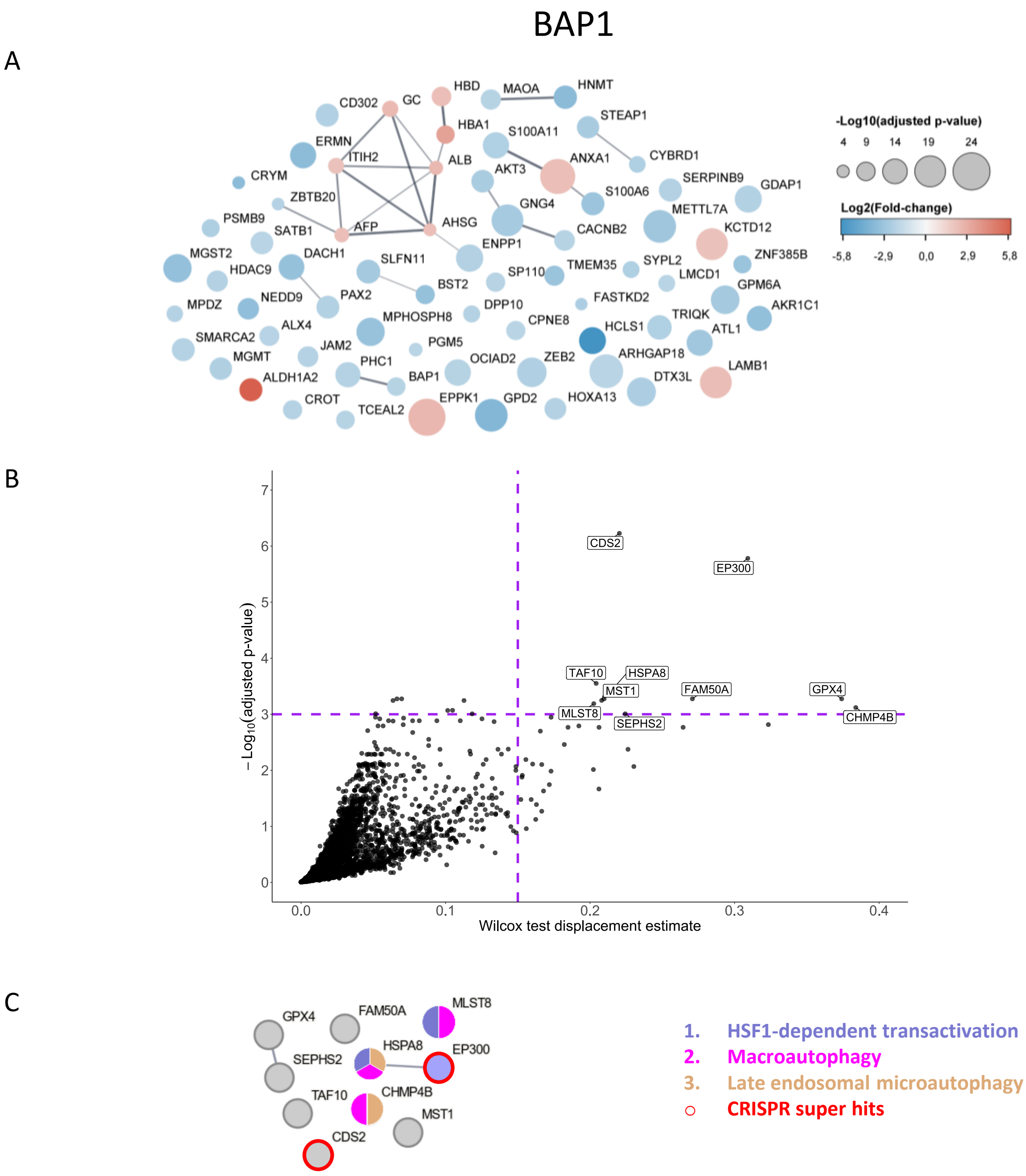


**Supplementary Fig. 15. <u>Differentially expressed proteins and predicted synthetic lethal targets in BAP1-KO or low expression cell lines.</u>** See Supplementary Figure 7 for legend.

### CREBBP

A

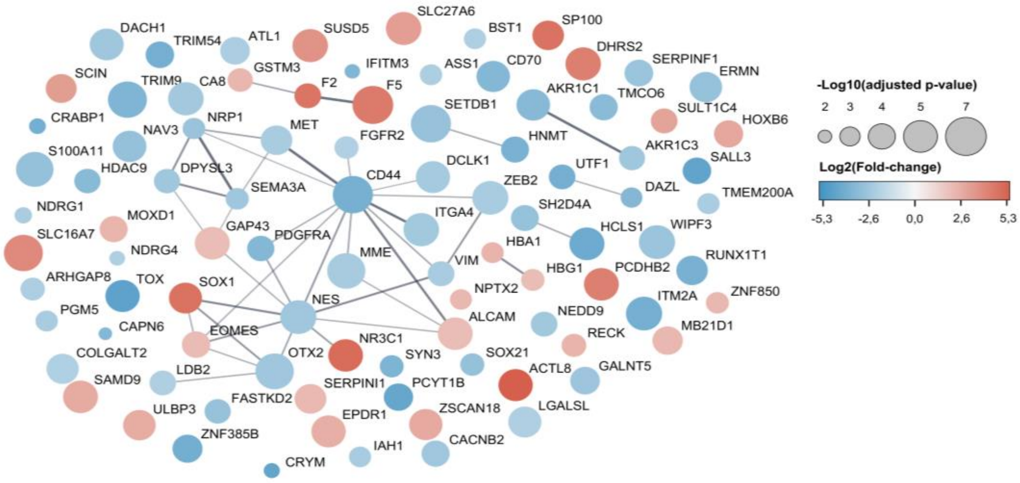


B

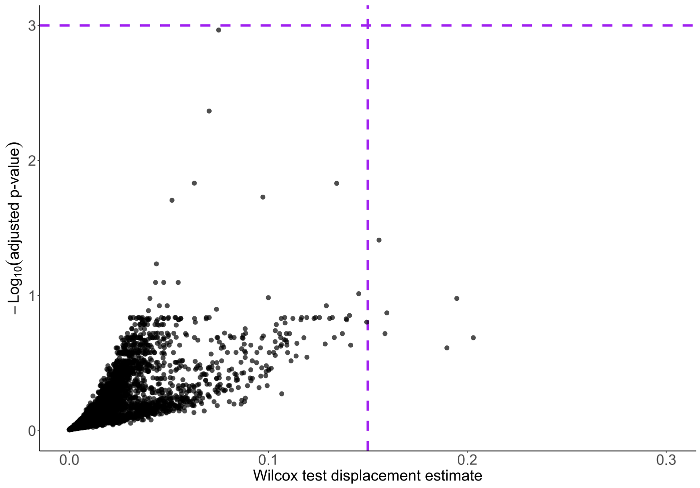


**Supplementary Fig. 16.** **<u>Differentially expressed proteins and predicted synthetic lethal targets in CREBBP-KO or low expression cell lines.</u>** See Supplementary Figure 7 for legend.

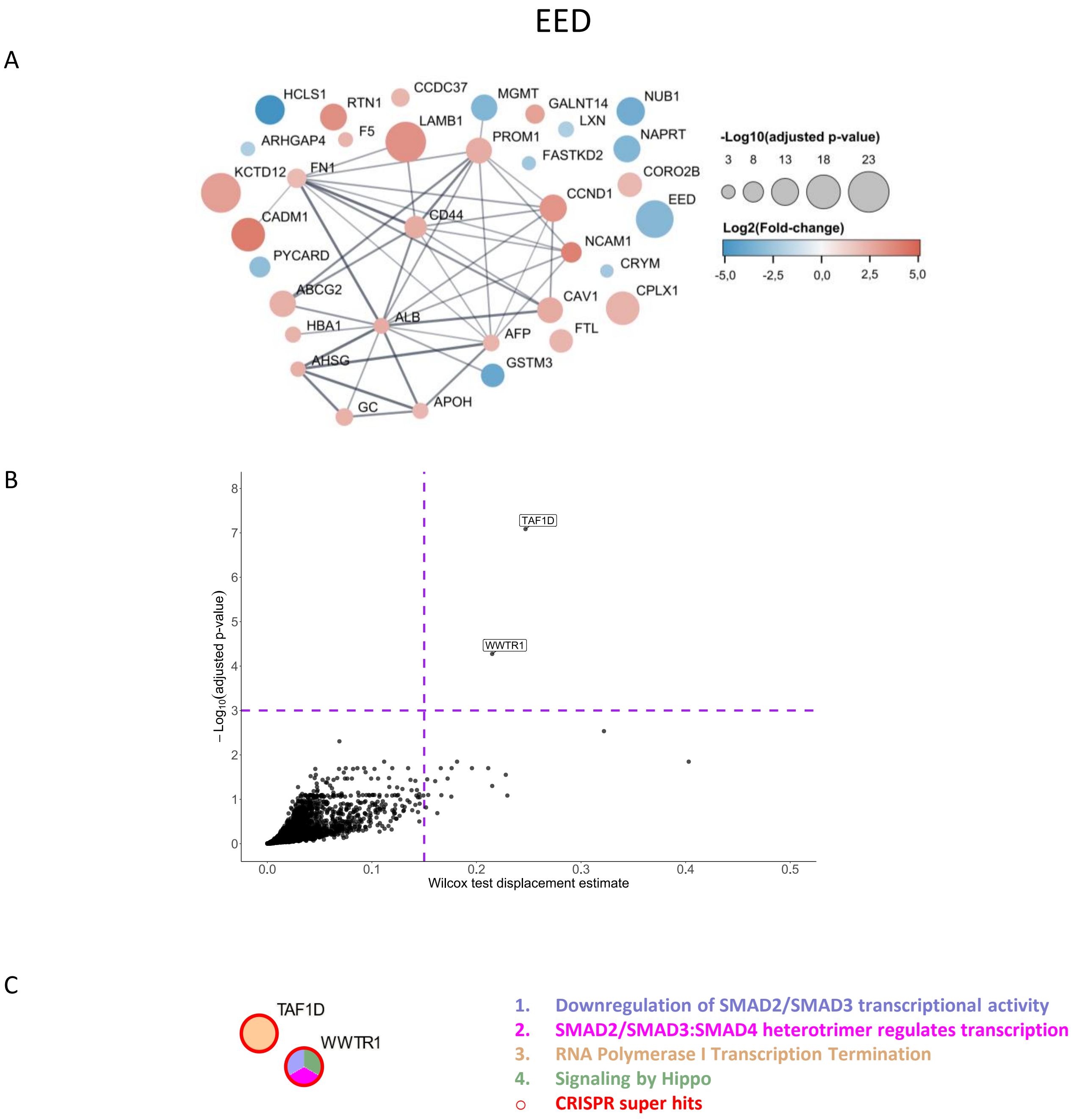


**Supplementary Fig. 17. <u>Differentially expressed proteins and predicted synthetic lethal targets in EED-KO or low expression cell lines.</u>** See Supplementary Figure 7 for legend.

### KMT2C

A

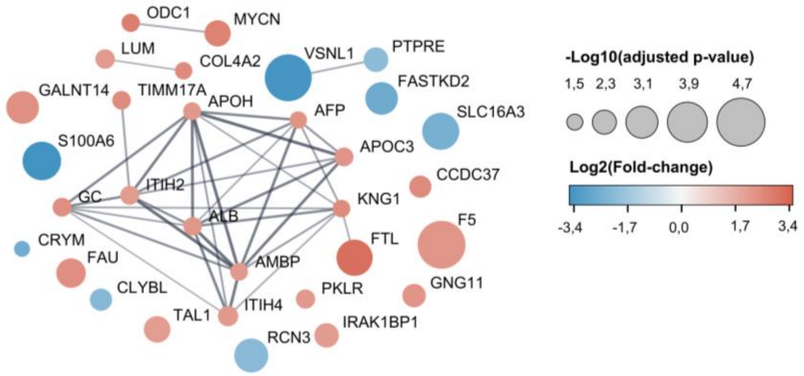


B

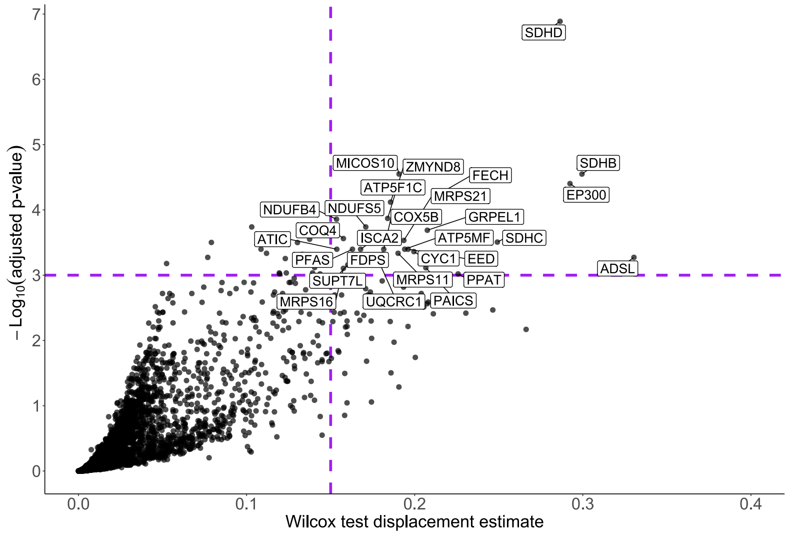


C

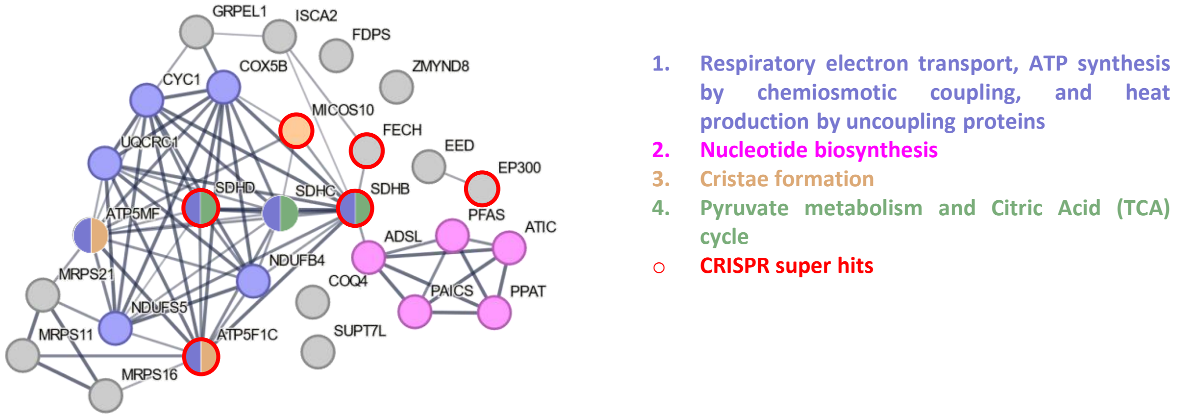


**Supplementary Fig. 18.** **<u>Differentially expressed proteins and predicted synthetic lethal targets in KMT2C-KO or low expression cell lines.</u>** See Supplementary Figure 7 for legend.

### KMT2D

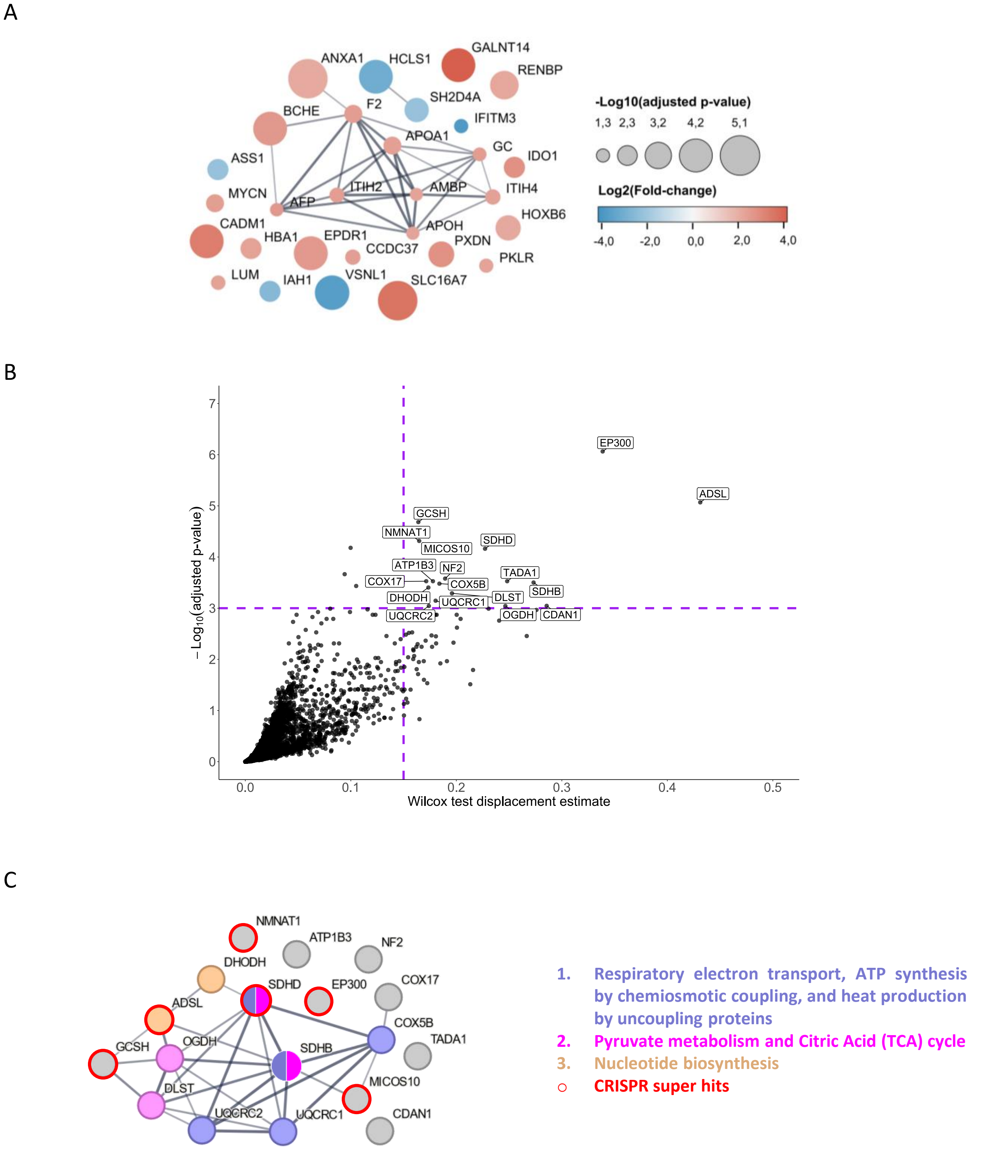


**Supplementary Fig. 19.** **<u>Differentially expressed proteins and predicted synthetic lethal targets in KMT2D-KO or low expression cell lines.</u>** See Supplementary Figure 7 for legend.

### SETD2

A

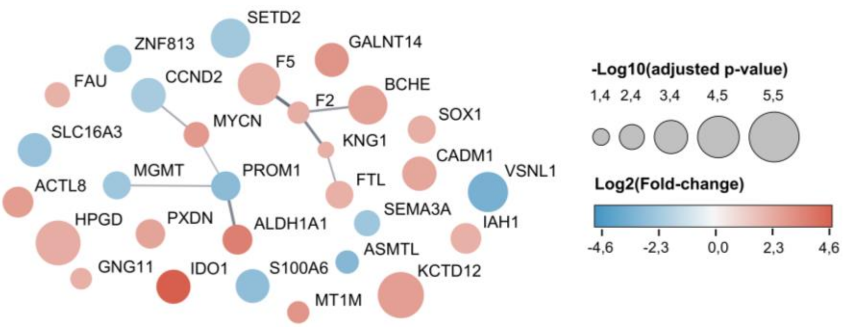


B

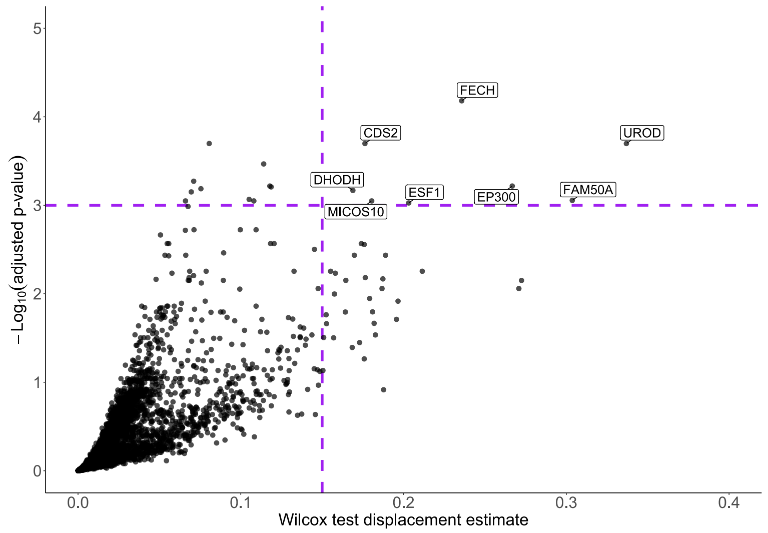


C

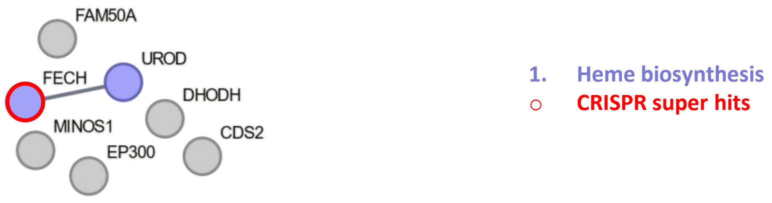


**Supplementary Fig. 20. <u>Differentially expressed proteins and predicted synthetic lethal targets in SETD2-KO or low expression cell lines.</u>** See Supplementary Figure 7 for legend.

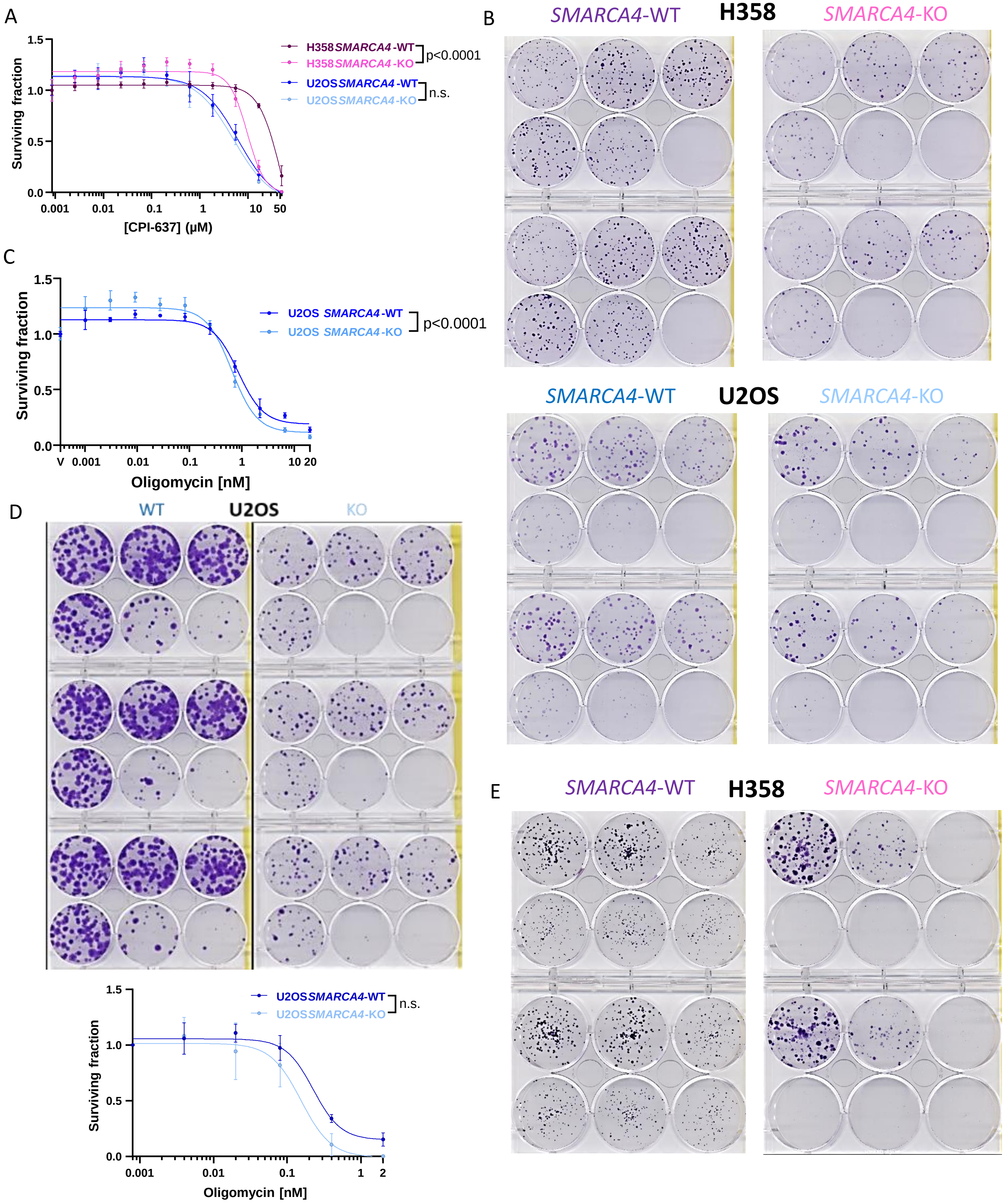


**Supplementary Fig. 21. Revalidation experiments of cytoxicity of inhibitors of EP300 and of mitochondrial respiration in SMARCA4 isogenic cell lines.** Datapoints are mean ± SD, with three technical replicates per point. **A)** Short-term cytotoxicity of EP300 inhibitor CPI-637 in H358 and U2OS cell lines. **B)** Long-term colony forming assays in *SMARCA4*-isogenic models exposed to increasing concentrations of CPI-637 for 10 days (U2OS cells) or 13 days (H358 cells). Replicate experiments of Fig. 7C. **C)** Short-term cytotoxicity of oligomycin in U2OS cell lines. **D)** Long-term colony forming assays in U2OS cell lines after exposure to oligomycin over 10 days. Two-way ANOVA testing WT/KO conditions and dose effects: p=0,0112. **E)** Long-term colony forming assays in H358 isogenic models exposed to increasing concentrations of oligomycin over 17 days. Replicate experiments of Fig. 7F.
